# A novel lipid-triggered allosteric site modulates LC3-LIR receptor binding activity

**DOI:** 10.64898/2026.03.10.710726

**Authors:** Deepanshi Gahlot, Jesu Castin, Shruti Mathur, Debajyoti Das, Ajit Kumar, Akanksha Arun, Chandrima Gain, Mridul Sharma, Ravi Kant Pal, Niyati Jain, Bichitra K. Biswal, Rajat Singh, Lipi Thukral

## Abstract

Membrane recruitment is a fundamental regulator of protein function. However, the allosteric mechanisms by which lipid binding controls protein activity remain poorly understood. In autophagy, the ubiquitin-like protein LC3 is lipid-anchored to autophagosomes, where it is essential for receptor recruitment and vesicle formation. While LC3-receptor interactions are structurally well defined, how membrane engagement governs LC3 functional dynamics has remained enigmatic. Here, we uncover that membrane binding triggers a major conformational transition in LC3, exposing functional pockets that are occluded in its cytosolic form. We demonstrate that this shift is mediated by dynamic coupling between the allosteric site (α3– loop5–β3–loop6) and the functional binding pockets. To conclusively test this mechanism, we utilised an ensemble-based protein design strategy guided by molecular dynamics to engineer the allosteric site. From a series of mutants, two variants emerged that stabilized LC3 conformation in either active or inactive state on the membrane. X-ray crystal structures of mutant LC3, biophysical assays, super-resolution microscopy, and TEM confirmed that the activated allosteric site mutant facilitates receptor binding and cargo capture. In contrast, the inactive variant is functionally inert on the membrane. Our work identifies a fundamental lipid-triggered allosteric site in LC3 that is critical for autophagy regulation and broader implication of membrane-dependent reprogrammable protein activities.

## Introduction

Allostery refers to the long-range communication between two spatially distinct sites within a protein, where binding or perturbation at one site induces a conformational change at another, thereby modulating activity^1^. Such mechanisms act like molecular relay systems, transmitting signals through the protein to regulate distant functional sites^2,3^. This principle underlies diverse biological processes, from fine-tuning enzyme catalysis^4^ and mediating protein-protein interactions to coordinating large assemblies such as transcriptional machinery and ribosomes^5,6^. Because of this versatility, allosteric sites have been widely exploited in drug discovery to achieve precise modulation of protein activity with greater specificity than traditional active site targeting^7^. Lipid-binding events are also known to directly or indirectly influence protein function^8,9^. Membranes, therefore, are not merely passive scaffolds for protein localization but active participants in shaping protein conformations and activities^10,11^. Despite its conceptual importance, the molecular basis of lipid-driven allostery remains poorly understood, largely because membrane-protein interactions are transient, heterogeneous, and challenging to capture structurally.

Autophagy is a powerful model pathway to study how membrane interactions regulate protein activity. It is a highly dynamic cellular quality control process in which isolation membranes are remodeled into double-membraned autophagosome vesicles^12^. The formation of autophagosome is driven by the coordinated action of numerous protein complexes on lipid membranes. These complexes regulate each step, from phagophore initiation and membrane expansion to cargo selection and lysosomal fusion for degradation^13,14^. Several autophagy proteins are recruited to the autophagosomal membrane during the formation of the vesicle^15^. Among these, LC3B (microtubule-associated protein 1A/1B-light chain 3 B), hereafter referred to as LC3, is the mammalian ortholog of yeast ATG8 and has been extensively studied for its structural and functional roles in autophagy. It plays a central role in autophagosome maturation by anchoring to lipid membranes via phosphatidylethanolamine (PE)^16,17^. In its lipidated form, termed LC3-II, it is anchored to autophagosomal membranes and represents the functionally active protein. LC3 consists of two conserved hydrophobic pockets (HP1 and HP2), which mediate interactions with short linear LC3-interacting regions (LIRs) motifs present in cargo receptors^18,19^, a property that is crucial for selective autophagy^20^. The LIR motif (W/Y/F-x-x-L/I/V) inserts into pockets and enables LC3 to recruit autophagy receptors, including p62, OPTN, and NBR1, amongst others^21–23^. Thus, the membrane association of LC3 is biologically significant, enabling interactions with over 160 autophagy receptors in human that mediate selective cargo recognition, sequestration, and ultimately cargo degradation in lysosomes^24,25^. While LC3 is widely used as a molecular switch and marker of autophagy^26^, it remains unclear how lipid association alters LC3 structure and dynamics to regulate receptor binding on the membrane.

We hypothesized that lipids may function as dynamic regulators and may modulate the activity of LC3 functional interfaces. Given that high-resolution structures were available for the cytosolic LC3 and as complex with LIR peptides, we employed physics-based molecular dynamics simulations to study its membrane association and how lipid interactions modulate its activity. We found that LC3 adopted a distinct structural conformation when associated with membranes. The membrane binding favored an open conformation of the hydrophobic pockets, while in the soluble state, they remained closed, suggesting allosteric coupling between the membrane interface and the binding sites. Robust long-range communication analysis further uncovered a network of interactions converging on a novel allosteric site that engages with lipids. Guided by these computational insights, we engineered a set of mutants in which the allosteric site was mutated to stabilize conformations resembling either the active or inactive binding states. Two final mutants of the allosteric site modulated receptor activity, which we validated through a series of experiments including biophysical binding assays, transmission electron microscopy (TEM), co-immunoprecipitation, and flux analyses. X-ray crystallography of the active mutant revealed rearrangements in both the receptor-binding pocket and a distal allosteric site, further validating the predictions from simulations. The ability to conformationally mimic diverse protein states through MD simulations provides a powerful platform to deepen our understanding of protein function, particularly at membranes where experimental studies remain highly challenging.

## Results

### LC3 undergoes lipid-triggered conformational remodeling on the autophagic membranes

LC3 transitions from a water-favored environment to a membrane-anchored conformation via lipidation **(Fig. 1a)**. While lipidation is well-established as a molecular switch that anchors LC3 to membranes, the subsequent conformational changes to the protein itself remain unclear. To study this molecular switch and to understand how functional changes associated with membrane binding occur, simulations were performed on the lipidated, membrane bound form of LC3 (LC3^M+^). The experimentally resolved structure was used, and a lipid tail was attached as a membrane anchor, as described earlier^27^. To capture physiological complexity of autophagosomal membrane lipids, we used a mixed-lipid bilayer model representative of ER-like membranes^28^. In addition, the unmodified protein without conjugated lipid anchor representing the cytosolic state^29^ of LC3 was also simulated, allowing direct comparison with membrane-dependent dynamics (LC3^M-^) **(Fig. 1b)**. Three independent simulations capturing µs-scale dynamics were performed for both states and showed convergence across replicates (**Supplementary Fig. 1**). To dissect how LC3 reorganizes upon membrane association, we classified protein into three structurally distinct regions: (i) the membrane interacting site (MIS)^27^, (ii) the canonical LIR-hydrophobic binding pockets (BP)^20,30^, and (iii) the remaining segments of the protein (**Fig. 1c; Supplementary Fig. 2**). To quantify how protein reorganizes upon membrane association, we first mapped its intra-protein contacts and defined a contact as pair of residues within 5 Å for more than 50% of the simulation time across three independent trajectories (**Supplementary Table 1**). This measure provides a structural understanding of how tightly different regions of the protein pack together. Remarkably, in LC3^M+^ state, each region exhibited unique patterns of conformational reorganization and spatial reorientation upon local contact with the membrane (**Fig. 1d**). Within MIS, where protein and lipids are in direct contact, the internal interactions are reduced by more than half compared to LC3^M-^ state. As a result, the membrane binding notably reduced protein compactness, particularly within MIS (**Supplementary Fig. 3**). Simultaneously, these lost interactions were effectively compensated by robust protein-lipid contacts in LC3^M+^ **(Fig. 1e)**. In the MIS, conserved salt bridges (E4–K39, R10–E36, D48–R70) that stabilize the LC3^M-^ conformation (4.7 ± 2.3 Å) are disrupted upon membrane binding in LC3^M+^ state, with distances extending to 7.1 ± 2.6 Å (**Supplementary Fig. 4**). The key basic residues (K5, K8, R10, R37, K39, K42) remain tightly associated with the membrane (<5 Å), where they adopt characteristic snorkeling orientations. Surprisingly, in the binding pockets that are highly hydrophobic, intraprotein contacts were rearranged, concomitant with increased accessibility from 437.17 ± 40.4 Å^2^ to 502.83 ± 64.6 Å^2^ in LC3^M-^ and LC3^M+^, respectively (**Supplementary Fig. 5**). In the context of driving forces, interactions within the LC3^M-^ state are dominated by electrostatic contacts within the MIS and hydrophobic contacts within the binding pocket. In contrast, the rest of the protein showed greater cohesion, reflected by increase in intra-protein contacts, with 57% of the newly formed membrane-bound-specific hydrophobic interactions localized in this region (**Supplementary Fig. 6, Supplementary Table 1**).

**Figure 1:**
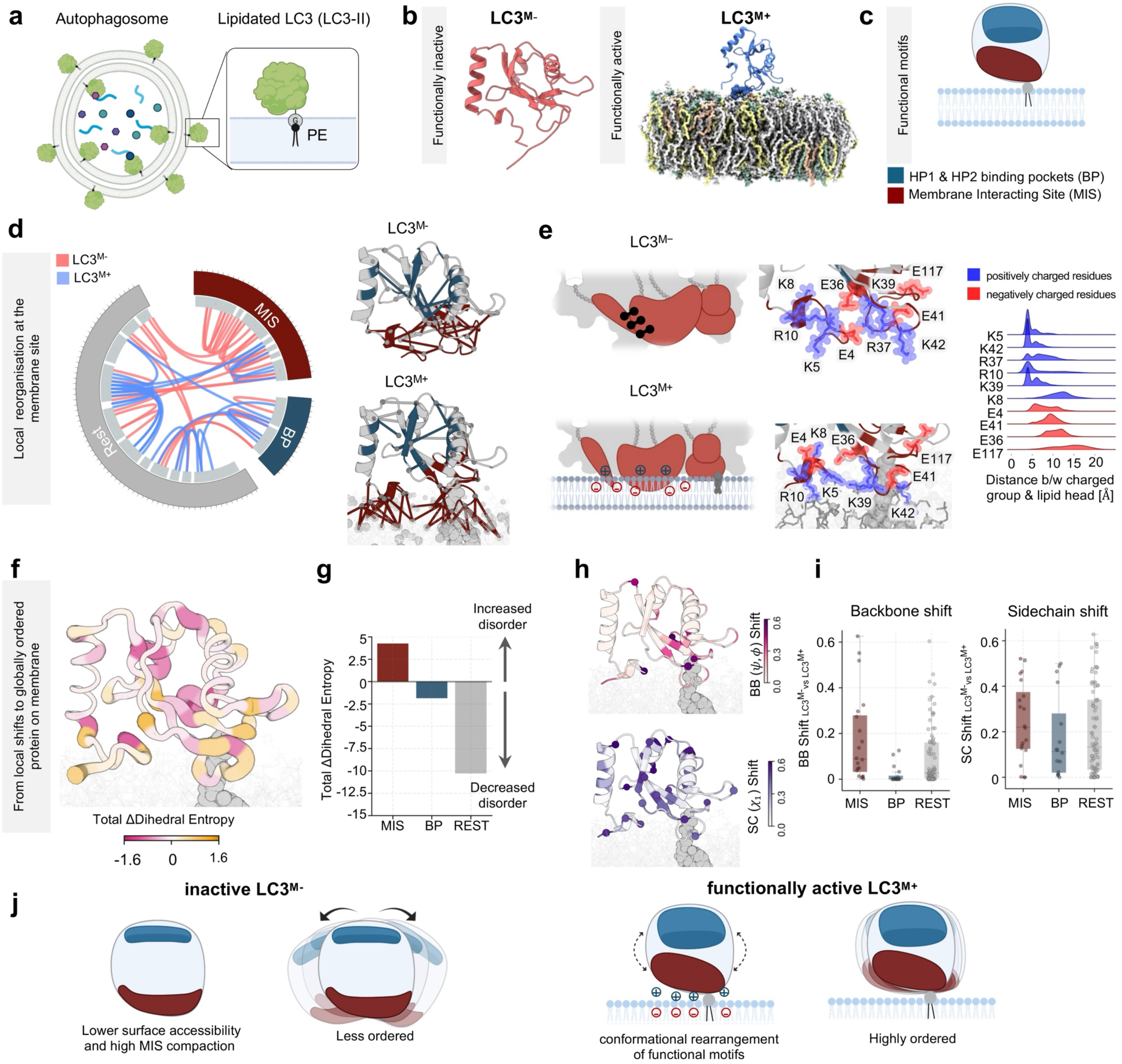
Membrane tethering induces conformational reorganization of LC3. (a) Schematic showing LC3 tethered to the autophagosomal membrane via a covalently attached PE chain. (b) The cytosolic (LC3^M-^) and membrane-bound (LC3^M+^) states of LC3 are shown in red and blue respectively. The ER-membrane lipids (DOPC, DOPE, DOPS, POPI) are depicted as colored sticks. (c) Two key functional motifs of LC3 are highlighted. The LIR-binding HP1 and HP2 pocket (BP), and membrane-interacting module (MIS). (d) Circos plots (left) illustrate state-specific intra-protein contacts for LC3^M−^ and LC3^M+^ states which are segmented into MIS, BP and rest of the protein. Each link denotes a contact unique to either state, with corresponding contacts mapped onto representative structures (right). For LC3^M+^, intra-protein and protein-membrane contacts are shown. (e) The schematic diagram (left) depicts the rewiring of the intraprotein network in the MIS. Zoomed-in snapshots (middle) reveal the side-chain orientation of charged residues within MIS where positively charged residues are shown in blue and negatively charged residues in red. The distribution plots (right) show the distances between charged residues within MIS and membrane surface in LC3^M+^. (f) The representative snapshot shows the difference in dihedral entropy (ΔEntropy= LC3^M+^ Entropy – LC3^M-^Entropy) in the putty representation. (g) The plot shows the distribution of total ΔEntropy calculated for three regions of the LC3 protein. Positive values indicate increased disorder, while negative values indicate decreased disorder. (h) The backbone and sidechain rotamer shift (LC3^M-^ vs LC3^M+^) data are mapped on the representative structure separately. (i) Backbone and sidechain shift values are plotted as box plots. The residues are grouped into three LC3 regions (j) Schematic representation summarizing LC3 conformational dynamics on the membrane. In the cytosolic state, LC3 is highly disordered and compact due to higher MIS compaction. Upon membrane association, local electrostatic rearrangements within the MIS trigger conformational shifts, resulting in a highly ordered global structure.

While intraprotein contact analysis revealed how interactions are rewired, quantifying flexibility captures how transitions between order and disordered regions of protein can modulate recognition preferences. To quantify conformational flexibility, we applied a Shannon entropybased structural order analysis, where higher entropy reflects conformational disorder and lower entropy reflects ordering^31^. This revealed persistent disorder within the MIS, particularly the N-terminus, whereas the remainder of the protein showed a marked entropy decrease, consistent with a stabilized membrane core upon membrane binding (**Fig. 1f-g, Supplementary Fig. 7**). These findings agree with NMR studies showing that the N-terminus samples multiple conformations^32^. To track local conformational changes, we performed rotamer analysis, which discretizes dihedral data into rotamer populations (backbone rotamers-*cis* and *trans*; sidechain rotamers-*gauche^-^*, *trans* and *gauche^+^*). Subsequently, we defined two metrics: *f*_*BB* *c*ℎ*an*g*e*_, as the fraction of residues with backbone rearrangements, and *f*_*SC* *c*ℎ*an*g*e*_, as the fraction of residues exhibiting side-chain rearrangements. While substantial backbone and sidechain rotamer shifts were found within MIS (*f*_*BB* *c*ℎ*an*g*e*_ =0.35, *f*_*SC* *c*ℎ*an*g*e*_ =0.55), the binding pocket showed localized sidechain adjustments (*f*_*BB* *c*ℎ*an*g*e*_ =0, *f*_*SC* *c*ℎ*an*g*e*_ =0.31), as shown in **Fig. 1h**. Interestingly, the rest of the protein not only gets more ordered (**Fig. 1g**) but also undergoes a significant shift in dihedral conformations (*f*_*BB* *c*ℎ*an*g*e*_ =0.18, *f*_*SC* *c*ℎ*an*g*e*_ =0.40; **Fig. 1i, Supplementary Fig. 8**). Therefore, these findings illustrate that membrane association reshapes intraprotein networks and converts localized disorder into ordered dynamics, highlighting the crucial role of lipids in driving LC3 conformational transitions (**Fig. 1j**).

### Lipid opens LC3 binding pockets via the α3–L5–β3-L6 allosteric switch

We next sought to understand the specific conformational changes in LC3 canonical binding pockets upon membrane attachment. LC3-LIR interactions are central to selective autophagy and adaptor recruitment. In particular, LIR peptide of p62 is extensively studied as an essential adaptor in selective autophagy^6^. To distinguish receptor-specific effects from membrane-induced changes, we also simulated the membrane-bound receptor state (LC3^M+R+^) along the apo state as described before (LC3^M+R-^) (**Fig. 2a**). The experimentally resolved structure of bound p62 LIR with LC3 was used to generate the LC3^M+R+^ state. Interestingly, from comparative analysis of binding pockets in cytosolic and membrane bound LC3 states, we found that out of 16 residues that participate in binding pocket, seven key residues in HP1 (D19, I23, K51) and HP2 (I35, F52, I66, R70) undergo in and out motions that govern pocket dynamics (**Fig. 2b; Supplementary Fig. 9**). As a result of these sidechain rearrangements, the canonical hydrophobic pockets HP1 and HP2 adopt distinct open conformation upon membrane association (**Supplementary Fig. 10**). As shown in **Fig. 2c**, across three independent trajectories, LC3^M−^ exhibits tightly packed hydrophobic pockets with average net volume 71.1 ± 2.8 Å³ and mean net inter-pocket distance of 10.1 Å. In contrast, binding pockets within the membrane-tethered states maintained an open conformation throughout the trajectories. LC3^M+R−^ expanded to 128.8 ± 2.4 Å³ and 10.9 Å intra-pocket distance, while LC3^M+R+^ further expanded to 175.4 ± 15.6 Å³ and 12.2 Å intra-pocket distance (**Supplementary Video 1**). Interestingly, a previous study has shown that K49, the gatekeeper residue of binding pockets, accommodates receptor binding via side chain flip^33^. Strikingly, in LC3^M+R-^ simulations, we found that the K49 sidechain flips out even in the absence of LIR, suggesting that lipids are priming receptor pockets for binding (**Supplementary Fig. 11**).

**Figure 2:**
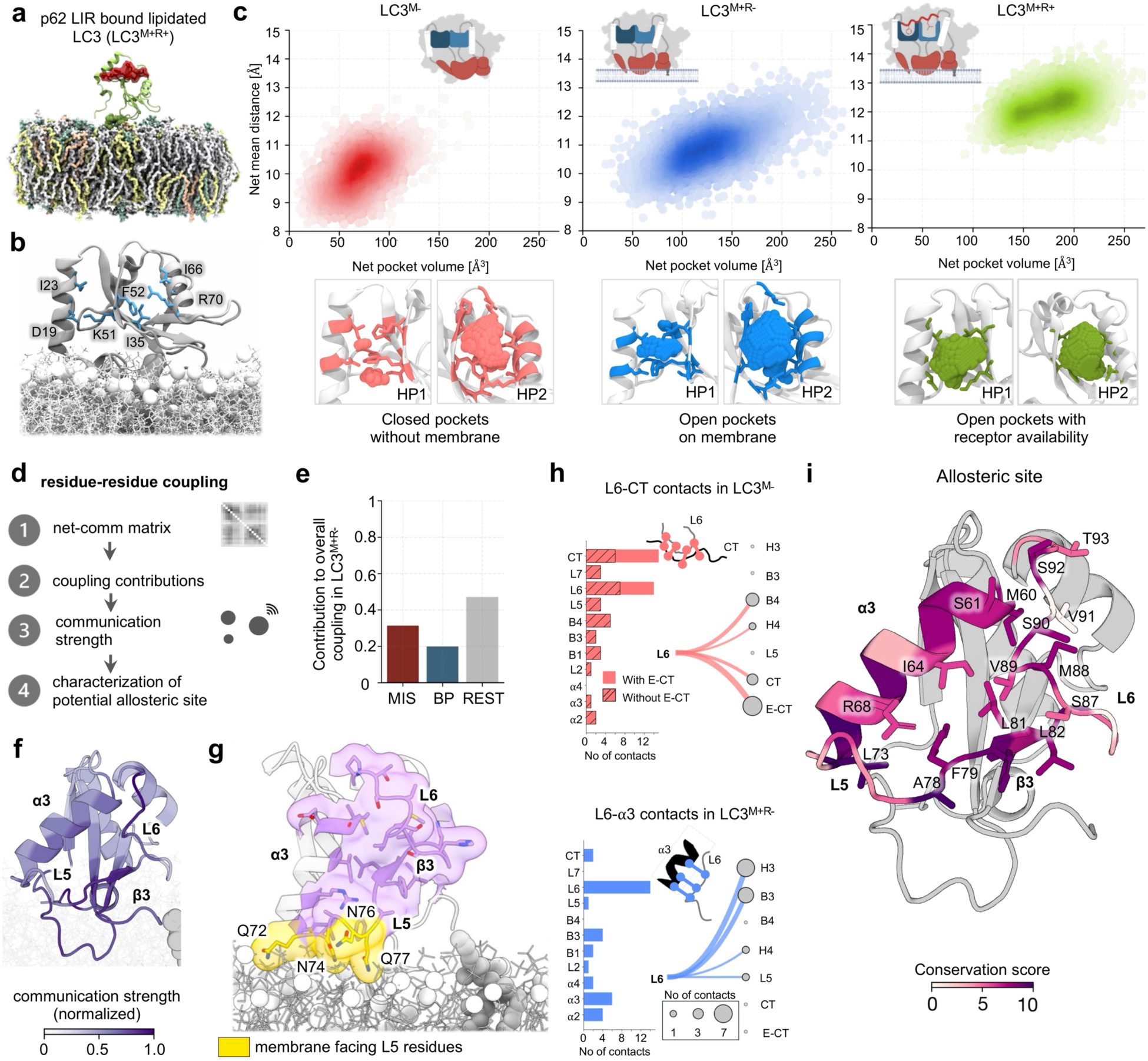
LC3 binding pockets undergo membrane-induced opening that is allosterically controlled by a conserved α**3-L5-**β3-L6 region. (a) The model represents the starting structure of LC3^M+R+^ (bound to p62-LIR) oriented on the ER-like membrane. The p62-LIR is highlighted as the transparent red surface. (b) The key residues (of HP1 and HP2) responsible for the pocket opening are shown as sticks and labelled. The snapshot corresponds to the representative structure of LC3^M+R+^ trajectories. (c) The net pocket volume is plotted against the net mean distance of pocket residues. The probability density distribution is shown for all three states of LC3. The schematic diagram shows the shape of binding pocket in respective states. (Bottom) The populated frames obtained from the density distributions are visualized. In the row below, snapshot displays HP1 and HP2 binding pockets, along with the pocket volume represented as vdW spheres. (d) The schematic diagram highlights key steps followed to compute residue-residue coupling. (e) The bar plot quantifies the contribution of each LC3 region (*i.e.,* MIS, BP and REST) to overall coupling. (f) Normalized communication strength computed for LC3^M+R-^ is mapped on the representative structure. The snapshot corresponds to the side-view of the protein. (g) The membrane-facing α3-L5-β3-L6 region residues (belonging to L5) are shown as yellow sticks along with transparent surface. The corresponding polar residues are labelled. Other allosteric site residues are colored purple. (h) The total number of unique contacts associated with each secondary structure of LC3 is plotted for both LC3^M-^ and LC3^M+R-^. Alluvial diagrams display unique contacts associated with L6. The thickness of the links and the size of the nodes represent the count. (i) The allosteric site is colored according to the conservation score, and the residues are shown as sticks.

Several LIR-containing receptors bind to LC3 for selective autophagy function^34^. In the context of chemical diversity of LIR motif, the LIRs can be classified into three types based on the amino acid at the Φ position (**Φ**-xx-Г): W-type, F-type, and Y-type, examples include p62, Optineurin (OPTN), and neighbor of BRCA1 gene 1 (NBR1), respectively. We next investigated whether the binding pocket opening mechanism is conserved across other LIRs. We simulated LC3 bound to OPTN and NBR1. Both the hydrophobic pockets showed consistent membrane-induced pocket opening, with volumes of 198 ± 25.3 Å³ and 169 ± 24 Å³, respectively (**Extended Fig. 1**), indicating that this conformational change is not restricted to a specific LIR type.

Since LC3 undergoes global reorganization upon membrane binding that extends beyond the membrane interface and canonical pocket, we asked whether these changes reflect long-range communication across the protein **(Fig. 1h)**. To address this, we performed residue-residue dynamic correlation analysis, which measures how the motion of one residue is coupled to another over the course of simulations^35^. From dynamic correlation, we derived net communication matrices and calculated per-residue communication strength, a normalized metric (0-1) that reflects how strongly a residue communicates with the rest of the protein **(Fig. 2d)**. Residues with high communication strength (>0.5) serve as dynamic hubs that propagate structural signals and mediate long-range coupling. As expected, within functional regions MIS and BP coupling increases upon membrane binding **(Supplementary Fig. 12)**. Interestingly, nearly half of the total coupling emerged from outside the functional regions, suggesting additional sites participate in the dynamic cooperativity (**Fig. 2e**). Indeed, analysis of per-residue communication strength revealed a distinct hub spanning α3, loop 5 (L5), β3, and loop 6 (L6) region **(Fig. 2f)**. Despite being spatially distal (∼14.8 Å from the HP2 pocket center), this region was strongly coupled to the canonical binding pocket (**Supplementary Fig. 13**). In particular, this site primarily comprises of residues in α3 (M60, S61, I64, R68), L5 (L73, A78, F79), β3 (F80, L81, L82) and upper L6 (S87 to T93) exhibit the highest mean communication strength (∼0.7). Comparison of previous experimentally resolved structures and simulation ensemble from our data display a triangular topology, with the above-mentioned residues facing inwards **(Supplementary Fig. 14)**. The base of triangular topology consists of L5 residues that are membrane-facing and polar residues (Q72, N74, N76, and Q77) participate in transient dynamic interactions with polar lipid head-groups (**Fig. 2g**, **Supplementary Fig. 15**). Next, we probed distance between the sides of triangular topology (α3-L6) to understand the conformational dynamics. Surprisingly, when compared across states during the entire length of trajectories, allosteric site conformations differ **(Supplementary Fig. 16)**. In the cytosolic LC3^M-^ state, L6 is away from α3 and is highly flexible and samples multiple orientations near the receptor-binding groove. However, LC3^M+R-^, upper L6 comes closer to α3 to form a tighter and stable triangular topology. This is evident from interaction preferences, whereby L6 engages with α3 and L5 in membrane-bound state rather than β4 and the C-terminal region as observed in LC3^M-^ state (**Fig. 2h)**. We, therefore, define this region as an allosteric site within LC3 that relays membrane-induced changes to the LIR binding interface. Notably, this site is highly conserved with 60% of residues showing high conservation (**Supplementary Fig. 17, Fig. 2i**). As shown in numerous studies, allosteric modulation in terms of conformational change can directly impact protein function^7^. From the above findings, we propose that lipid binding induces conformational changes within allosteric site that may regulate LC3 activity.

### Protein-ensemble centric design of allosteric site

We hypothesized that modulating the conformation of allosteric site shift the thermodynamic equilibria, and as a consequence alter the protein function. From above molecular insights, we aimed to conformationally mimic: (i) a dynamic allosteric site with closed pockets and impaired function, as observed in LC3 ^M-^, and (ii) a stabilized allosteric site seen in LC3^M+^ trajectories with open binding pockets primed for receptor engagement. To test this, we used a dynamics-guided design strategy, exploiting time-resolved trajectories to engineer mutants biased toward these alternative states (**Fig. 3a**). We introduced a series of allosteric site mutations that can potentially strengthen the triangular topology to induce increased stability as active mutant candidates or disrupt local interactions to increase flexibility within allosteric site, indicating inactive candidates.

**Figure 3:**
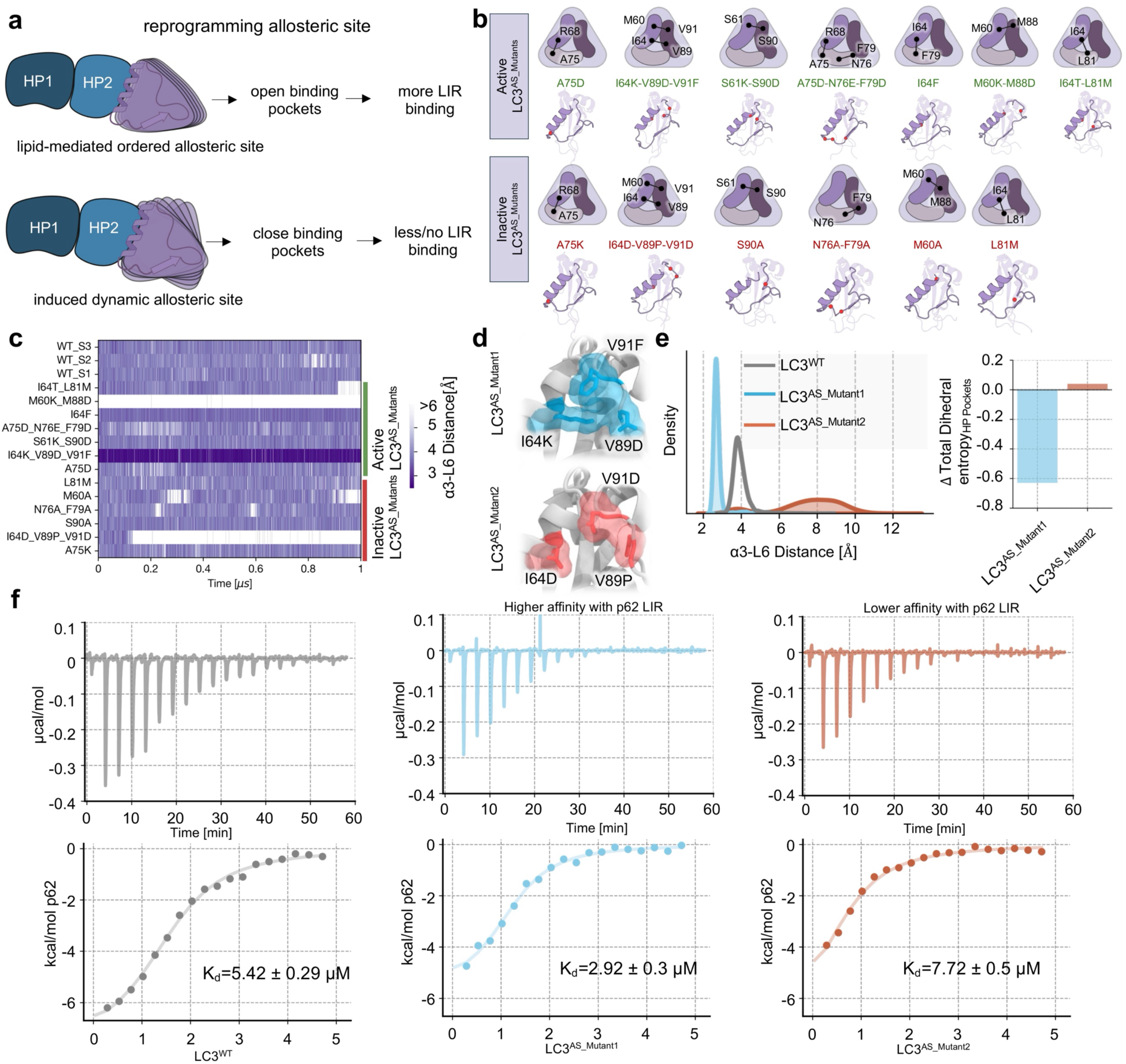
MD-guided design of active LC3^AS_Mutant1^ and inactive LC3^AS_Mutant2^. (a) The schematic diagram illustrates the rationale behind generating active LC3^AS^ and inactive LC3^AS^ mutant candidates using the ensemble-centric approach. (b) The mode of residue-residue interaction intended to be perturbed is shown on the schematic representation of the α3-L5-β3-L6 region (for each mutant). The snapshots from the last frame of each trajectory are shown along with the mutation sites marked as red spheres. (c) (Left) The probing distance for the α3-L5-β3-L6 region (minimum distance between residue positions 64 and 89) is plotted as a heatmap for each mutant candidate as a function of simulation time. (d) The mutation positions are visualized as sticks along with a transparent surface. (Note: All the simulations are performed on the ER-like membrane; LC3^M+R-^). (e) The probing distance distribution is plotted for the chosen mutant candidates-active LC3^AS_Mutant1^ (I64K-V89D-V91F) and inactive LC3^AS_Mutant2^ (I64D-V89P-V91D), along with LC3^WT^. (Right) The bar plot quantifies the total ΔEntropy of hydrophobic pockets (ΔEntropy = EntropyMut – EntropyWT). (f) The ITC data plots show binding of LC3^WT^, LC3^AS_Mutant1^, and LC3^AS_Mutant2^ mutants to p62 LIR peptide. The calorimetric titration for p62 titrated into LC3^WT^ is shown and derived values for Kd are shown. The experimental conditions were kept consistent across all the experiments.

In total thirteen mutants were designed, six were predicted to favor the inactive LC3^M+^ state, while seven favored the active LC3^M+^ state. Each variant was simulated for 1 µs, with allosteric site compactness used as a probe of conformational state (**Fig. 3b-c**). Among these, a triple mutation targeting residues on the sides of the allosteric site (I64 in α3, V89 and V91 in L6) predicted an interesting trend. Here, mutations favoring interactions within allosteric region I64K-V89D-V91F exhibited a stabilized allosteric site, with distances confined to 2.8 ± 0.36 Å (**Supplementary Video 3**). In contrast, I64D-V89P-V91D disrupted the local interactions of the allosteric site with perturbed hydrophobic interactions and a kink in L6 via the proline substitution inducing flexibility. Therefore, we selected these two mutations and refer to former mutation as LC3^AS_Mutant1^ and the latter as LC3^AS_Mutant2^ (**Fig. 3d**). Both mutants preserved the ubiquitin fold of LC3 as well as membrane interactions, with positively charged residues (K5, R10, R37, K39) mediating lipid contacts and acidic residues (E4, E36, E41, E117) remaining solvent-exposed (**Supplementary Fig. 18**). Pocket disorder analysis confirmed this distinction, with LC3^AS_Mutant1^ showing a reduction in conformational disorder relative to the LC3^WT^, whereas inactive LC3^AS_Mutant2^ showed a slight increase (**Fig. 3e**).

To validate these predictions, we performed isothermal titration calorimetry (ITC) assay using the p62 LIR peptide (Figure 3c). LC3^WT^ showed a K_d_ of 5.42 ± 0.19 µM with the peptide. The LC3^AS_Mutant1^ showed stronger binding (K_d_ = 2.92 ± 0.3 µM, ∼2-fold increase) while LC3^AS_Mutant2^ mutant exhibited weaker binding (K_d_ = 7.72 ± 0.5 µM, ∼1.5-fold decrease) (**Fig. 3f; Supplementary Table 2**). Thermodynamic analysis revealed altered entropy contributions for both mutants, consistent with conformational reorganization upon p62 engagement (**Supplementary Fig. 19**). Together, these results confirm our predictions that LC3^AS_Mutant1^ is showing receptor binding due to changes within allosteric site that stabilize the “orthosteric” binding pocket. On the other hand, LC3^AS_Mutant2^ shows 1.5-fold decreased binding, implying closed inactive conformation. This demonstrates that distal allosteric regulation can define alternative functional states of LC3 and directly modulate receptor binding affinity.

### Structural basis of stabilization in active LC3^AS_Mutant1^ revealed by X-ray crystallography

To define the structural basis of stabilization, we solved two crystal structures of the active LC3^AS_Mutant1^ (I64K-V89D-V91F), in apo form and in complex with the p62-LIR peptide (**Fig. 4a**; **Table 1**). Structural comparison with LC3^WT^ revealed that the engineered substitutions induce pronounced local rearrangements at the allosteric site (**Fig. 4b**). In particular, mutant residues I64K and V91F, together with wild-type residue F79, form a continuous π–cation interaction axis that reinforces L6 into a stable position. The second side-chain shift was observed in L6 of the triangular topology. Residue H86 flips inward in both mutant structures, engaging with E102 via NE2 (2.7 Å) while its ND1 interacts with the backbones of N84 and G85 (**Fig. 4c**). These features are consistent with our MD simulations where we predict an ordered allosteric site in LC3^AS_Mutant1^.

**Figure 4:**
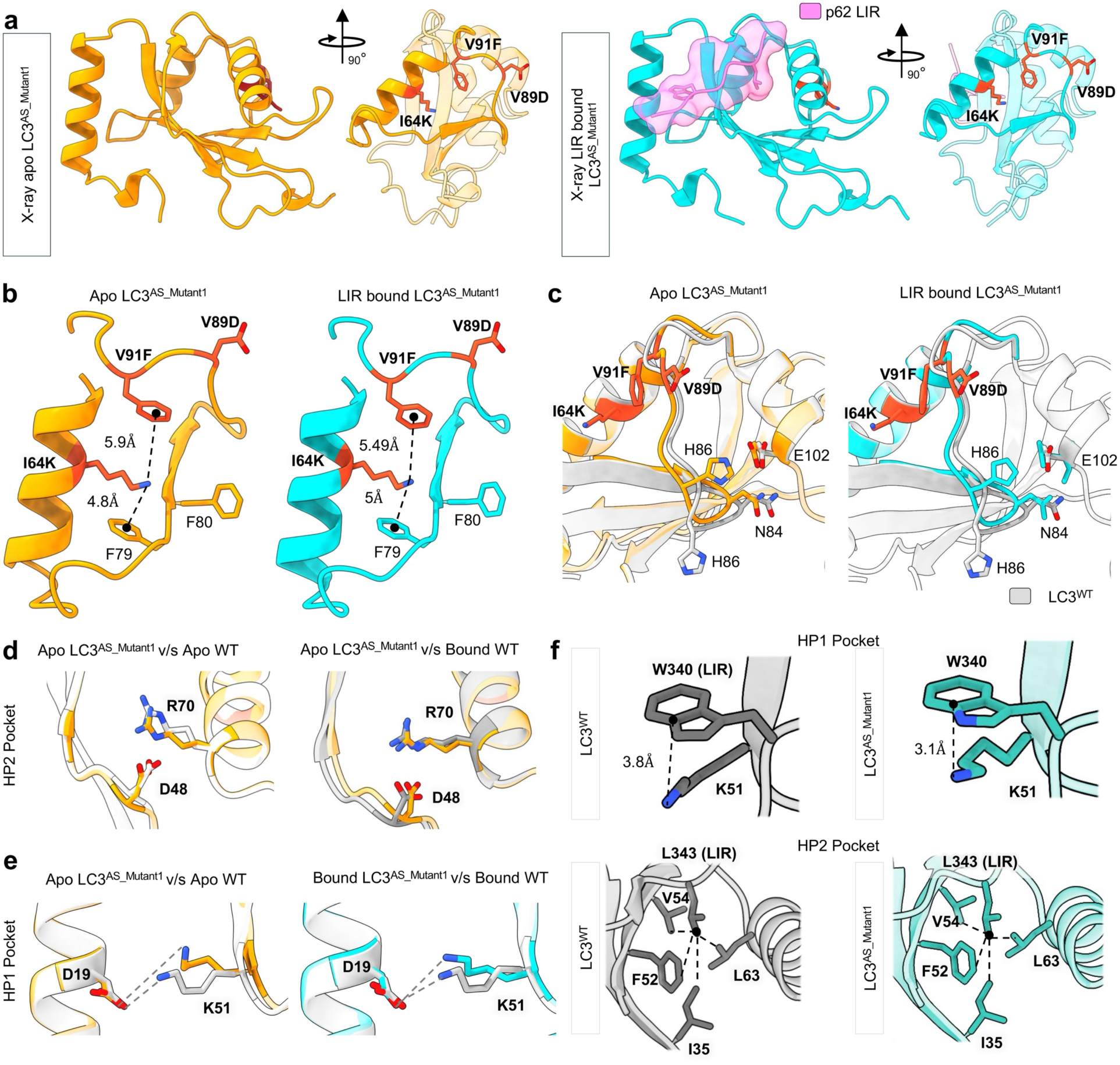
X-ray crystallography captures changes induced by mutations in functional binding pockets. (a) X-ray crystal structures of active LC3^AS_Mutant1^ without p62-LIR (left) and with p62-LIR (right) are visualized. Both front and side views are shown. The mutated residues are shown as sticks in the side view. The p62-LIR is visualized as magenta cartoon with transparent surface. Critical LIR residues (W340, L343) are shown as sticks. (b) The α3-L5-β3-L6 region is shown, and the mutated residues are visualized as dark orange sticks. π-cation interactions V91F-I64K and F79-V64K are highlighted. The distances are measured between the NZ atom of I64K and the aromatic center of phenylalanine residues-F79 and F91. (c) The relative orientation of basal L6 residues H86 and N84 with respect to α4-residue E102 is shown. The orientation is compared with the WT crystal structure in complex with p62-LIR (PDB: 2ZJD) (grey cartoon) via superimposition. (d) Changes in the relative orientation of electrostatic residue pair R70-D48 in apo LC3^AS_Mutant1^ are shown. The orientation is compared with the corresponding WT structure in the absence (left, PDB: 3VTU) and presence of p62-LIR (right, PDB: 2ZJD). (e) (Left) The relative orientation of D19-K51 in apo LC3^AS_Mutant1^ is compared with the apo WT structure (PDB: 3VTU). (Right) The same pair in the presence of LIR is compared with the WT LIR-bound structure (PDB: 2ZJD). (f) The orientation of K51 relative to the critical LIR residue W340 is shown for both WT (PDB: 2ZJD) and active LC3^AS_Mutant1^ crystal structures bound to p62-LIR. (Bottom) The depth of p62-LIR residue L343 into the HP2 pocket is visualized for both WT and active LC3^AS_Mutant1^ crystal structures. The relevant HP2 residues I35, F52, V54, and I63 are marked as sticks.

**Table 1.**
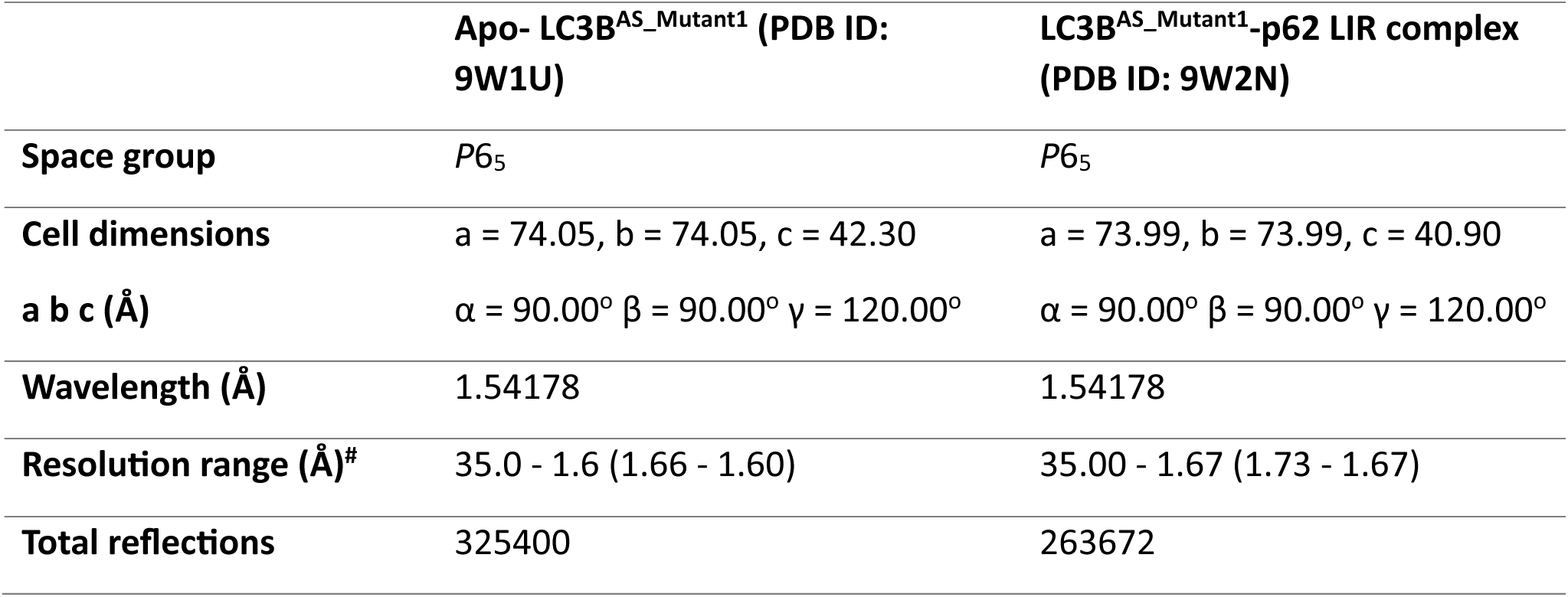
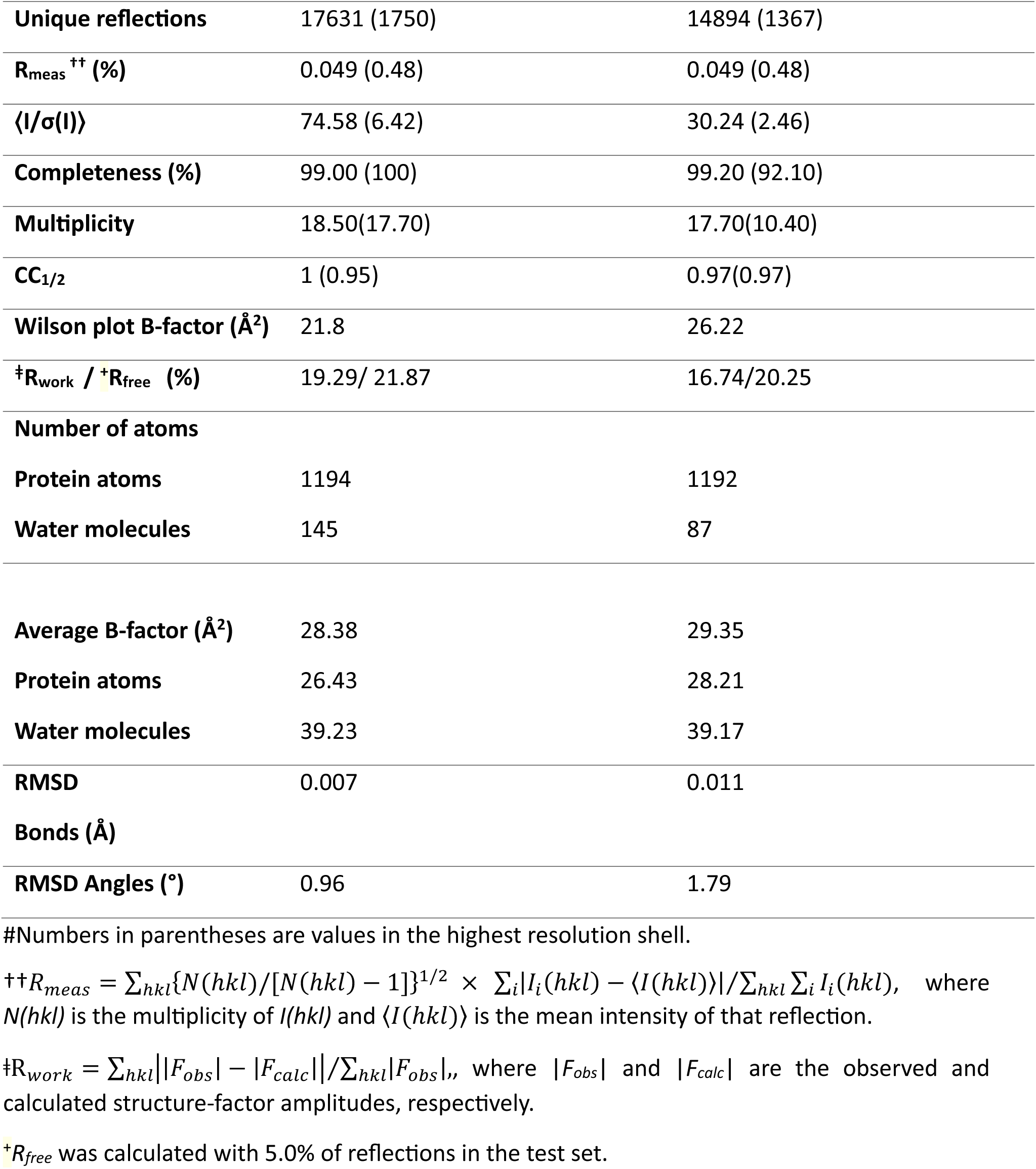
Summary of data collection and refinement statistics for Apo LC3B^AS_Mutant1^ mutant and p62 LIR-bound LC3B^AS_Mutant1^ structures.

Remarkably, stabilization of the allosteric site also influences the canonical binding pocket. Superimposition of apo LC3^AS_Mutant1^ with wild type apo structure revealed that the HP2 pocket closely mimics the LIR bound conformation. Residue R70 of HP2 structurally resembles the ligand-engaged conformation (**Fig. 4d**). Because of this, the mutant structures display a reinforced R70– D48 salt bridge compared to LC3^WT^. In addition, compared to wild-type, the HP1 pocket in the apo mutant shows changes in the relative orientation of conserved residue position K51 with respect to D19 whereby the bound mutant structure shows a subtle side chain shift. When comparing apo LC3^AS_Mutant1^ and apo WT, the distance increased by 0.93 Å in the mutant, whereas 0.51 Å increase was observed when comparing LIR-bound LC3^AS_Mutant1^ and LIR-bound WT (**Fig. 4e**). As a proof of concept, the bound mutant structure is similar to wild-type bound with all other changes intact including the gatekeeper residue K49 maintains its position, facilitating LIR accommodation, while K51 forms stronger hydrogen bonding interactions with W340 of the p62 peptide. From the p62 standpoint, in mutant structures, the depth of L343 of p62 LIR bound to HP2 relatively increased (**Supplementary Fig. 20**). These rearrangements coincide with a subtle reorientation of the p62 peptide at HP2 relative to the wild-type LC3–p62 complex, suggesting enhanced stabilization of the peptide in the active mutant (**Fig. 4e**). Several ordered waters observed in LC3^WT^ (e.g., HOH 144 in 2ZJD) are displaced, widening the pocket and reducing the energetic cost of accommodating the bulky tryptophan side chain of the LIR motif.

Together, these high-resolution structures establish the structural basis of the active LC3^AS_Mutant1^. The stabilization of the allosteric site is transmitted to the canonical binding pocket, predisposing the LC3 mutant toward the stable conformation and facilitating tighter LIR binding. The comparative analysis showed that in the active LC3^AS_Mutant1^ apo structure, the pockets are primed for LIR binding. Importantly, the observed features mirror those predicted in our MD ensembles, providing direct structural validation of the dynamics-guided design approach.

### The lipidated active LC3^AS_Mutant1^ facilitates cargo sequestration

To determine whether the active LC3^AS_Mutant1^ (I64K-V89D-V91F) and inactive LC3^AS_Mutant2^ (I64D-V89P-V91D) differ from LC3^WT^ in their ability to sequester the autophagy substrate p62/SQSTM1, we studied them using autophagic cellular assays. Three constructs: eGFP-tagged LC3^WT^, active LC3^AS_Mutant1^, inactive LC3^AS_Mutant2^ constructs were assessed for p62 sequestration. All cells were co-transfected with the mCherry-tagged autophagy substrate p62, following which we examined whether the inactive LC3^AS_Mutant2^ shows reduced colocalization with p62, while the active LC3^AS_Mutant1^, by virtue of its stable substrate binding pocket, displays greater p62 colocalization, when compared to the eGFP-tagged-LC3^WT^ control. Consistent with our design principle that stabilization of the allosteric site promotes pocket stabilization, active LC3^AS_Mutant1^ displayed substantially greater colocalization with p62 **(Fig. 5a and 5c)**, which appears in most instances as complete engulfment of mCherry-p62 puncta by eGFP-labelled LC3/autophagosomes. Increased colocalization of active LC3^AS_Mutant1^ with p62 did not appear to relate to changes in size or number of autophagosomes (data not shown). In contrast, the inactive LC3^AS_Mutant2^ showed reduced colocalization with mCherry-p62 **(Fig. 5a and 5c)**, although this reduction did not acquire statistical significance. Supporting these observations, co-immunoprecipitation of eGFP-tagged-LC3^WT^ or active LC3^AS_Mutant1^ or inactive LC3^AS_Mutant2^, revealed greater (∼8-fold) binding of the active LC3^AS_Mutant1^ with mCherry-p62, when compared to control LC3 **(Fig. 5b)**, while as noted via immunofluorescence, the inactive LC3^AS_Mutant2^ showed no difference in p62 binding when compared to control LC3 **(Fig. 5b)**. Taken together, these results show substantially increased affinity between LC3 and p62 when LC3 is mutated at I64K, V89D, and V91F supporting our hypothesis that stable pocket conformations of LC3 allow greater affinity towards substrate peptides. Consistently, transmission electron microscopy (TEM) showed greater instances of autophagosomes with clearly identifiable sequestered organelle cargo, e.g., endoplasmic reticulum (indicated by yellow arrowheads) and mitochondria in cells expressing the active LC3^AS_Mutant1^ when compared to those expressing LC3^WT^ **(Fig. 5d)**. TEM also confirmed that expressing LC3^WT^ or active LC3^AS_Mutant1^ mutant had no impact on the ability to form autophagosomes, indicated by intact double-membraned structures (shown by red arrows) in both conditions. These data demonstrate that active LC3^AS_Mutant1^, by its ability to engage with cytoplasmic autophagic cargo, shows greater ability to sequester cargo in autophagosomes when compared to LC3^WT^ controls. To determine if this greater ability to sequester cargo in autophagosomes by the active LC3^AS_Mutant1^ translates to enhanced cargo turnover in lysosomes, we examined the effect of expression of the active LC3^AS_Mutant1^ on lysosomal flux of defined organelle receptor proteins, which are known to be degraded by macroautophagy. Strikingly, expressing the active LC3^AS_Mutant1^ led to increased turnover of endogenous LC3 (p<0.05), p62 (p<0.01), FAM134B (p<0.05) [the selective ERphagy receptor]^36^, and a trend towards increased turnover of VPS4A (p<0.09) [the selective lipophagy receptor]^37^, and a trend towards greater turnover of LC3^AS_Mutant1^ mutant per se when compared to LC3^WT^ **(Fig. 5e-f; Supplementary Fig. 21)**. Taking together these results strongly support the idea that the active LC3^AS_Mutant1^ facilitates enhanced cargo sequestration as well as degradation in lysosomes.

**Figure 5:**
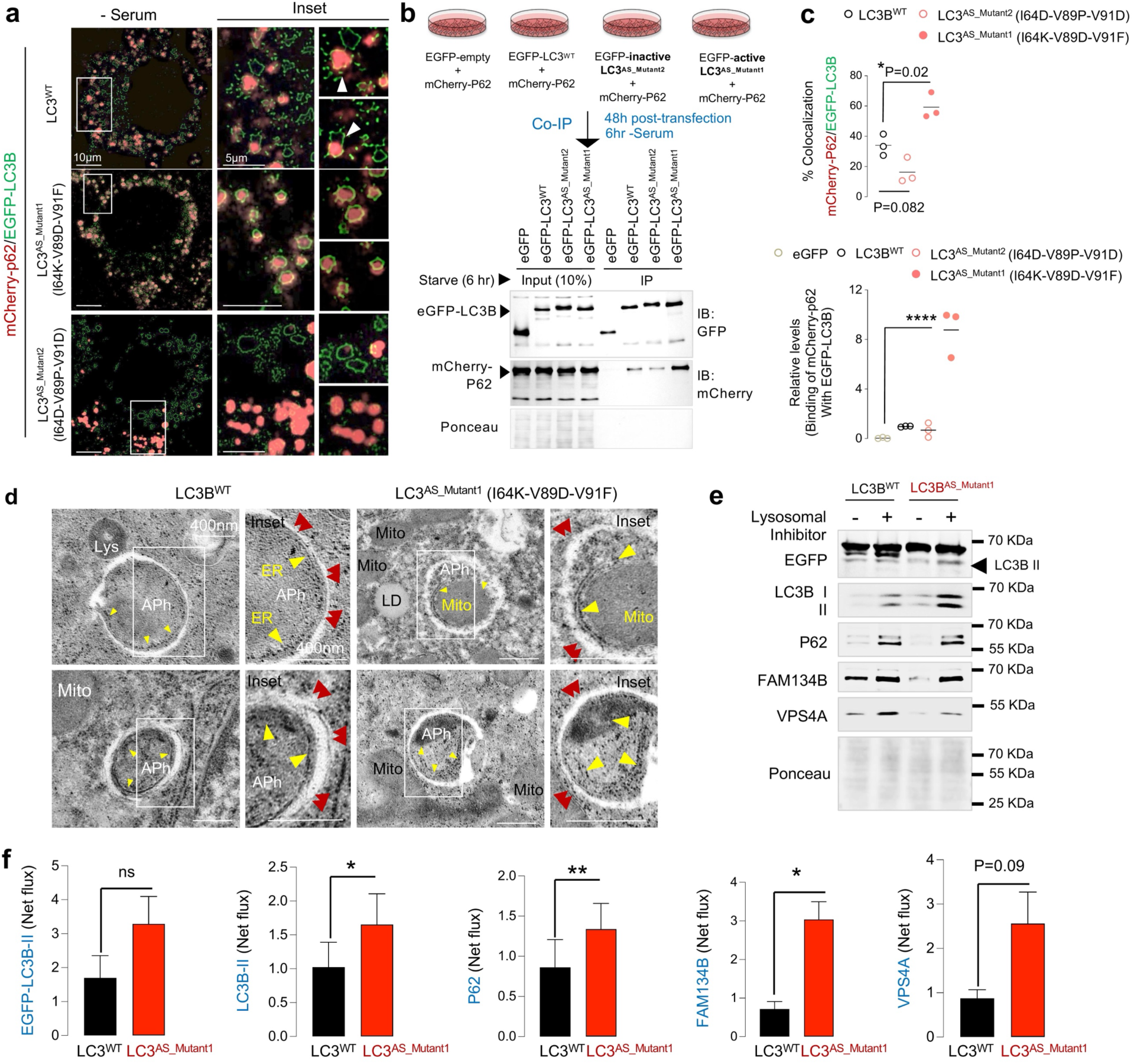
The lipidated active LC3^AS_Mutant1^ facilitates cargo sequestration. (a) Direct fluorescence for EGFP-LC3 (LC3^WT^ or indicated mutants) and mCherry-P62 in AML12 hepatocytes and subjected to serum starvation for 6 hrs (n=3). At least 30 cells were analyzed per replicate. Quantification for % colocalization is shown. (b) Co-IP of EGFP-LC3 (LC3^WT^ or indicated mutants) with mCherry-P62 in AML12 hepatocytes as per plan indicated (n=3). (c) Quantification for % colocalization (images in a) and binding of mCherry-p62 with EGFP-LC3 (blots in b) are shown. (d) TEM in livers expressing LC3B^WT^ or the active LC3^AS_Mutant1^ (I64K-V89D-V91F) depicting double-membraned autophagosomes (indicated by red arrows) engulfing cargo (ER indicated by yellow arrowheads) and mitochondria (Mito). Magnified insets are shown. Magnification bar = 400nm. (e) Immunoblots for indicated proteins in AML12 hepatocytes expressing LC3B^WT^ or active LC3^AS_Mutant1^ (I64K-V89D-V91F) in presence or absence of lysosomal inhibitors (n=3). (f) Densitometric quantification of relative flux under the indicated conditions. Relative flux was calculated as the ratiometric quantification of densitometric values (+Inhibitor / –Inhibitor) for each condition. Mean values are shown. p<0.05, *p<0.01, ***p<0.0001, ns = not significant; one-way ANOVA and Tukey corrected or unpaired Student’s t-test.

## Conclusions

Long-range coupling of lipids and proteins is a recurring theme in biology^38^, yet the underlying molecular mechanisms by which lipid interactions modulate protein functional sites remain poorly understood. In this work, we identify a membrane-coupled structural mechanism of LC3 regulation that is intrinsic to the protein itself. Leveraging biomolecular simulations, we show that membrane binding induces a global conformational reorganization of LC3 **(Fig. 1)**. This process of direct lipid binding rewires electrostatic interactions within membrane-interacting regions and reshapes intramolecular contacts, thereby opening the hydrophobic pockets that recognize LIR motifs. Via long-range communication analysis, our work reveals that LIR pocket accessibility of LC3 is dynamically coupled to a distal α3–loop5– β3–loop6 allosteric site, which becomes ordered upon membrane association and serves as a long-range regulator of binding-site conformation **(Fig. 2)**. While lipids often bind directly to membrane-embedded proteins to transmit conformational information^39^, we demonstrate that dynamic and transient lipid engagement in the vicinity of an allosteric site is sufficient to induce its ordering. By reprogramming LC3 allosteric site, we generated variants that remain membrane-associated yet are directly governed by this regulatory site (**Fig. 3**). In particular, mutations designed to destabilize or stabilize the allosteric site were sufficient to bias LC3 into alternative functional states, inactive LC3^AS_Mutant2^ (I64D-V89P-V91D) and active LC3^AS_Mutant1^ (I64K-V89D-V91F). High-resolution crystallography of active LC3^AS_Mutant1^ revealed a stabilized allosteric site and subtle yet functionally relevant rearrangements within the binding pocket, providing structural validation of the MD-predicted mechanism (**Fig. 4**). Complementary experiments, including super-resolution fluorescence microscopy, TEM and autophagy flux assays further confirmed that active LC3^AS_Mutant1^ exhibited enhanced binding and cargo sequestration (**Fig. 5**). Together, these findings show that LC3 function is governed by membrane-induced conformational dynamics and illustrate how lipid-mediated allostery, widely recognized in channels and receptors^39^, also operates in a small globular protein central to autophagy.

## Methods

### Generation of starting structures with membrane

The full-length human cytosolic LC3B (125 a.a.) structure (modelled from PDB ID: 3VTU as template) was taken from our previous study ^29^, whereas the lipidated structural model of human LC3B protein was taken from another study of ours^27^. The lipidated structures were inserted in a physiologically relevant bilayer membrane of 400-lipids, prepared with the CHARMM-GUI web server^40^. The membrane contains ER-like lipid composition including DOPC (65%), DOPE (20%), POPI (10%), and DOPS (5%)^29^. The membrane was equilibrated and simulated using the parameters files given by CHARMM-GUI, before placing it with the lipidated structure. The orientation of the lipidated structure was taken our previous study^28^. To generate the LIR-bound lipidated structures, we utilized the crystal structure coordinates of p62 (PDB ID: 2ZJD)^21^, Optineurin, OPTN (PDB ID: 2LUE)^22^, Neighbor of BRCA1 Gene 1, NBR1 (PDB ID: 2L8J)^23^ LIR. Both lipidated and crystal structures were superimposed, and the coordinates for LIR peptide were saved in complex with lipidated LC3. The p62, OPTN, or NBR1 LIR bound to lipidated LC3B were placed in the ER-like membrane and subjected to simulations.

### All-atom MD simulations

We simulated three states of LC3 protein: cytosolic (LC3^M-^), membrane-bound apo (LC3^M+R-^) and membrane-bound LC3 in complex with p62-LIR (LC3^M+R-^) (**Supplementary Table 3**).

i. **Membrane-bound (LC3^M+R-^/ LC3^M+R+^) simulations:** In the membrane-bound lipidated LC3 simulations (LC3^M+R-^ and LC3^M+R+^), the lipidated protein was placed on the ER-like membrane, utilizing our previous conformation^28^, in a rectangular box (dimension: 115.3 Å x 115.3 Å x 140.0 Å). The water molecules with TIP3P representation, and Na^+^ counter ions were added to neutralize the systems. The CHARMM36 all-atom forcefield^41^ was incorporated, and the simulations were performed with GROMACS 2018.3^42^. The electrostatic interactions were computed with the Particle Mesh Ewald summation with the grid spacing set at 0.16nm^43^. The van der Waals cut-off was set to 1.2 nm, and LINCS constraints were applied^44^. The steepest descent method was applied for energy minimization. Two-step equilibration runs (NVT and NPT) were carried out, restraining the heavy atoms. During the production run, the temperature was maintained at 310 K using Nose-Hoover thermostat with the temperature coupling (τ_T_) set at 0.5 ps^45^. The pressure was maintained at 1 bar using semi-isotropic Parrinello-Rahman barostat, and the pressure coupling (τ_P_) was set at 2ps and compressibility of 4.5 x 10^-5^ bar^-1^ along XY axes^46^.
ii. **Cytosolic state (LC3^M-^) simulations:** The starting structure of the cytosolic simulations were obtained from our previous study^29^. The simulation box size was adjusted to match that used in the membrane-bound systems, and three independent replicas were performed to maintain consistency in total simulation time with the membrane-bound trajectories.

The time step of 2 fs was used for the numerical integration of the equation of motion. The coordinates were saved every 20 ps. Three replicas for each LC3^M-^, LC3^M+R-^, and LC3^M+R+^ systems were simulated for 1 μs, and the cumulative simulation time reached 9 μs. With the same parameter settings, we also simulated the LC3^M+R+^ system bound to NBR1 (PDB ID: 2L8J) and OPTN (PDB ID: 2LUE) LIRs (1 μs each), totaling 2 μs with GROMACS 2021.4^42^.

### Intra-protein contact analysis

The residue-residue contacts across different trajectories were computed with the in-house Python script compiled with MPI support. If any of the heavy atoms in a residue pair are within a 5Å cut-off distance, the residue pair (Ri, Rj) was considered to be in contact at the frame f. This was iterated throughout the trajectory, and contact probabilities were calculated for all the combinations of residue pairs. The residue pairs with a contact probability > 0.5 were considered to have stable contacts. With this criterion, the contact probability matrix was discretized into a simple contact matrix where the value 1 denotes the stable contact (between the residues Ri, Rj) and the value 0 denotes the lack of stable contact. To extract the unique contacts that are specific for one particular form of LC3, the discretized contact matrices corresponding to two different LC3 forms (states) were subtracted.

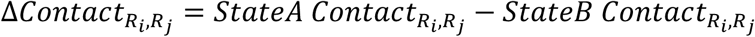

If Δ*Contact*_*Ri*,*R*j_ is 1, then the corresponding contact is specific for state A. In case of -1, the corresponding contact is specific to state B.

### Computing residue-wise disorderness

To compute the dihedral disorderness of a residue from the trajectory, we calculated dihedral entropy. The rotamer states of the dihedral values across time were assigned as explained in Singh et al^35^. For each residue dihedral (𝜓, 𝜙 *and* 𝜒_1_), we calculated Shannon entropy *H* of its rotamer states (*Cis* and *Trans* for backbone; *Gauche^-^*, *Trans* and *Gauche^+^* for side chain) observed across the simulation. Shannon entropy of a dihedral 𝑋 (*H*(𝑋)) is expressed as

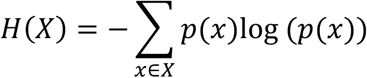

where 𝑋 can be 𝜓, 𝜙 *and* 𝜒_1_. 𝑥 represents the rotamer state of the dihedral 𝑋 can adopt. The probability of a dihedral 𝑋 being in a rotamer state 𝑥 is computed in 𝑝(𝑥).

Once separately computing the Shannon entropy for 𝜓, 𝜙 *and* 𝜒_1_ of a residue, we sum up all the entropies and apply a normalization scheme by dividing the maximum entropy value out of three dihedrals. This results in “net dihedral entropy” of a residue^31^. Thus, net entropy of a residue *R*_*i*_ is as follows.

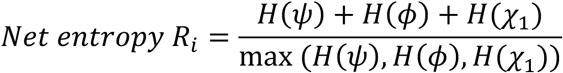

### Computing rotamer shift

To screen the dihedral changes between two forms of LC3, we utilized the time evolution of rotamers. We define rotamer shift as the mean change in dihedral (𝜓, 𝜙, *and* 𝜒_1_) rotamer populations between two LC3 states. Thus, the rotamer shift for the residue *R*_*i*_is given by the following equation.

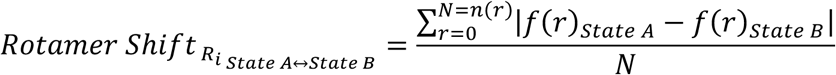

The 𝑟 denotes the rotamer state of the dihedral. For backbone dihedrals, the rotamer states include *Cis* and *Trans*. For the sidechain, the rotamers include *Gauche^-^*, *Trans* and *Gauche^+^*. Thus, 𝑁 is the total number of rotamers for a dihedral (backbone or sidechain). Each of the rotamer fractions *f*(𝑟) (fraction of a rotamer 𝑟 of a residue throughout the trajectory) is computed for two forms of LC3 and subtracted. The rotamer shift is then defined as the average of these differences in the rotamer fractions. The resulting shift varies from 0 to 1. For the backbone rotamer shift of a residue, we picked the maximum shift observed from psi (ψ) and phi (Φ). For sidechain rotamer shift, we only considered 𝜒_1_. Rotamer change of residue (backbone or sidechain) is considered to be substantial if the obtained shift value is greater than 0.2.

### Pocket volume calculations in MD trajectories

Epock tool^47^ was utilised to get the pocket centre coordinates for volume calculations. Using POVME 3.0^48^, the binding site volume in each trajectory was calculated as a function of time. Probe size used for computing volume is 6 Å, whereas each grid is 0.5 Å apart.

### Computing residue-residue correlations from dihedral dynamics

To understand the influence of each residue over the other, dihedral and dihedral-dependent parameters were utilized. Initially, all residue dihedrals (ψ, ϕ for backbone and χ1 for sidechain) were computed across the trajectory with *gmx_chi*. From the residue dihedrals, the rotamer and orderness states were extracted. While rotameric states were assigned as explained in a previous study^35^, the orderness states were derived from the time-evolution of rotamer states (of the corresponding dihedral) following the k-neighborhood scheme (see supplementary methods). Thus, these three parameters, such as raw dihedral angles, their rotamer and orderness states for each residue were considered to quantify residue-residue correlations. We define residue-residue correlations as the sum of normalised Mutual Information (MI) between residues Ri and Rj calculated for the backbone (BB) and sidechain (SC) dihedrals.

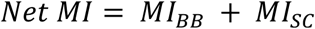

where 𝑀𝐼_*BB*_ = 𝑀𝐼: + 𝑀𝐼_𝜙_; 𝑀𝐼_*SC*_ = 𝑀𝐼*_x_*_1_

Mutual Information of a dihedral type (BB or SC) is the sum of individual MI calculated for dihedrals, rotamer and orderness states. Individual matrices were normalized by dividing by the obtained maximum value (otherwise known as channel capacity).

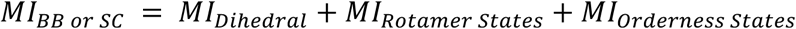

The *MI_Dihedral_* was computed as implemented in Kraskov et al^49^ since dihedral angles are continuous variables, while the other two (*MI_Rotamer States_, MI_Orderness States_*) were calculated as explained in Singh et al^35^ (see supplementary methods). The resulting net MI matrices were normalized by the same scheme and further used for computing net communication. Net communication underlying residues *R_i_, R_j_* is the sum of MI between *R_i_* and *R_j_* neighborhoods. We define the neighborhood as 3Å around the residue selection. The net communication matrix was then normalized by the same max-scaling approach. We refer to the net communication between two residues as their coupling. From the normalized net communication matrix, the communication strength of each residue was computed. The communication strength of residue *R_i_* is defined as the sum of net communication between residue *R_i_* and the rest of the residues in LC3 protein.

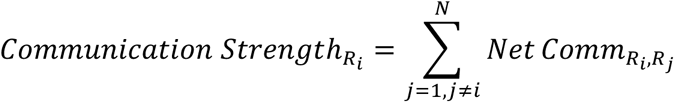

Total coupling between two segments/regions was calculated by summing the coupling for every residue pair *R*_*i*_ *R*_j_ belonging to the segments *Se*𝑔_*i*_ *Se*𝑔_j_.

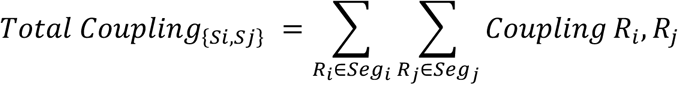

We further computed the relative contribution of the coupling between a given region and the rest of the protein to the overall net-communication. This is explained in the supplementary methods in detail. All these calculations were performed with in-house Python scripts (Github repository: https://github.com/CSB-Thukral-Lab/LC3_conformations_scripts_data).

### Mutant LC3 All-atomistic simulations

The mutant candidates chosen to mimic the allosteric site dynamical states were simulated. The mutations were pooled into two categories: i) potential allosteric site stabilizing mutants and ii) allosteric site destabilizing mutants. To enhance the stability of allosteric site, we introduced complementary charged interactions (A75D, I64K-V89D-V91F, S61K-S90D, A75D-N76E-F79D, and M60K-M88D) and, in parallel, hydrophobic or polar interaction network that can strengthen the triangular topology (I64, and I64T-L81M). Alongside, we designed another set of mutant candidates expected to destabilize the allosteric site by disrupting the interaction network by inducing charge-repulsion or disrupting polar/hydrophobic network (S90A, N76A-F79A, M60A, and L81M). We generated simulation data for these 13 candidates, totaling 13 μs (1 μs each) (**Supplementary Table 4**). The mutations were introduced to the WT structure of the LC3B^M+^ on the ER membrane. The introduction of mutations was done with the Chimera tool^50^ by choosing the most favorable Dunbrack rotamer^51^. Using the same orientation and protocol followed for WT, the mutant systems embedded in ER-like membrane were set up and run using GROMACS 2021.4^42^ with CHARMM36 forcefield.

### Geometric Analysis

All distance and angle calculations were done with the MDAnalysis Python library^52^. Detailed methods for computing various geometric parameters throughout the simulation are provided in the supplementary methods. VMD^53^, ChimeraX^54^, and Pymol^55^ tools were utilised for structural visualization and rendering. Python libraries Matplotlib^56^, Seaborn^57^ and Pycirclise^58^ were used for data plotting.

### Isothermal Titration Calorimetry (ITC)

#### Protein purification

Constructs for mice LC3B^WT^, LC3B^AS_Mutant2^(I64D-V89P-V91D), and LC3B^AS_Mutant1^ (I64K-V89D-V91F) mutants were cloned from an eGFP pcDNA3.1 plasmid into the pet28a vector. The constructs were confirmed via Sanger sequencing. Overexpression was performed in Rosetta cells and Induction was carried out at an OD600 of 0.5–0.7 by adding 0.2 mM β-D-1-thiogalactopyranoside (IPTG). The cells were kept at 20°C for 22–24 hours in a shaker incubator. The cell pellet was resuspended in lysis buffer (20 mM sodium dihydrogen phosphate, pH 7.5, 1.2 M sodium chloride, and 10 mM imidazole). Cells were lysed using a sonicator, and the lysate was applied to a His Trap Chelating HP column charged with nickel(II)sulphate to purify (His)6-tagged LC3B^WT^, LC3B^AS_Mutant1^, and LC3B^AS_Mutant2^ mutants. The column was washed with wash buffer (20 mM imidazole) and proteins were eluted using elution buffer (250 mM imidazole). Further purification was performed via gel filtration chromatography (FPLC) using a Superdex 75pg column (Cytiva) into the binding buffer (20 mM HEPES, pH 7.5, 200 mM NaCl, and 0.5 mM Tris(2-carboxyethyl) phosphine (TCEP). The purified proteins were kept at 4°C until further use. The peptide p62 (sequence: DDWTHLS) was synthesized by Genescript and resuspended in a binding buffer and subsequently used for further experiments.

#### Isothermal Titration Calorimetry (ITC) assay

Calorimetric titrations were performed at 25°C using a Microcal PEAQ-ITC (MicroCal, LLC). The proteins were exchanged in a binding buffer through gel filtration. The titrations involved 19 injections of 2 μL each, where p62 peptide at concentrations of 480-500 μM was titrated into 300μL of 20 μM LC3B WT and mutant constructs. Each experiment was conducted in triplicate. The integrated heat data from the titrations were analyzed using the MicroCal PEAQ-ITC analysis software, employing the fitted offset option to automatically subtract the control heat. The resulting isotherm was fitted using a one-site binding model.

### LC3B^AS_Mutant1^ X-ray Crystallization

#### Construction of plasmids for expression in *E. coli*

The agar stabs for the LC3B^AS_Mutant1^ were obtained from Gene to protein (G2P) Pvt. Ltd in peT28a(+) vector in *E. coli* cells. Primer details were as follows: LC3B^AS_Mutant1^-I64KV91F-FP, 5’-GTACATATGCCGTCCGAGAAGACCTTCA-3’. The amplified cDNA PCR product was inserted between NdeI and XhoI restriction sites into the pET28a+ expression vector (Novagen). The positive clones were validated by DNA sequencing results.

#### Over-Expression and Purification of recombinant LC3B^AS_Mutant1^ mutant

LC3B^AS_Mutant1^ was purified as a 125-amino acid recombinant protein bearing N-terminal 6x-histidine affinity tag with a thrombin cleavage site. Protein was recombinantly expressed in Escherichia coli in Rosetta cells with isopropyl β-D-1-thiogalactopyranoside (IPTG) induction. The culture was incubated overnight at 37 °C, in a shaker incubator (220 rev min^-1^) and induced for 4 hrs using 1mM IPTG. Cell pellets were either stored at −80 °C or directly used in the following protein purification.

The bacterial culture pellets for the LC3B^AS_Mutant1^ were suspended in a lysis buffer containing 100 mM Tris (pH 8.0), 200 mM NaCl and a trace amounts of lysozyme, DNase I (Roche, Germany) and one tablet of Complete EDTA free protease inhibitor mix (Roche, Basel, Switzerland) and subjected to sonication over ice. The lysed pellet was centrifuged at 11,000 RPM for 60 min at 4°C. The supernatant was collected, and affinity purification was performed with a pre-packed 1 ml HisTrap HP column (Cytivia cat no. 17524701). It was first equilibrated with 100 mm Tris-HCl buffer (pH 8.0), 200 mM sodium chloride and 5 mM imidazole, thereafter supernatant loaded manually. The elutions were collected and loaded on 15% SDS-PAGE gel for further analysis. The confirmed fractions were concentrated and loaded to a HiLoad 16/60 Superdex 75 gel filtration column, (GE Healthcare, USA) pre-equilibrated with 100 mm Tris-HCl buffer (pH 8.0) and 100 mm NaCl. Fractions from the middle of the peak corresponding to monomeric protein were collected and loaded on SDS PAGE gel for clarity. The pure protein fractions were pooled, concentrated using a 10,000 da membrane filter to 18-20 mg/ml. Final concentrations of protein was determined by measuring absorbance at 280 nm using Nanodrop spectrophotometer (Thermo Scientific). The protein aliquots were flash frozen in liquid Nitrogen and stored at −80 °C until further usage.

#### p62 peptide synthesis

The 11-mer p62 peptide was purchased from Gene2protein (G2P) (Hyderabad, INDIA). The synthesized peptide was purified by HPLC to achieve >98% purity and presented in lyophilized form with TFA salt. The identity of the peptide was confirmed with mass spectrometry (**Supplementary Fig. 22**), and the sequence (SGGDDDWTHLS) of the peptide is listed in Table 1.

#### Crystallization & structure determination

All crystallization trials were carried out in 96-well sitting drop plates using the sitting drop vapor diffusion method. Crystallization experiments were performed on mosquito robotic system Xtal3 (sptlabtech) at National Institute of Immunology (NII), INDIA by varying protein and cryoprotectant ratios.

### Apo structure

The crystals for LC3B^AS_Mutant1^ were obtained in 200 mM ammonium sulphate, 25% PEG 4000, 100 mM sodium acetate pH 4.6, at 24 °C. Crystals appeared in sitting-drop vapour-diffusion setup in about two weeks. They were transferred into the cryoprotectant solution (reservoir solution with 20% (v/v) ethylene diol), followed by flash-cooling at -173 °C in liquid nitrogen.

### p62-LIR peptide bound structure

After the initial crystallization screening, good single crystals were also obtained within 3 weeks, in a drop containing 2 M sodium potassium tartrate tetrahydrate, 200 mM ammonium sulfate, and 100 mM sodium citrate (pH 5.6). The crystal soaking procedure was performed to obtain the protein–peptide complex in this condition. The p62-LIR peptide was dissolved in water, and a 1:10 molar ratio of protein: peptide was used for setting up soaking experiments. 1 µL of the high concentration peptide was used for setting up soaked protein crystals at different intervals (1-2 weeks) and equilibrated against the same reservoir buffer condition.

### Data Collection

The diffraction data for the crystals was collected at 100 K at the in-house x-ray diffractometer at National Institute of Immunology (NII) having CuKα radiation (1.5418 Å) generated by SuperBright rotating anodes on an RIGAKU FR-E^+^ SuperBright coupled to an image plate detector R-AXIS IV++. The data were processed and scaled with HKL2000 program suite^59^. The crystals belonged to *P6_5_* hexagonal space group, with unit cell parameters of *a* = 73.64 Å, *b* = 73.64Å, and *c* = 40.51Å and α= 90°, β=90°, γ=120°. The apo crystals contained one molecule in the asymmetric unit, calculated Matthew’s coefficient and solvent content values were 2.0 Å^3^/Da and 38.12% respectively. X-ray data collection, scaling and refinement statistics are summarized in Table 1.

### Structure determination and refinement

The diffraction data was indexed, integrated and scaled using the HKL2000. The structure was solved by molecular replacement in *Phaser-MR*^60^, CCP4.4 (Collaborative Computational Project, Number 4, 1994) software suite. LC3B protein in the PDB: 2ZJD^21^ was used as the model template for p62 bound structure. Rigid body refinement, followed by iterative cycles of restrained atomic parameters was performed. TLS refinement was done in REFMAC^61^. Initial model building visualization of the electron density maps, was carried out in Coot^62^. MolProbity^63^ was used to assess the stereochemistry of both the structures.

Both 2|*F_o_*|-|*F_c_*| and |*F_o_*|-|*F_c_*| Fourier maps from the peptide-soaked crystals permitted the tracing and building of p62 peptide to LC3B^AS_Mutant1^ structure in the asymmetric unit. The initial phases of the structure were determined by molecular replacement, using 2ZJD as the search template. Molecular replacement was performed with MOLREP in REFMAC for calculation of initial phases. The structures were modified manually with Coot and refined with *PHENIX_Refine*.

Phasing and refinement statistics are summarized in Table 1. In both structures, there were no residues in the disallowed regions of the Ramachandran plot. At the end of the refinement cycles, water molecules were fitted through inspection of different Fourier maps. The final refined structures of apo LC3B^AS_Mutant1^, in the asymmetric unit contained 1020 protein atoms, and 143 water molecules, with a final R_work_ of 19.29% and R_free_ of 21.87% at 1.60 Å resolution. While the LC3B^AS_Mutant1^ -p62 LIR complex possesses 1090 protein atoms and 87 water molecules. The final refined structures of the complex LC3B^AS_Mutant1^ -p62 LIR complex, in the asymmetric unit contained 1090 protein atoms, and 143 water molecules, with a final R_work_ of 16.74% and R_free_ of 20.25% at 1.67 Å resolution. The final structure factors and structural coordinates have been submitted to PDB with ID 9W1U, 9W2N respectively for apo-LC3B^AS_Mutant1^ and p62 LIR-bound _LC3BAS_Mutant1._

### Cell culture

AML12 cells (ATCC, CRL-2254) were cultured in DMEM/F-12 medium (Gibco, 11320033) supplemented with 10% fetal bovine serum (FBS) (Gibco, 10082), 1% Insulin-Transferrin-Selenium (Gibco, 41400-045), 40 ng/ml dexamethasone (Sigma-Aldrich, D4902) and 1% penicillin-streptomycin (P/S) (Gibco,15140). Cells were maintained at 37°C in 5% CO2. Wherever indicated, cells were washed once with PBS and incubated in serum-free DMEM/F-12/P/S in presence or absence of 0.25 mM oleic acid (OA) (Cayman Chemical, 29557) for the indicated durations.

### Image acquisition, processing and quantification

Images were acquired using optically demodulated structured illumination super resolution microscopy (SIM-Live SR) equipped with Nikon-Ti2 CSU-W1 spinning disc confocal with FI60 Plan Apochromat Lambda D 100x Oil Immersion Objective Lens, N.A. 1.45, W.D. 0.13mm, F.O.V. 25 mm, DIC, Spring Loaded. Images were captured using Kinetix 22 back-illuminated sCMOS Camera for C-mount, 22 mm FOV.2400x2400, 83FPS@16-bits, 6.5 μm pixel. Image pixels were 0.107 and 0.065 μm, respectively. Blue, green, red, and far-red fluorescence was excited by 405, 488, 560, or 637 nm lasers. All laser lines possess a quad-pass dichroic beam splitter (Di01-T405/488/568/647). Spinning disc was configured (CSU-W1) with Live SR super-resolution modality, 3D deconvolution and Denoise.ai to improve both axial and lateral resolution and allowed to uncover structural and molecular details at subcellular resolution (∼105 nm). Images were acquired at similar exposure times in the same imaging session for each experimental replicate. All images were subjected to identical post-acquisition processing methods and similar metrics.

Cells were transfected with indicated plasmid DNA and seeded onto a glass bottom 27 mm culture dish (Thermo Scientific, 150682) for 48 h. After the corresponding treatments, cells were washed once with PBS, incubated in live cell imaging solution (Invitrogen, A14291DJ), and imaged using a Nikon-Ti2 CSU-W1 spinning disc confocal microscope, and single planes and Z stacks (∼ 0.2 µM) were acquired with ×100 objective and 1.45 numerical aperture. Images were captured with up to 1000 fps and real-time 3D-image visualization with minimal phototoxicity to the samples. Image analysis Image processing and analysis was performed in NIS Elements AR and Fiji (ImageJ; NIH). Individual frames were denoised by applying Denoise ai and deconvoluted using an advanced 3D Deconvolution algorithm in NIS elements. NIS AI and General Analysis 3 (GA3) modules were used for advanced image processing and quantitative analysis. Percentage colocalization was calculated using the JACoP plugin. Mander’s M1 colocalization coefficient was used as a quantitative metric. A minimum of 30 cells per condition per experimental replicate were quantified and analyzed.

### GFP-tagged LC3 co-immunoprecipitation

48h post transfection, AMl12 cells co-expressing GFP–LC3^WT^ or GFP-LC3B^AS_Mutant2^ or GFP-LC3B^AS_Mutant1^ with mCherry–p62 were starved for 6h in the DMEM/F12 medium without FBS. GFP empty vector were used as negative controls. Lysates (500 μg of proteins) were incubated with Anti-GFP magnetic bead slurry (GFP-Trap magnetic agarose kit, Chromotek, gtmak-20) for 1 h at 4 °C in rotation. Bound proteins were eluted by boiling (95 °C for 5 min) in 2 × SDS–PAGE sample buffer. Immunoprecipitated (IP) proteins and original lysates (input) were resolved on an SDS–PAGE, and membranes were probed for GFP and m-Cherry. Bands obtained by immunoblotting were quantified using ImageJ software and normalized to corresponding mCherry-p62 changes.

### Transmission electron microscopy

Liver tissue slices were fixed in 2.5% glutaraldehyde and 2% formaldehyde in 0.1 M sodium cacodylate buffer for 1hr. After wash, samples were embedded in 4% agarose gel and post-fixed in 1% osmium tetroxide. After wash, samples were dehydrated through a graded series of ethanol concentrations and propylene oxide. After infiltration with Eponate 12 resin, the samples were embedded in fresh Eponate 12 resin and polymerized at 60°C for 48 hours. Ultrathin sections of 70 nm thickness were prepared and placed on formvar-coated copper grids, then stained with uranyl acetate and lead citrate. The grids were examined using a JEOL 100CX transmission electron microscope at 60 kV and images were captured by an AMT digital camera (Advanced Microscopy Techniques Corporation, model XR611) (Electron Microscopy Core Facility, UCLA Brain Research Institute). Image processing performed with Fiji^66^. "Enhance Contrast" feature was applied, with "saturated pixels" set to 0.35% and "equalize histogram" selected. The Image was converted to 8-bit, and the "Despeckle" feature was applied. The number of autophagosomes (APh) and deformities in autophagosome structures were observed and quantified by visual inspection of the recorded CCD micrographs.

### Autophagy Flux

Autophagy flux was determined 48h post-transfection in AMl12 cells co-expressing GFP–LC3^WT^ or GFP-LC3B^AS_Mutant2^ or GFP-LC3B^AS_Mutant1^ following serum starvation for 5 hr in the DMEM/F12 medium without FBS. One hr after FBS removal, cells were exposed or not to lysosomal inhibitors (20mM NH4Cl and 100µM leupeptin) for 4 hr prior to harvest. Lysates were subjected to immunoblotting and turnover rates of various autophagy receptors was determined by subtracting the densitometric value of the non-inhibited sample from the corresponding value of the inhibited sample.

### Sequence analysis

The LC3 ortholog dataset was obtained from the Autophagy3D repository^65,66^. Multiple sequence alignment was performed with MAFFT v7.490^67^. From MSA, the conservation score was computed following the method described in Livingston and Barton., 1993^68^ (as implemented in the Jalview tool). The algorithm quantifies the number of shared physicochemical properties among residues in each alignment column. A score of 0 suggests no common properties among the aligned residues, whereas a score of 10 denotes conservation of all the physicochemical properties. The scores range from 0 (indicating no shared physicochemical property) to 10 (complete conservation of all physicochemical properties defined by the method).

## Data and Code availability

The data files for the atomistic simulations of LC3B^M-^, LC3B^M+^, and LC3B^M+R+^, along with OPTN-LIR-LC3B^M+R+^, NBR1-LIR-LC3B^M+R+^, LC3B^AS_Mutant2^, and LC3B^AS_Mutant1^ mutants will be made available. The scripts and data from the simulations will be made available in the GitHub link https://github.com/CSB-Thukral-Lab/LC3_conformations_scripts_data. The high-resolution figures and supplementary videos are available in the figshare link https://figshare.com/s/f009879f892b56df8e0f. The coordinates and structures of the mouse apo LC3B^AS_Mutant1^ mutant and p62 LIR-bound LC3B^AS_Mutant1^ mutant complex have been deposited into the Protein Data Bank (PDB) under accession codes 9W1U and 9W2N, respectively.

## Acknowledgements

LT is thankful for the funding support from the DBT/Wellcome Trust India Alliance (IA/21/2/505925). D.G was supported by CSIR doctoral fellowship, and JC was supported by DBT/Wellcome Trust India Alliance. SM acknowledges the research grant from the Science and Engineering Research Board (SERB), DST WOS-A fellowship for (DST/WOS-A/LS-383/2021). NJ would like to thank the DBT Ramalingaswami fellowship grant (35/2019). We are grateful for the generous help provided at the X-ray diffraction facility at BRIC-National Institute of Immunology (NII), India. LT acknowledges the support from CSIR-IGIB for infrastructure and CSIR-4PI for supercomputing facilities. LT acknowledges CSIR-IGIB intramural grant OLP2503 for the workforce support. We would like to thank all the CSB lab members for support and constructive feedback on our work. We also thank Dr Ullas Kolthur Seetharam and Dr Arvind Penmatsa for providing insightful comments and suggestions that helped improve the manuscript.

## Ethics declarations

Competing interests

L.T., D.G., and N.J. are inventors on a patent application describing a mutant LC3B allosteric switch for autophagy regulation.

## Extended data figures

**Extended Figure 1.**
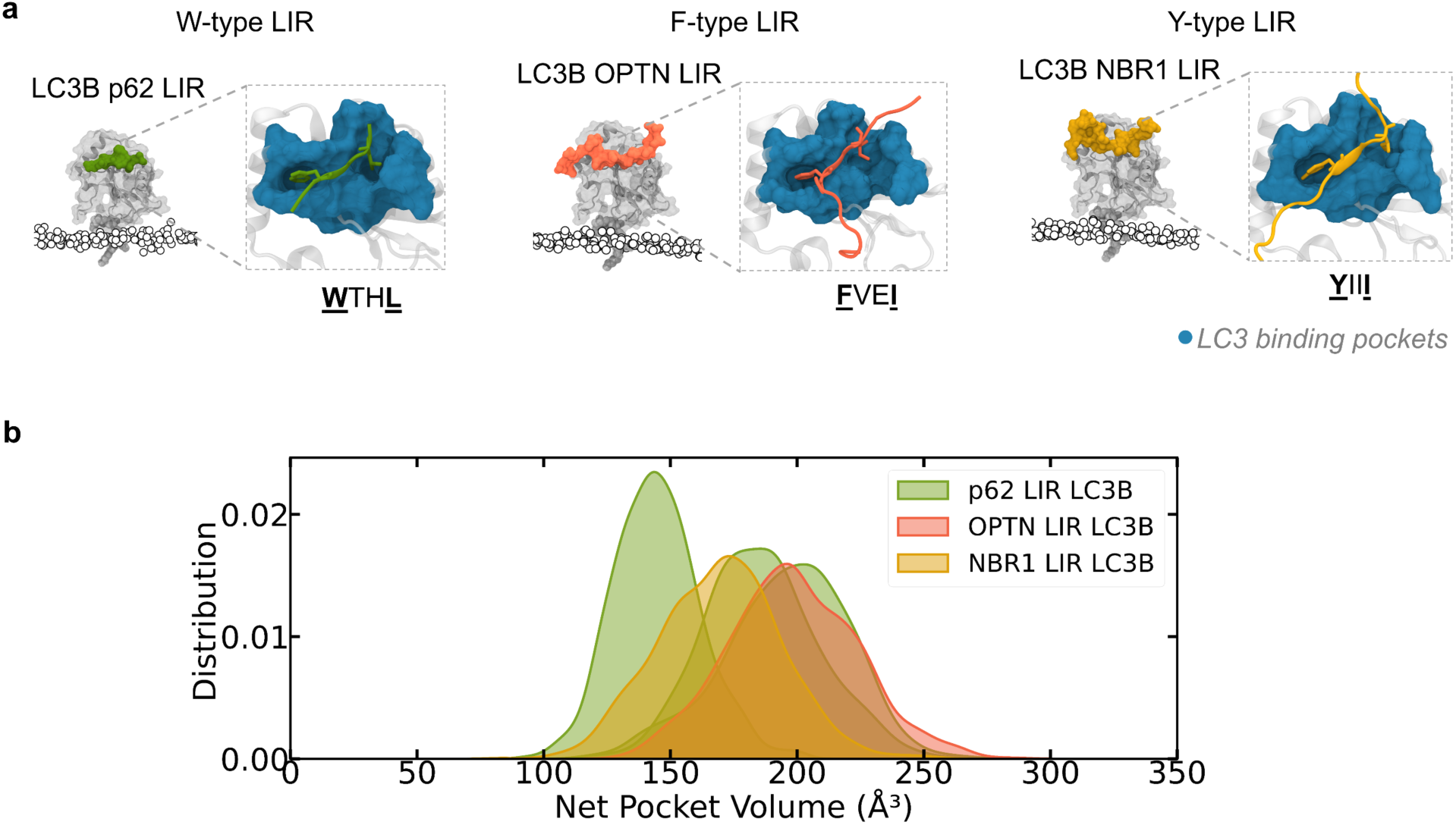
The pocket opening is a general mechanism for W/F/Y-type LIRs. (a) The structural snapshots display the membrane-embedded, lipidated LC3 bound to W-type, F-type, and Y-type LIR peptides, represented in green, orange, and yellow, respectively. The LIR peptides are shown in surface representation, while the LC3 protein backbone is depicted using both cartoon and transparent surface representations. Membrane head group phosphates are highlighted as spheres. A zoomed-in view illustrates the LC3-LIR interaction, with side chains of residues at the Θ and Γ positions of the LIR motif shown in stick representation. The LC3 receptor binding pockets are displayed in teal surface representation. The corresponding LIR motif sequences are provided below. (b)The distribution plots depict the net pocket volumes of membrane-embedded LC3^M+R+^ bound to different LIR peptides.

## Source Data

### Source Data Fig.1

Numerical source data from MD simulations (state-specific contacts, distances between charged residues and lipid head, dihedral delta entropy and rotamer shift)

### Source Data Fig.2

Numerical source data from MD Simulations (binding pocket landscape, region-wise contribution to overall coupling, residue-wise communication strength, secondary structure-associated unique contacts and conservation score of the allosteric site)

### Source Data Fig.3

Numerical source data from MD Simulations (Allosteric site α3-L6 distances for all mutant candidates, binding pocket entropy for selected mutant designs).

Statistical source data from ITC experiments

### Source Data Fig.5

Statistical source data from cell biology experiments (% of LC3/p62 colocalization for LC3^WT^ and LC3^Mutants^, relative levels of eGFP LC3-mCherry p62, net flux of LC3 and various receptors)

### Source Data Extended Data Fig.1

Numerical source data from MD simulations (HP1, HP2 and net volume data for p62-LIR, OPTN-LIR and NBR1-LIR bound simulations)

## Supplementary information

Supplementary information is provided in a separate PDF.

## New structures

## References

1. Changeux, J.-P. & Christopoulos, A. Allosteric Modulation as a Unifying Mechanism for Receptor Function and Regulation. Cell 166, 1084–1102 (2016).

2. Wang, J. et al. Mapping allosteric communications within individual proteins. Nat Commun 11, 3862 (2020).

3. Vithani, N. et al. G Protein Activation Occurs via a Largely Universal Mechanism. J. Phys. Chem. B 128, 3554–3562 (2024).

4. Doshi, U., McGowan, L. C., Ladani, S. T. & Hamelberg, D. Resolving the complex role of enzyme conformational dynamics in catalytic function. Proc. Natl. Acad. Sci. U.S.A. 109, 5699–5704 (2012).

5. Hersey, A. N., Kay, V. E., Lee, S., Realff, M. J. & Wilson, C. J. Engineering allosteric transcription factors guided by the LacI topology. Cell Systems 14, 645–655 (2023).

6. Walker, A. S., Russ, W. P., Ranganathan, R. & Schepartz, A. RNA sectors and allosteric function within the ribosome. Proc. Natl. Acad. Sci. U.S.A. 117, 19879–19887 (2020).

7. Han, B., Salituro, F. G. & Blanco, M.-J. Impact of Allosteric Modulation in Drug Discovery: Innovation in Emerging Chemical Modalities. ACS Med. Chem. Lett. 11, 1810–1819 (2020).

8. Lemmon, M. A. Membrane recognition by phospholipid-binding domains. Nat Rev Mol Cell Biol 9, 99–111 (2008).

9. Kusumi, A. et al. Dynamic Organizing Principles of the Plasma Membrane that Regulate Signal Transduction: Commemorating the Fortieth Anniversary of Singer and Nicolson’s Fluid-Mosaic Model. Annual Review of Cell and Developmental Biology 28, 215–250 (2012).

10. Lu, S., et al. Ras Conformational Ensembles, Allostery, and Signaling. Chem. Rev. 116, 6607– 6665 (2016).

11. Gorfe, A. Mechanisms of Allostery and Membrane Attachment in Ras GTPases: Implications for Anti-Cancer Drug Discovery. CMC 17, 1–9 (2010).

12. Bento, C. F. et al. Mammalian Autophagy: How Does It Work? Annu. Rev. Biochem. 85, 685– 713 (2016).

13. Itakura, E. & Mizushima, N. Characterization of autophagosome formation site by a hierarchical analysis of mammalian Atg proteins. Autophagy 6, 764–776 (2010).

14. Vargas, J. N. S., Hamasaki, M., Kawabata, T., Youle, R. J. & Yoshimori, T. The mechanisms and roles of selective autophagy in mammals. Nat Rev Mol Cell Biol 24, 167–185 (2023).

15. Mizushima, N., Yoshimori, T. & Ohsumi, Y. The Role of Atg Proteins in Autophagosome Formation. Annu. Rev. Cell Dev. Biol. 27, 107–132 (2011).

16. Kabeya, Y. LC3, a mammalian homologue of yeast Apg8p, is localized in autophagosome membranes after processing. The EMBO Journal 19, 5720–5728 (2000).

17. Thukral, L., Sengupta, D., Ramkumar, A., Murthy, D., Agrawal, N. & Gokhale, R. S. The Molecular Mechanism Underlying Recruitment and Insertion of Lipid-Anchored LC3 Protein into Membranes. Biophysical Journal 109, 2067–2078 (2015).

18. Johansen, T. & Lamark, T. Selective autophagy mediated by autophagic adapter proteins. Autophagy 7, 279–296 (2011).

19. Birgisdottir, Å. B., Lamark, T. & Johansen, T. The LIR motif – crucial for selective autophagy. Journal of Cell Science 126, 3237–3247 (2013).

20. Noda, N. N. et al. Structural basis of target recognition by Atg8/LC3 during selective autophagy. Genes to Cells 13, 1211–1218 (2008).

21. Ichimura, Y. et al. Structural Basis for Sorting Mechanism of p62 in Selective Autophagy. Journal of Biological Chemistry 283, 22847–22857 (2008).

22. Rogov, V. V. et al. Structural basis for phosphorylation-triggered autophagic clearance of Salmonella. Biochemical Journal 454, 459–466 (2013).

23. Rozenknop, A. et al. Characterization of the Interaction of GABARAPL-1 with the LIR Motif of NBR1. Journal of Molecular Biology 410, 477–487 (2011).

24. Wild, P., McEwan, D. G. & Dikic, I. The LC3 interactome at a glance. Journal of Cell Science jcs.140426 (2014) doi:10.1242/jcs.140426.

25. Behrends, C., Sowa, M. E., Gygi, S. P. & Harper, J. W. Network organization of the human autophagy system. Nature 466, 68–76 (2010).

26. Klionsky, D. J. et al. Guidelines for the use and interpretation of assays for monitoring autophagy (4th edition)1. Autophagy 17, 1–382 (2021).

27. Zhang, W. et al. Autophagosome membrane expansion is mediated by the N-terminus and cis-membrane association of human ATG8s. eLife 12, e89185 (2023).

28. Fracchiolla, D., Chang, C., Hurley, J. H. & Martens, S. A PI3K-WIPI2 positive feedback loop allosterically activates LC3 lipidation in autophagy. Journal of Cell Biology 219, e201912098 (2020).

29. Jatana, N., Ascher, D. B., Pires, D. E. V., Gokhale, R. S. & Thukral, L. Human LC3 and GABARAP subfamily members achieve functional specificity via specific structural modulations. Autophagy 16, 239–255 (2020).

30. Noda, N. N., Ohsumi, Y. & Inagaki, F. Atg8-family interacting motif crucial for selective autophagy. FEBS Letters 584, 1379–1385 (2010).

31. Sun, X., Singh, S., Blumer, K. J. & Bowman, G. R. Simulation of spontaneous G protein activation reveals a new intermediate driving GDP unbinding. eLife 7, e38465 (2018).

32. Bavro, V. N. et al. Crystal structure of the GABAA -receptor-associated protein, GABARAP. EMBO Reports 3, 183–189 (2002).

33. Suzuki, H. et al. Structural Basis of the Autophagy-Related LC3/Atg13 LIR Complex: Recognition and Interaction Mechanism. Structure 22, 47–58 (2014).

34. Atkinson, J. M. et al. Time-resolved FRET and NMR analyses reveal selective binding of peptides containing the LC3-interacting region to ATG8 family proteins. Journal of Biological Chemistry 294, 14033–14042 (2019).

35. Singh, S. & Bowman, G. R. Quantifying Allosteric Communication via Both Concerted Structural Changes and Conformational Disorder with CARDS. J. Chem. Theory Comput. 13, 1509–1517 (2017).

36. Khaminets, A. et al. Regulation of endoplasmic reticulum turnover by selective autophagy. Nature 522, 354–358 (2015).

37. Das, D. et al. VPS4A is the selective receptor for lipophagy in mice and humans. Molecular Cell 84, 4436–4453.e8 (2024).

38. Patrick, J. W., et al. Allostery revealed within lipid binding events to membrane proteins. Proc. Natl. Acad. Sci. U.S.A. 115, 2976–2981 (2018).

39. Cong, X., Liu, Y., Liu, W., Liang, X. & Laganowsky, A. Allosteric modulation of protein-protein interactions by individual lipid binding events. Nat Commun 8, 2203 (2017).

40. Lee, J. et al. CHARMM-GUI Input Generator for NAMD, GROMACS, AMBER, OpenMM, and CHARMM/OpenMM Simulations Using the CHARMM36 Additive Force Field. J. Chem. Theory Comput. 12, 405–413 (2016).

41. Huang, J. et al. CHARMM36m: an improved force field for folded and intrinsically disordered proteins. Nat Methods 14, 71–73 (2017).

42. Abraham, M. J. et al. GROMACS: High performance molecular simulations through multi-level parallelism from laptops to supercomputers. SoftwareX 1–2, 19–25 (2015).

43. Essmann, U. et al. A smooth particle mesh Ewald method. The Journal of Chemical Physics 103, 8577–8593 (1995).

44. Hess, B., Bekker, H., Berendsen, H. J. C. & Fraaije, J. G. E. M. LINCS: A linear constraint solver for molecular simulations. J. Comput. Chem. 18, 1463–1472 (1997).

45. Nosé, S. A molecular dynamics method for simulations in the canonical ensemble. Molecular Physics 52, 255–268 (1984).

46. Parrinello, M. & Rahman, A. Polymorphic transitions in single crystals: A new molecular dynamics method. Journal of Applied Physics 52, 7182–7190 (1981).

47. Laurent, B. et al. Epock: rapid analysis of protein pocket dynamics. Bioinformatics 31, 1478– 1480 (2015).

48. Wagner, J. R. et al. POVME 3.0: Software for Mapping Binding Pocket Flexibility. J. Chem. Theory Comput. 13, 4584–4592 (2017).

49. Kraskov, A., Stögbauer, H. & Grassberger, P. Estimating mutual information. Phys. Rev. E 69, 066138 (2004).

50. Pettersen, E. F. et al. UCSF Chimera—A visualization system for exploratory research and analysis. J Comput Chem 25, 1605–1612 (2004).

51. Shapovalov, M. V. & Dunbrack, R. L. A Smoothed Backbone-Dependent Rotamer Library for Proteins Derived from Adaptive Kernel Density Estimates and Regressions. Structure 19, 844– 858 (2011).

52. Michaud-Agrawal, N., Denning, E. J., Woolf, T. B. & Beckstein, O. MDAnalysis: A toolkit for the analysis of molecular dynamics simulations. J Comput Chem 32, 2319–2327 (2011).

53. Humphrey, W., Dalke, A. & Schulten, K. VMD: Visual molecular dynamics. Journal of Molecular Graphics 14, 33–38 (1996).

54. Pettersen, E. F. et al. UCSF CHIMERAX : Structure visualization for researchers, educators, and developers. Protein Science 30, 70–82 (2021).

55. PyMOL. Schrödinger, LLC and Warren DeLano (2020).

56. Hunter, J. D. Matplotlib: A 2D Graphics Environment. Comput. Sci. Eng. 9, 90–95 (2007).

57. Waskom, M. seaborn: statistical data visualization. JOSS 6, 3021 (2021).

58. Shimoyama, Y. pyCirclize: Circular visualization in Python. GitHub (2022).

59. Otwinowski, Z. & Minor, W. [20] Processing of X-ray diffraction data collected in oscillation mode. in Methods in Enzymology vol. 276 307–326 (Elsevier, 1997).

60. McCoy, A. J., et al. Phaser crystallographic software. J Appl Crystallogr 40, 658–674 (2007).

61. Murshudov, G. N., Vagin, A. A. & Dodson, E. J. Refinement of Macromolecular Structures by the Maximum-Likelihood Method. Acta Crystallogr D Biol Crystallogr 53, 240–255 (1997).

62. Emsley, P., Lohkamp, B., Scott, W. G. & Cowtan, K. Features and development of Coot. Acta Crystallogr D Biol Crystallogr 66, 486–501 (2010).

63. Chen, V. B. et al. MolProbity : all-atom structure validation for macromolecular crystallography. Acta Crystallogr D Biol Crystallogr 66, 12–21 (2010).

64. Schindelin, J. et al. Fiji: an open-source platform for biological-image analysis. Nat Methods 9, 676–682 (2012).

65. Neha, Castin, J., Fatihi, S., Gahlot, D., Arun, A. & Thukral, L. Autophagy3D: a comprehensive autophagy structure database. Database 2024, (2024).

66. Malhotra, N. et al. AI-based AlphaFold2 significantly expands the structural space of the autophagy pathway. Autophagy 19, 3201–3220 (2023).

67. Katoh, K. & Standley, D. M. MAFFT Multiple Sequence Alignment Software Version 7: Improvements in performance and usability. Molecular Biology and Evolution 30, 772–780 (2013).

68. Livingstone, C. D. & Barton, G. J. Protein sequence alignments: a strategy for the hierarchical analysis of residue conservation. Computer Applications in the Biosciences 9, 745–756 (1993).

