## Supplementary information for "A novel lipid-triggered allosteric site modulates LC3-LIR receptor binding activity"

This PDF file includes:

Supplementary Notes

Supplementary Methods

Supplementary Figures 1 to 24

Supplementary Tables 1 to 4

Supplementary Videos 1 to 3

**Supplementary Note 1: Nomenclature used in the paper**

*“LC3”* in this paper refers to the LC3B, the Human ortholog of ATG8 protein.

i) **LC3 States:**
   a. **LC3^M-^:** LC3^M-^ signifies the cytosolic state of LC3 in soluble form (known as proLC3).
   b. **LC3^M+R-^:** LC3^M+R-^ denotes the membrane-inserted LC3 (M+) in the apo state (R-). The 120^th^ glycine residue is covalently linked to phosphatidyl ethanolamine (PE) lipid (known as LC3-II).
   c. **LC3^M+R+^:** LC3^M+R+^ denotes the membrane-inserted LC3 (M+) with the p62 LIR peptide bound to the hydrophobic pockets (HP1, HP2). p62-LIR is used as the key peptide to study the receptor-bound state on the membrane.
ii) **Regions of LC3:**
    a. **MIS:** Membrane interacting site (MIS) refers to the residues of LC3 whose lipid interaction propensity is greater than 0.5 across all independent replica of LC3^M+^ simulations. The site is primarily comprised of residues in the N-terminal (residues 1 to 6, and 9 to 12), Loop3 (residues 37 to 43, and 45 to 46) and C-terminal (residue 119).
    b. **BP:** Binding pockets (BP) refer to canonical binding pockets- hydrophobic pockets 1 and 2 (HP1: residues 19, 23, 32, 34, 51, 53, and 108, HP2: residues 35, 52, 55, 58, 63, 66, 67, and 70).
    c. **REST:** REST defines the remainder residues of the LC3 protein that do not belong to either MIS or BP.
    d. **Allosteric site (AS):** The allosteric site (AS) defines the specific residues in the rest of the protein whose communication strength is greater than 0.65 (L6-facing α3: residues 60, 61, 64, and 68; Loop 5: residues 73, 78, 79; β3: residues 80 to 82; Loop 6: residues 87 to 93).
iii) **Functional states:**
     a. **Active:** The LC3 is considered functionally “active”, given there are two concomitant changes: i) stable allosteric site, ii) stable binding pockets primed for receptor binding.
     b. **Inactive:** The LC3 is considered functionally “inactive” when i) the allosteric site exhibits instability and ii) the binding pockets are unstable.
iv) **Mutants:**
    a. **LC3^AS_Mutant1^:** LC3^AS_Mutant1^ refers to the active mutant I64K (α3)- V89D (L6)- V91F (L6). This mutant is considered to be active because it introduces stability to the allosteric site, which leads to increased receptor binding affinity.
    b. **LC3^AS_Mutant2^:** LC3^AS_Mutant2^ refers to the inactive mutant I64D (α3)- V89P (L6)- V91D (L6). This is considered inactive as it destabilises the allosteric site, eventually resulting in decreased receptor binding affinity.

**Supplementary Methods**

**Assigning orderness states to the residue dihedral dynamics as a function of time:**

From the time evolution of dihedral rotamer states, we applied the transition indicator function. If a residue adopts a rotamer r_x_ at time t and maintains the same rotamer state at t+1, the transition indicator will assign this as a non-transitioning event. The transition event is marked otherwise. Thus, we reduce the rotamer information to a simple discrete transition state (*“transition”* when rotamer state changes from t to t+1; *“no transition”* otherwise) as a function of time. For each time point, we applied the k-neighbourhood binning scheme to group the frames into distinct time bins. For example, the time bin for t = 5 ns will include 2 ns, 3 ns, 4 ns, 6 ns, 7 ns, and 8 ns (when k = 3). For each time bin of transition states, we computed Shannon entropy, and if the entropy is greater than 0.5, we assign each time point in the bin as *“disordered”*. Otherwise, they are assigned *“ordered”*. This gives the orderness states across time.

**Computing mutual information between dihedral-dihedral variables:**

As the dihedral values are continuous in nature, we estimated mutual information (MI) with the nearest neighborhood method as implemented in the Scikit Python library. For two dihedral value sets as a function of time $d_{i}, d_{j}$, the MI is calculated as follows

$$MI\left( d_{i},d_{j} \right)=\psi\left( k \right)+\psi\left( N \right)-\left\langle\psi\left( n_{dix}+1 \right) \right\rangle-\left\langle\psi\left( n_{djy}+1 \right) \right\rangle$$

where,

- $\psi$ is the digamma function
- $k=3$ is the neighborhood parameter
- $N$ represents the total number of samples
- $n_{dix}$ and $n_{djy}$ denote the number of neighbors within the k^th^ neighborhood of a point $p\left( x,y \right)$ along the x and y axes, respectively
- $\left\langle. \right\rangle$ represents the mean

Neighborhoods are determined as follows

- For each point $p\left( x,y \right)$, the Chebyshev distance to all other points is computed
- The third smallest distance (since k=3) is considered as the neighborhood radius (k3 radius)
- A point is considered to be in a neighborhood along the x or y axis if the Manhattan distance of the point $p\left( x,y \right)$ is less than the k3 radius
- The number of points that are falling in the neighborhood along x-axis ($n_{dix}$) and y-axis ($n_{djy}$) is then obtained
- This is iterated for all the points

**Computing mutual information between rotamer-rotamer or orderness state-orderness state variables:**

For rotamer or orderness states, we computed MI with the following equation for two discrete variable sets.

$$MI\left( s_{i}, s_{j} \right)=\sum_{x\in s_{i}} \sum_{y\in s_{j}} p\left( x,y \right) log\left( \frac{p\left( x,y \right)}{p\left( x \right)p\left( y \right)} \right)$$

where

- $s_{i}, s_{j}$ are the rotamer or orderness states of residues $r_{i}, r_{j}$ across time
- $p\left( x,y \right)$ represents the joint probability of $s_{i}$ and $s_{j}$ to exist in $x$ and $y$ states, respectively
- $p\left( x \right)$ denotes the marginal probability of $s_{i}$ existing in $x$ state
- $p\left( y \right)$ denotes the marginal probability of $s_{j}$ existing in $y$ state

**Computing the relative contribution of region-specific coupling to the overall net communication:**

To quantify how each LC3 region contributes to the overall net communication coupling, we calculated the relative contribution. We computed the mean residue-residue coupling between residues in a given region $Reg_{a}$ and all remaining residues $Reg_{NOT a}$. This was normalized by the mean coupling over all residue pairs in the protein. Finally, we scaled the resulting value by the weight, which is defined as the fraction of residue pairs belonging to $Reg_{a}, Reg_{NOT a}$ relative to the total number of residue pairs.

$$Contributio{n_{Reg}}_{a}=\frac{Mean Couplin{g_{Reg}}_{a}}{Mean overall coupling}\times weight$$

where $Mean Couplin{g_{Reg}}_{a}$is given by,

$$Mean Couplin{g_{Reg}}_{a}=\frac{1}{n\left( Reg_{a},Reg_{NOT a} pairs \right)}\sum_{\begin{aligned} i\in Reg_{a}, \\ j\in Reg_{NOTa} \end{aligned}} Coupling_{i,j}$$

And $Mean overall coupling$ can be written as follows,

$$Mean overall coupling=\frac{1}{n(overall pairs)}\sum_{\begin{aligned} i\in All residues, \\ j\in All residues, \\ i\neq j \end{aligned}} Coupling_{i,j}$$

Weight is defined as follows,

$$weight=\frac{n(Reg_{a},Reg_{NOT a} pairs)}{n(overall pairs)}$$

**Computing distances between charged residues and membrane:**

The charged atom groups of MIS residues were chosen (NH2 for arginine, NZ for lysine and OE2 for glutamic acid). The minimum distance between the charged group and the phosphate head in the upper leaflet was calculated as a function of time.

**Mean distances of HP1 and HP2 pockets:**

To track the impact of residue side chain movements on the pocket volume, we calculated distances between terminal atoms of each pocket residue and the pocket center of geometry (COG). For the HP1 pocket, CG, CD, CG, CD, NZ, CD1, and CZ atoms were selected for D19, I23, P32, I34, K51, L53, and F108, respectively. For HP2 pocket, CD, CZ, CG2, CG, CG2, CD1, CD, CD, CD, CD, and NH1 were taken for I35, F52, V54, P55, V58, L63, I66, I67, and R70 residues respectively. The correlation between the distance and the volume was computed with the Pearson correlation coefficient. We define the mean distance of HP1 and HP2 (intra-pocket distance) at a given time as follows.

$$Mean Distance_{HP1}=\frac{\sum_{R_{i}} Distance_{R_{i}-HP1 COG}}{n\left( R \right)} \left[ where R=\{D19, I23, K51\}; n\left( R \right)=3 \right]$$

$$Mean Distance_{HP2}=\frac{\sum_{R_{i}} Distance_{R_{i}-HP2 COG}}{n\left( R \right)} \left[ where R=\{I35, F52,R70,I66\}; n\left( R \right)=4 \right]$$

Based on the importance of residue contribution to the pocket volume, we chose these specific residues for monitoring the pockets. We define net mean distance as the sum of mean distances calculated for HP1 and HP2 pockets as a function of time.

**Allosteric site estimation calculations:**

For monitoring the dynamical state of the allosteric site in different LC3 states and mutant candidates, we computed the minimum distance between residue positions 64 (of α3) and 89 (of L6) due to their consistent interaction when the allosteric site is stable. We define this as the allosteric site probing distance.

**Estimating the convergence of trajectories:**

i) **Standard error of mean RMSD at different trajectory time-blocks:** The RMSD values were divided into 200ns time blocks. For each time block, the standard error of the mean was computed. Low standard error signifies less variability in conformations across time and more convergence.

$$S.E_{mean RMSD}=\frac{\sigma}{\sqrt{n}}$$

$\sigma$ represents the standard deviation of RMSD in a given time-block, and $n$ represents the number of values in a given time-block.

**Extracting representative structures from the trajectory:**

The GROMOS algorithm was applied to cluster the frames of the simulation trajectory using *gmx cluster*, with an RMSD cut-off set at 0.2 nm. The representative structure of a trajectory was taken from the most populated cluster.

**Solvent accessible surface area calculations:**

The solvent accessible surface area of residues was computed using the Shrake Tupley algorithm implemented in the mdtraj Python library. The default setting was considered (probe radius= 0.14 nm and number of sphere points=960). The overall (net) pocket SASA was obtained by summing the individual SASA values of the corresponding pocket residues.

**Supplementary Figures**

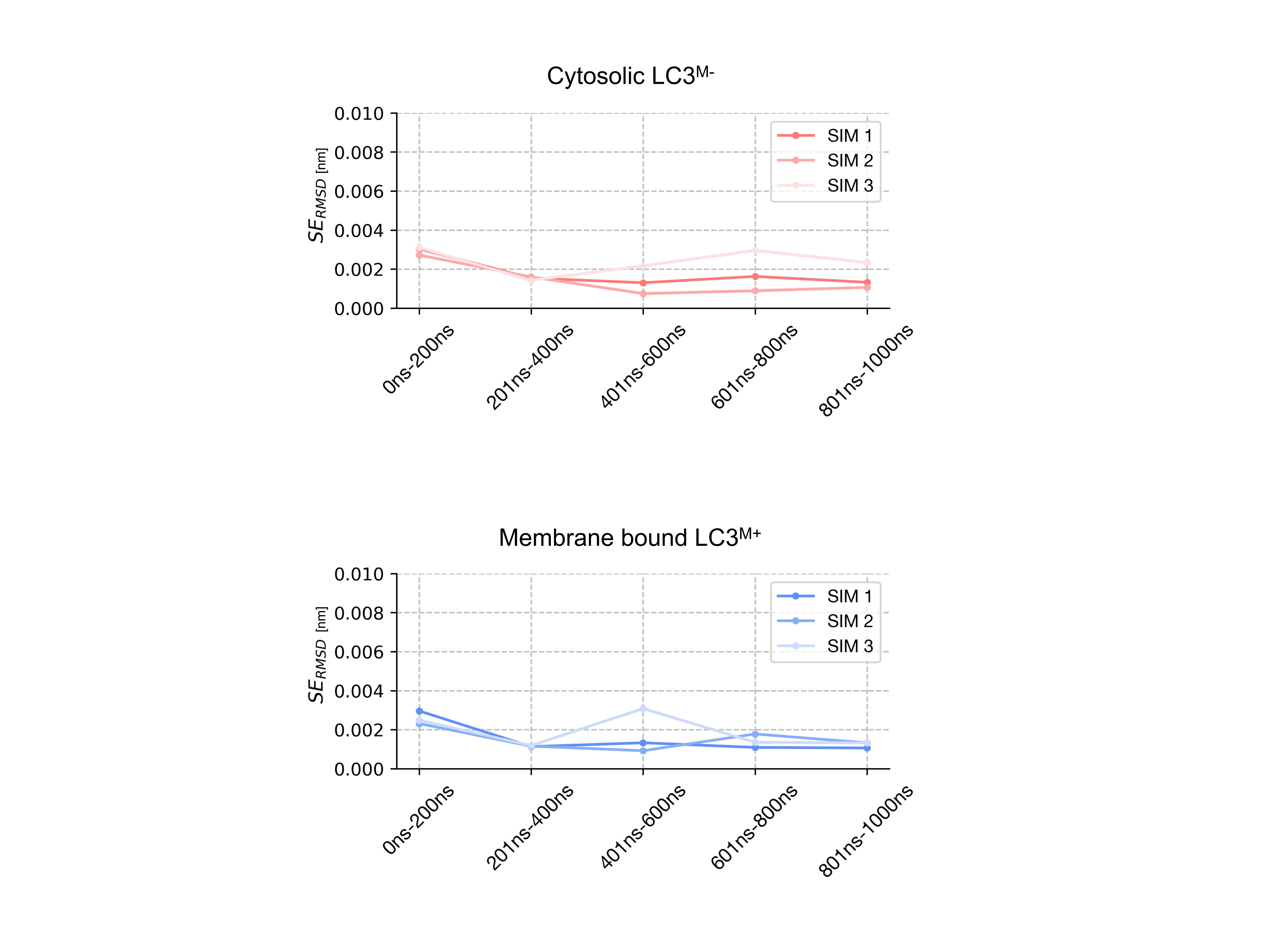

**Supplementary Figure 1. Standard error of mean RMSD in LC3^M˗^ and LC3^M+^** **Simulations**. The line plots highlight the standard error of mean RMSD across different time blocks to represent convergence of the trajectories, where red and blue lines represent cytosolic LC3^M˗^ and membrane-bound LC3^M+^ states, respectively.

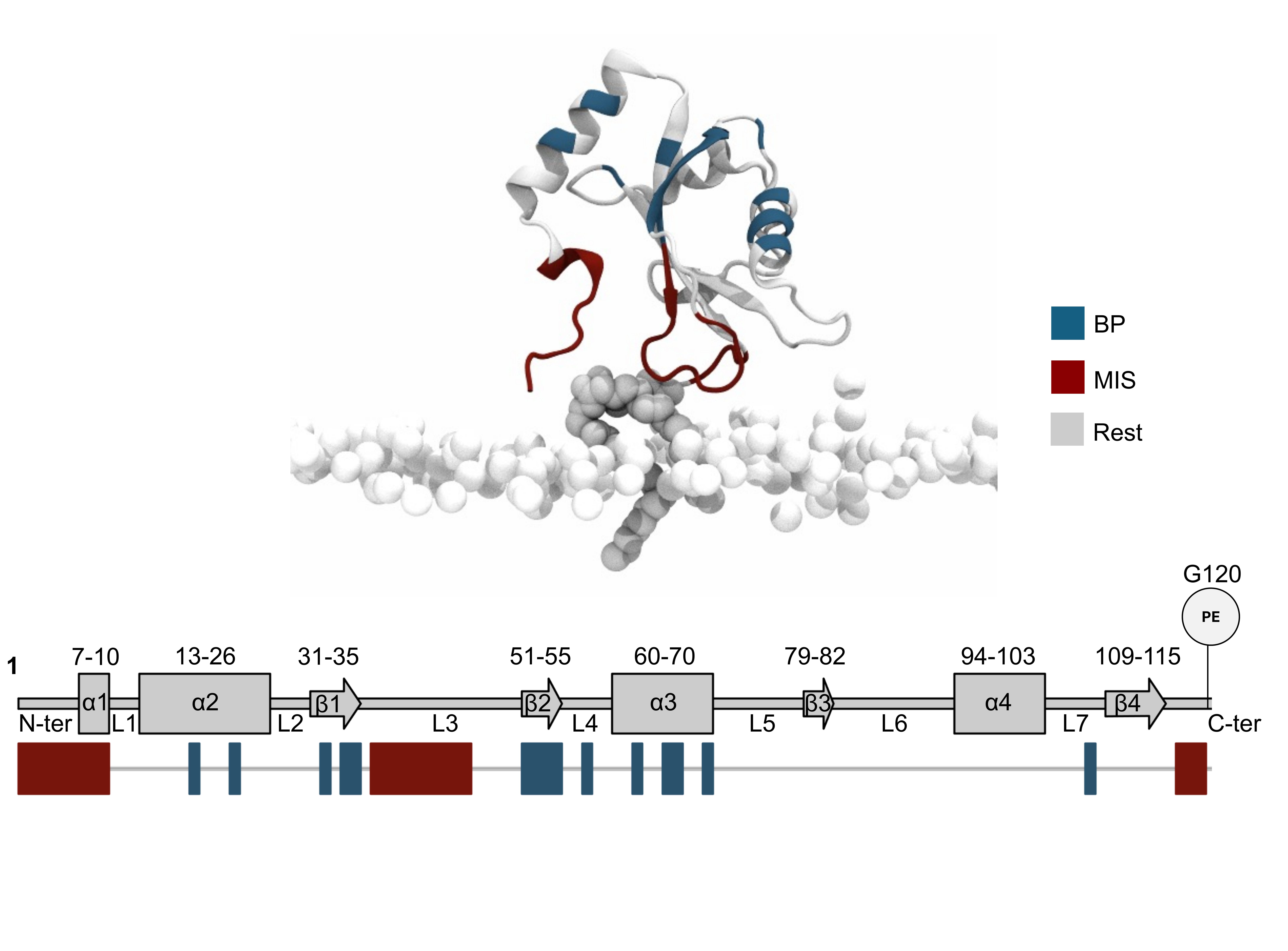

**Supplementary Figure 2. LC3 structural architecture of LC3.** The cartoon representation of LC3 structure highlights the local secondary structural segments and residues that constitutes these regions. The functional motifs involving hydrophobic pockets, including binding pockets (HP1 and HP2), and membrane interacting site, are colored as teal and maroon, respectively. The PE tail and phosphate group of ER membrane lipids are shown as white Vanderwal spheres. The lower panel presents a schematic of the secondary structure map of LC3, with sequence annotations marked on it.

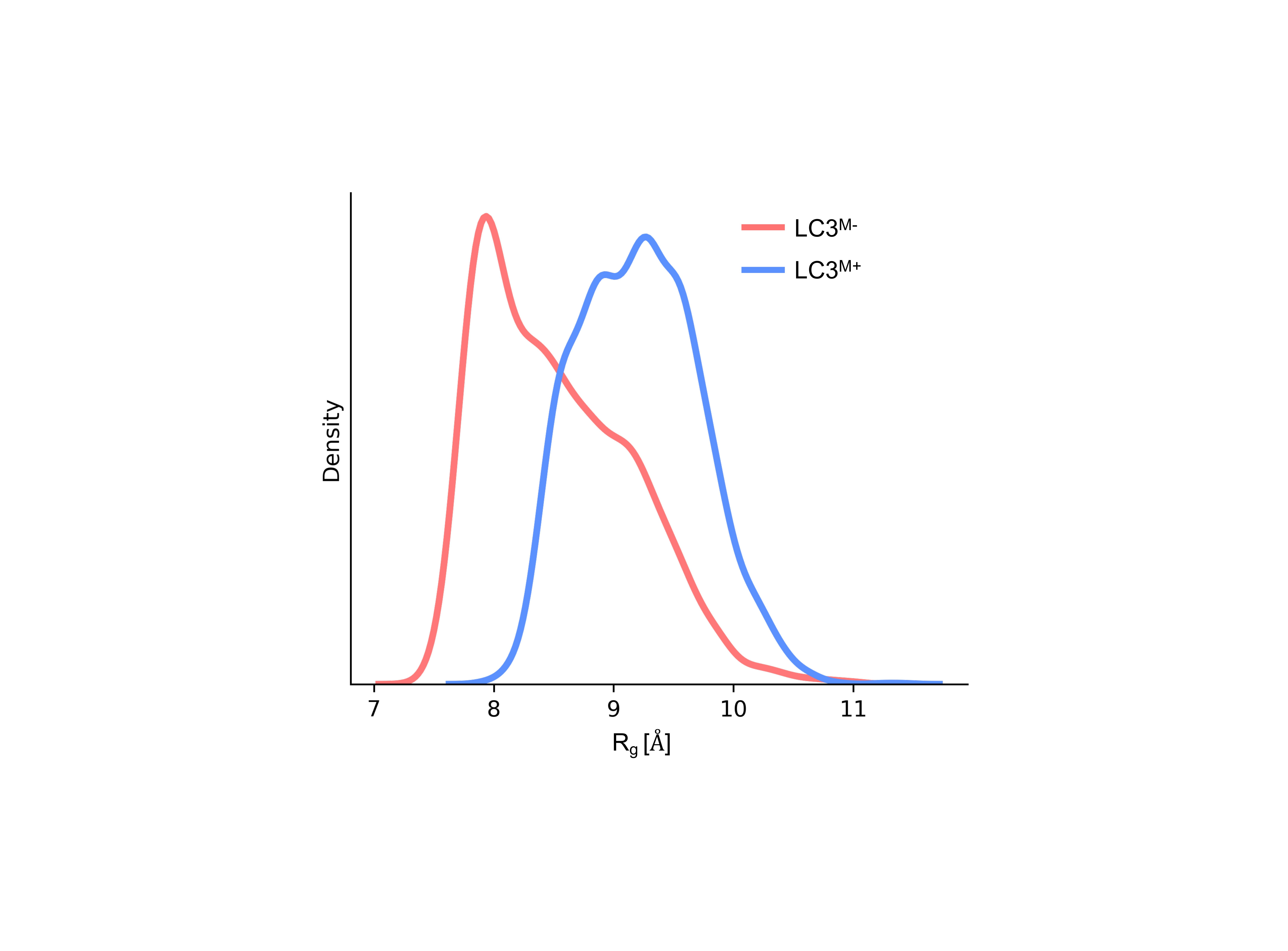

**Supplementary Figure 3. The lipid interactions alter the compactness of the MIS region.** The distribution of radius of gyration measured for N-terminal (residues 1 to 10) and L3 residues (residues 37 to 41) of MIS is shown for cytosolic LC3^M˗^ and membrane-bound receptor state LC3^M+^.

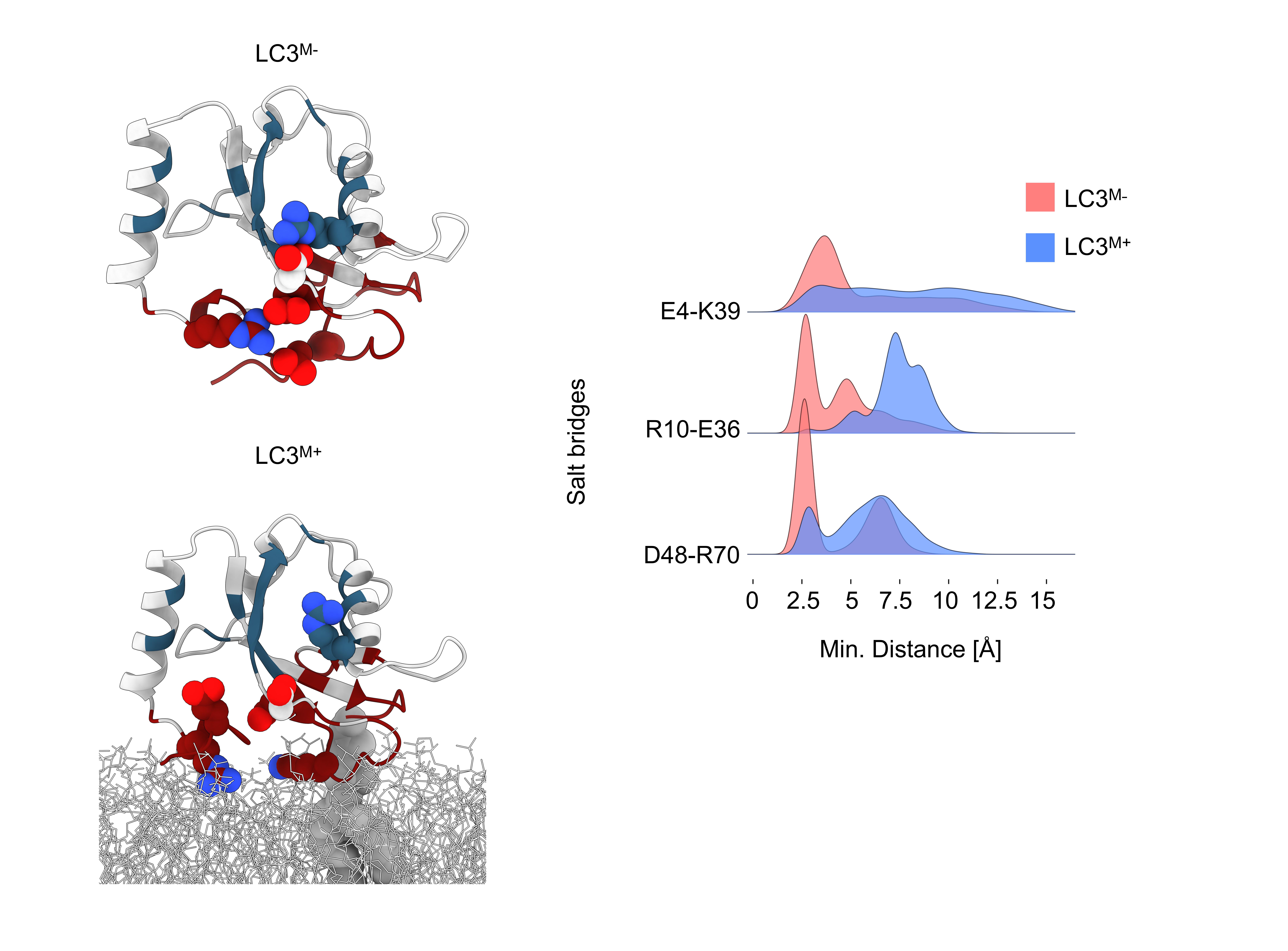

**Supplementary Figure 4: Disruption of conserved salt bridges in the membrane-bound state.** Representative structural snapshots (left) show the presence of conserved salt bridges (E4-K39, R10-E36, and D48-R70) in the cytosolic LC3^M˗^ (top) compared to their loss in membrane association LC3^M+^ (bottom). The corresponding distance distribution (right) highlights that in LC3^M˗^ state, the three salt-bridges are stable in all trajectories whereas in the LC3^M+^ state, they are disrupted and display broader distance distribution.

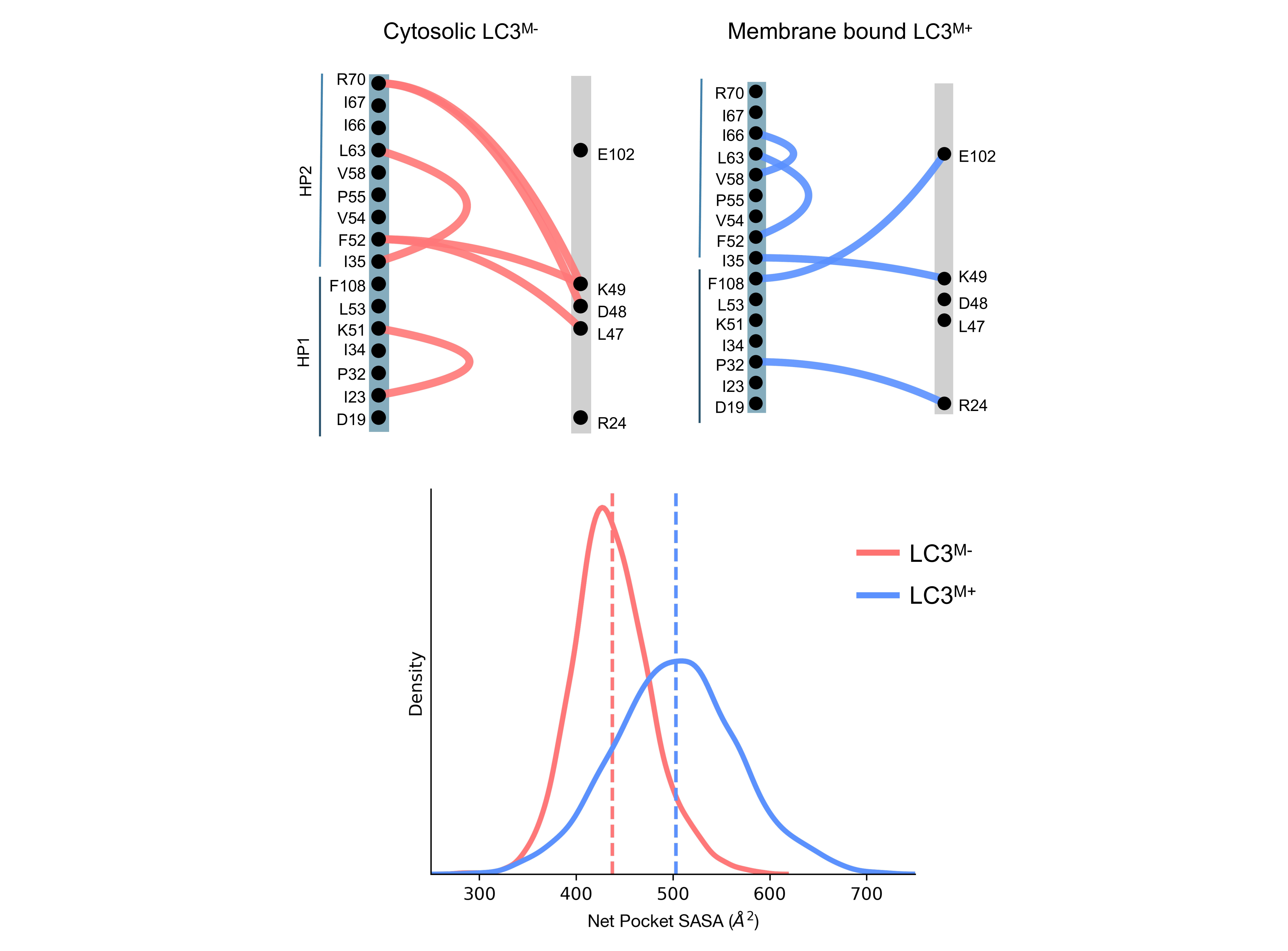

**Supplementary Figure 5. Membrane binding reorganizes binding pockets contacts, leading to increased exposure of the hydrophobic binding pockets (BP).** The alluvial diagrams (top) represent the unique contacts observed in cytosolic LC3^M˗^ and membrane-bound state LC3^M+,^ respectively. The unique contacts of the cytosolic state (LC3^M-^) are obtained by comparing with the apo LC3-II (LC3^M+^) and vice versa. The density plot (bottom) highlights the distribution of net solvent-exposed surface area (SASA) of HP1 and HP2 binding pockets in LC3^M˗^ and LC3^M+^ states. The dashed lines represent the mean of the net pocket SASA.

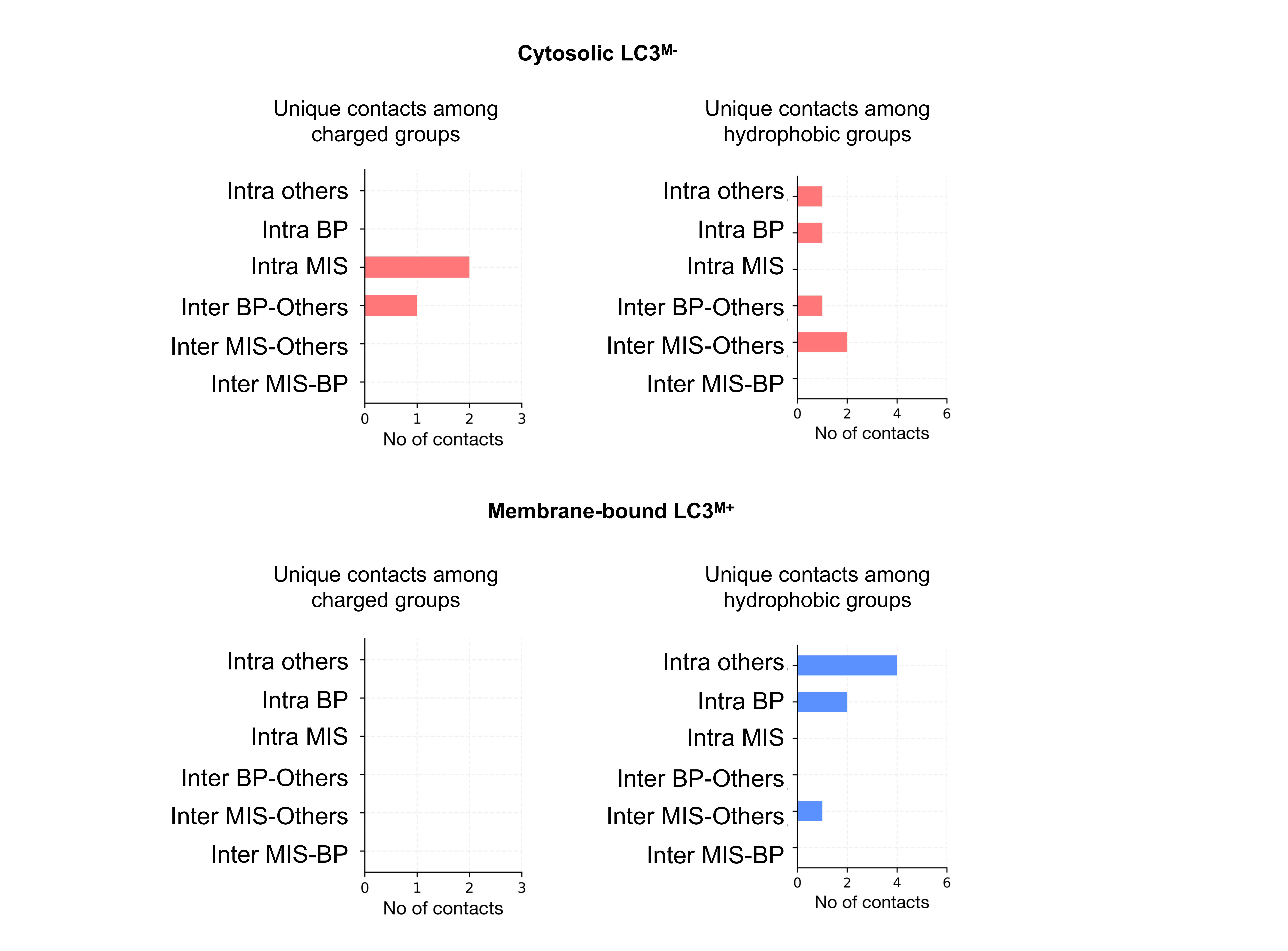

**Supplementary Figure 6. Decomposition of intra-protein contacts**. The vertical bar plots highlight the number of unique charged and hydrophobic contacts between different functional regions of the LC3 protein in two states, namely, cytosolic LC3^M-^ and membrane-bound LC3^M+^ states. This highlights how contacts within LC3 are reorganized across different functional regions due to membrane interaction.

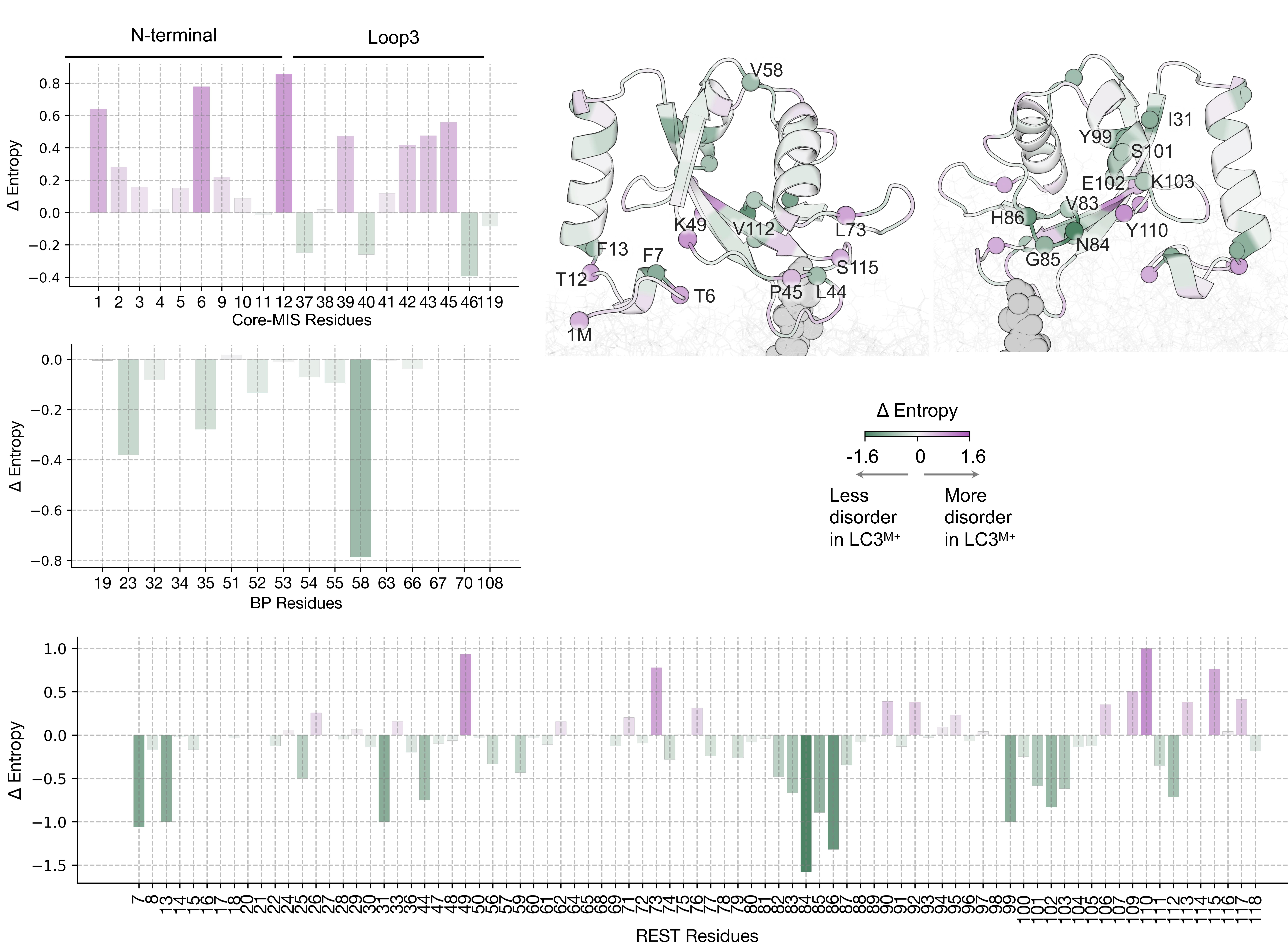

**Supplementary Figure 7. Dihedral entropy differences comparing LC3^M-^ and LC3^M+^.** Δ Entropy (Entropy LC3^M+^ - Entropy LC3^M˗^) is plotted for each region of LC3. The positive value indicates increased disorder in LC3^M+^ state, and the negative value indicates a decrease in disorder. The values are mapped on the representative structure of LC3^M+^. Residues with $\left| \Delta Entropy \right|>0.5$ are marked as spheres and labeled.

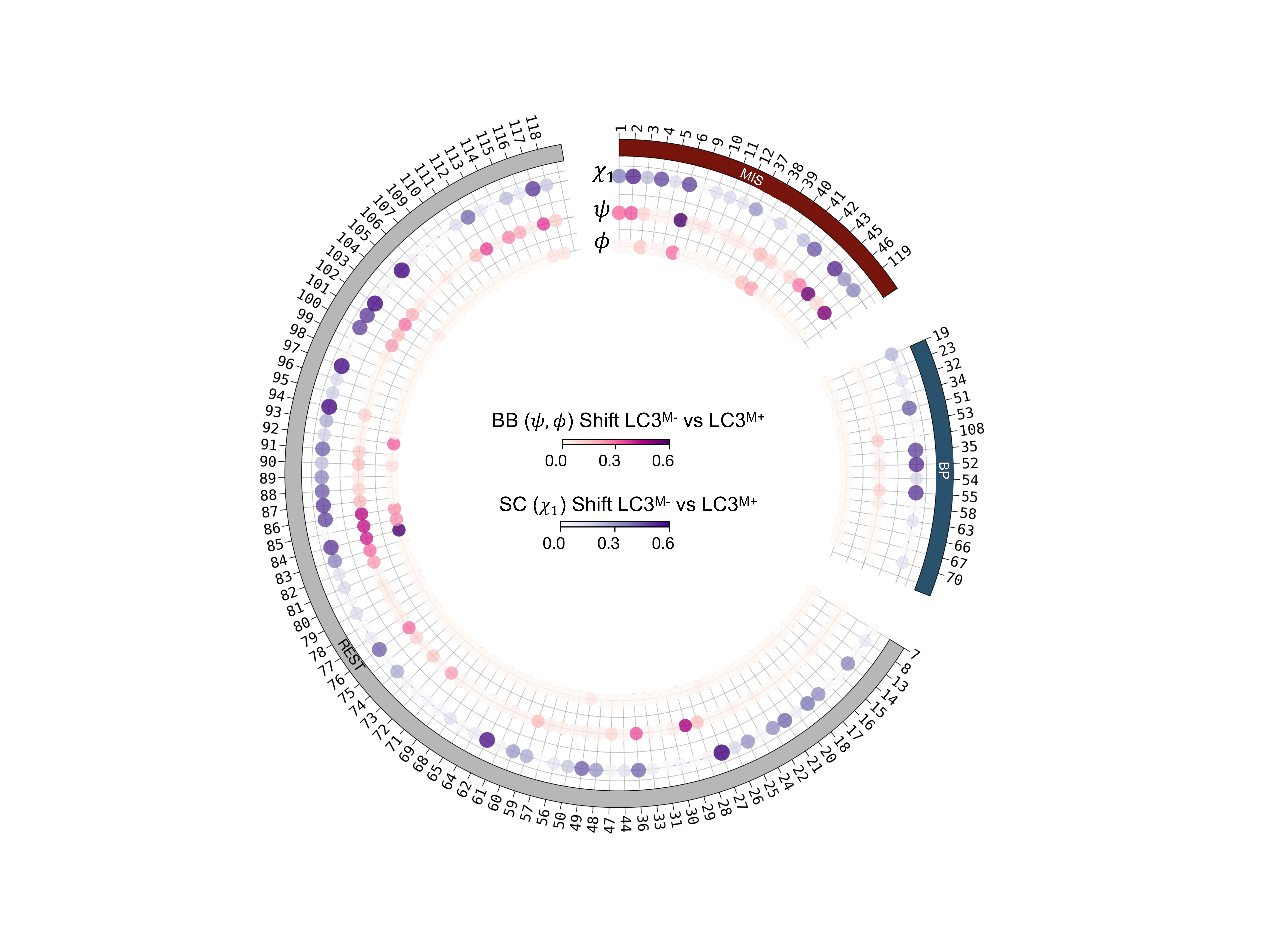

**Supplementary Figure 8. LC3 undergoes dihedral conformational changes on the membrane.** Rotamer shifts in the backbone (φ, ψ) (BB) and sidechain (χ1) (SC) were calculated by comparing LC3^M-^ and LC3^M+^ and are visualized as a circular scatter map. The size and color intensity of each point correspond to the shift. Residues are grouped according to their respective structural regions within LC3.

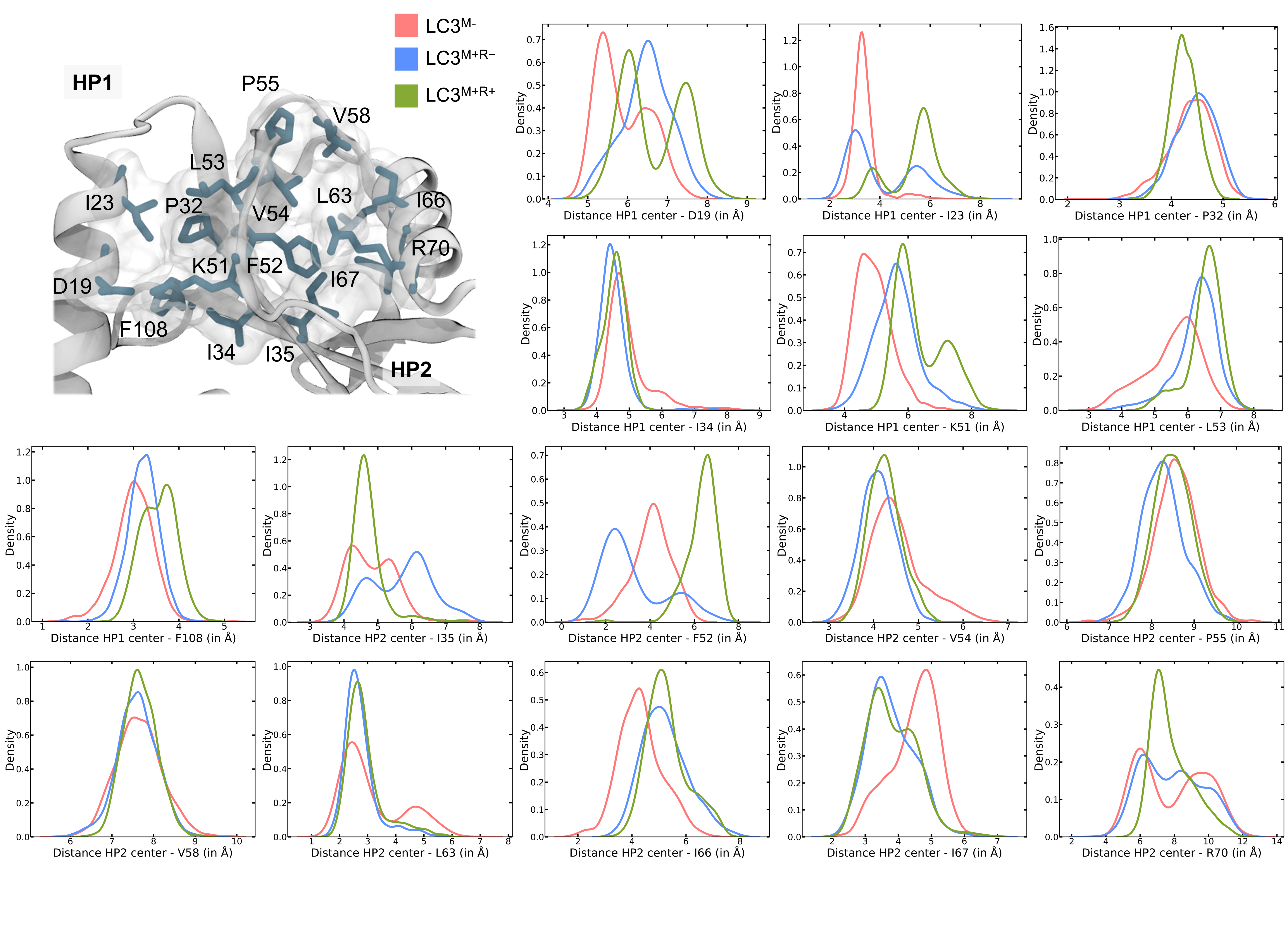

**Supplementary Figure 9. The distance of residues in BP from the pocket center changes upon membrane binding.** The structural snapshot displays the residues lining the receptor binding pockets of LC3 in stick representation. The density plots display the density of the distance measured between the terminal heavy atom of the BP residues sidechain and the pocket center in three different LC3 states. The colors represent the three states of LC3.

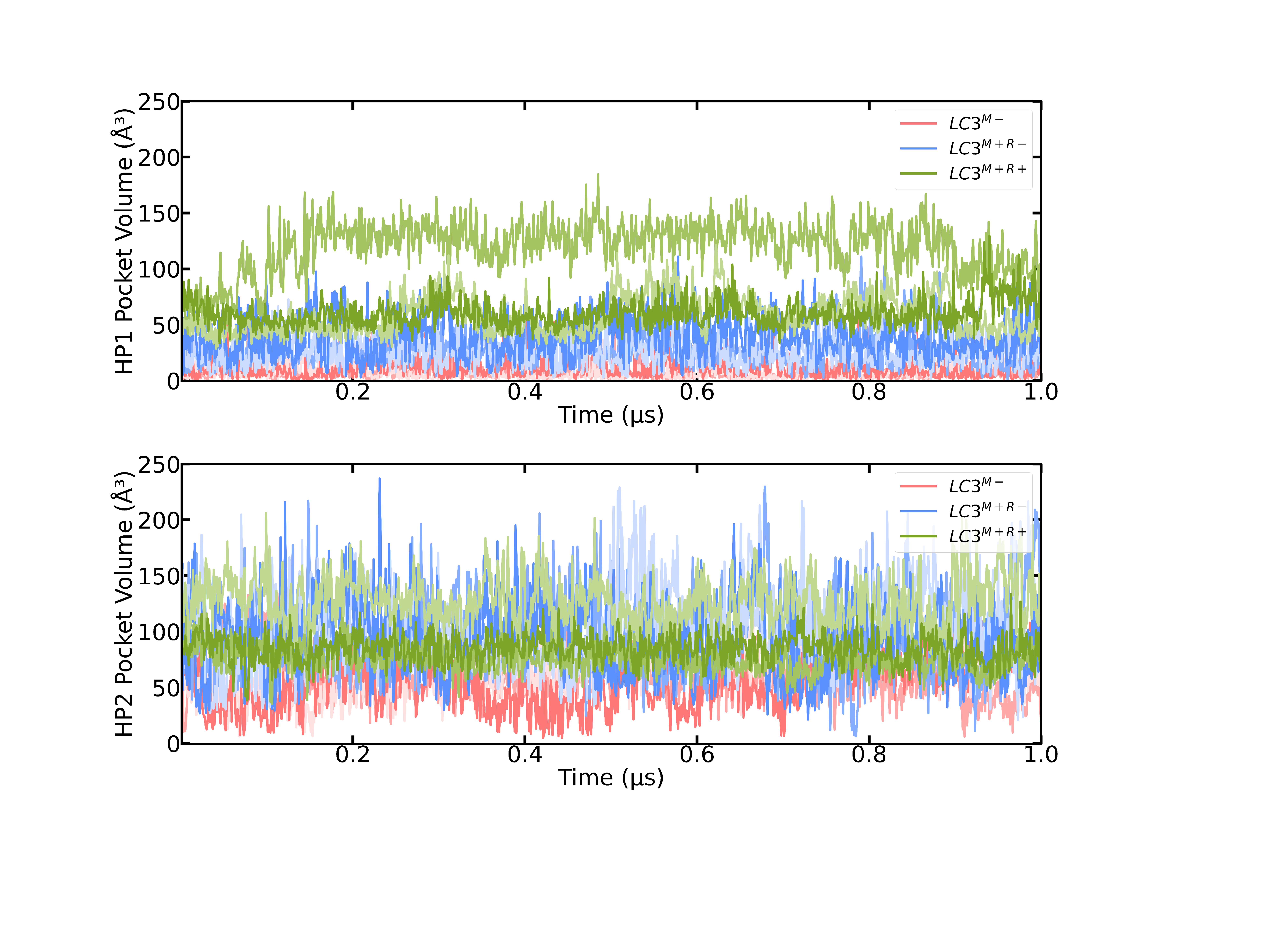

**Supplementary Figure 10. Time evolution of HP1 and HP2 volume for three LC3 states.** The line-plot displays the time evolution of HP1 and HP2 pocket volume in cytosolic (LC3^M-^), membrane-bound apo (LC3^M+R-^) and LIR-bound states (LC3^M+R+^) in three independent 1 μs trajectories.

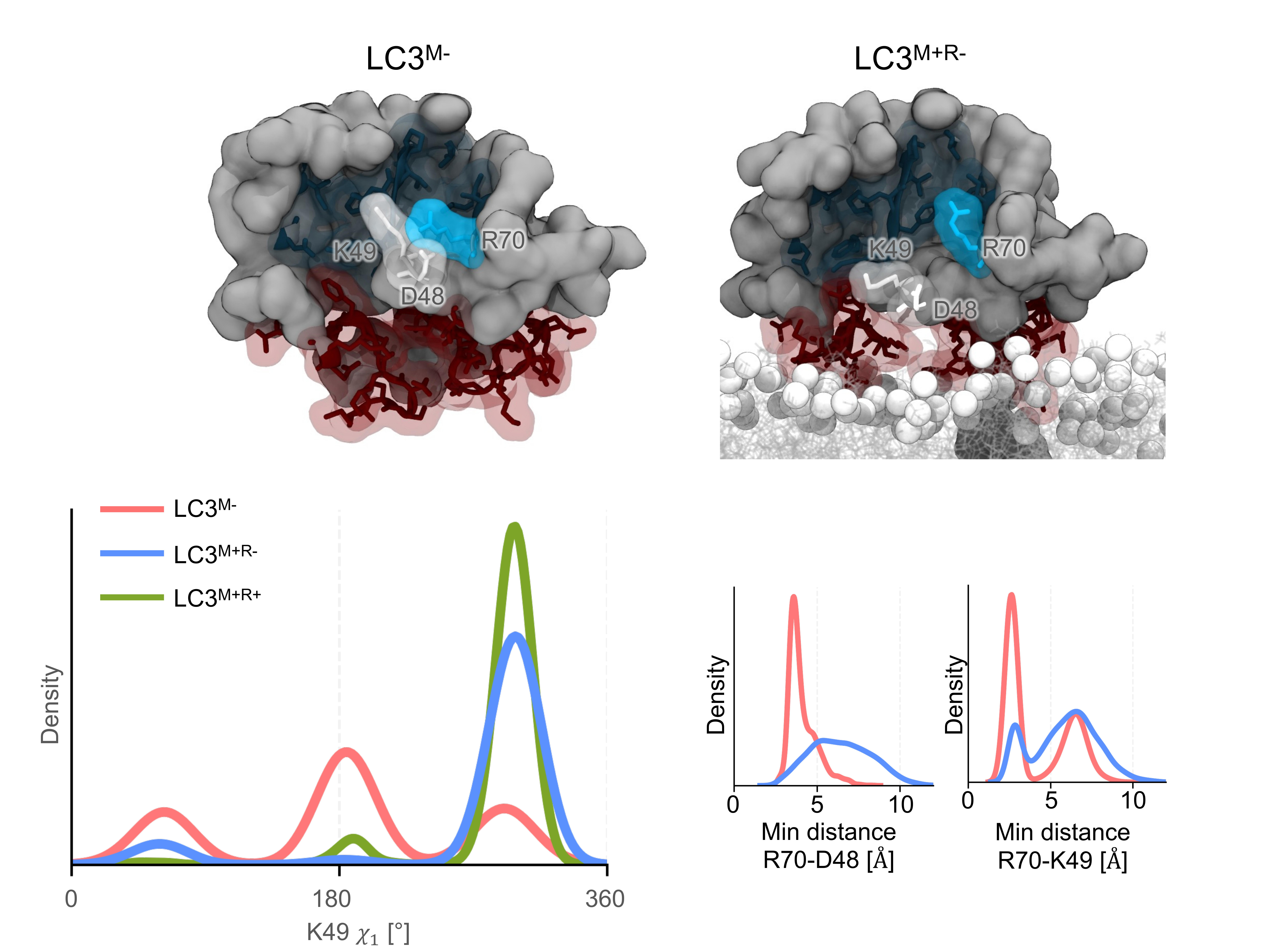

**Supplementary Figure 11. K49 flips out in the presence of the membrane.** Representative snapshots from the simulations are shown for the cytosolic (LC3^M-^) and membrane-bound apo (LC3^M+R-^) states. The relative orientation of K49 with respect to D48 and R70 (HP2 residue) is shown as sticks. The BP and MIS regions are highlighted as a transparent surface. The density plot (bottom left) shows the distribution of the K49 χ_1_ angle for all three LC3 states. The minimum distances of residues K49 and D48 with respect to R70 are shown as density plots (bottom right).

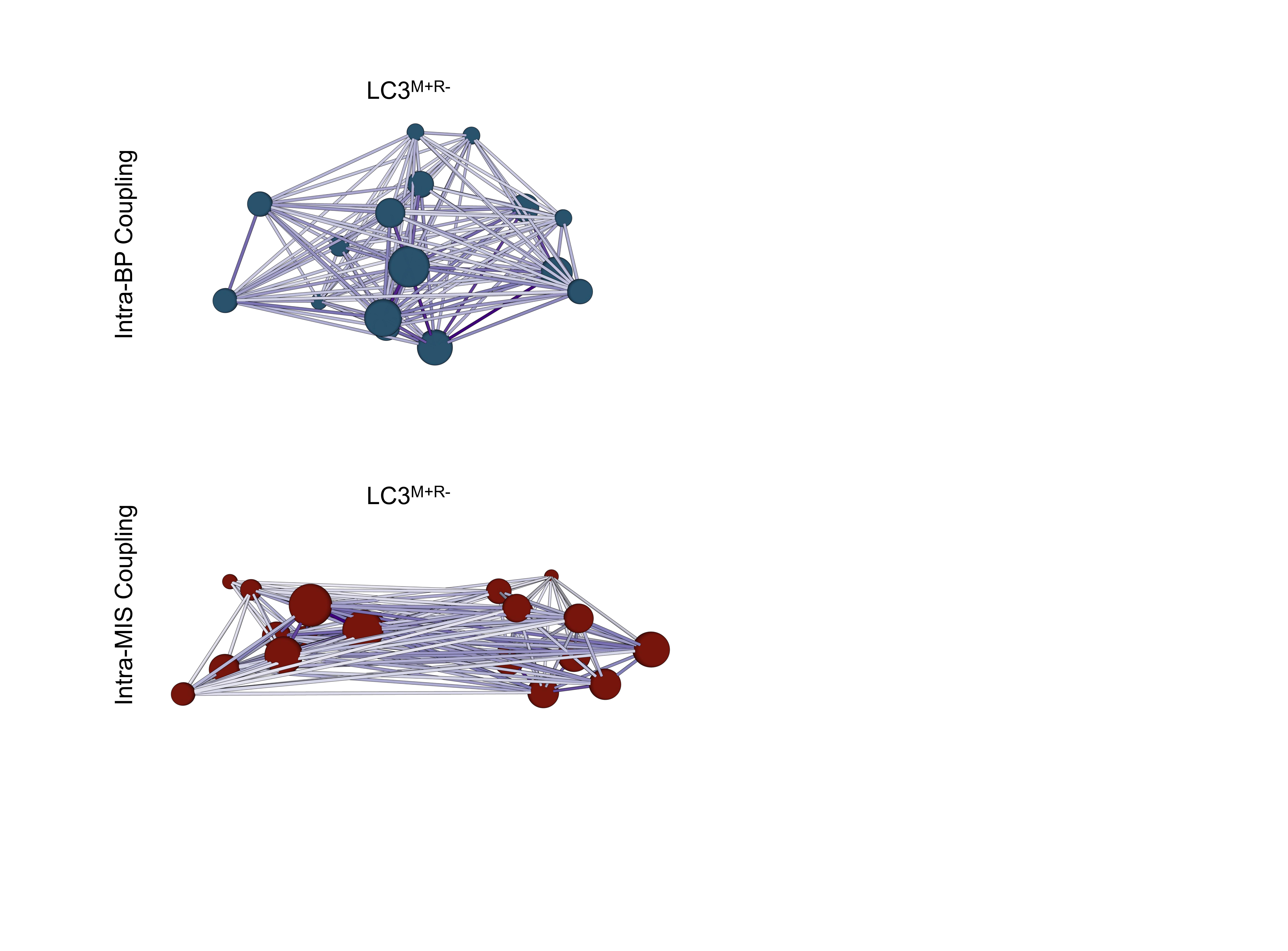

**Supplementary Figure 12: Coupling observed in MIS and BP regions of LC3.** The intra-BP and intra-MIS coupling is shown for the membrane-bound (LC3^M+R-^) state. The network diagram represents the coupling among BP and MIS residues, where each node represents a residue, and the edge color corresponds to the coupling strength (darker the color, the stronger the coupling). The nodes are sized according to the communication strength with the rest of the BP or MIS residues.

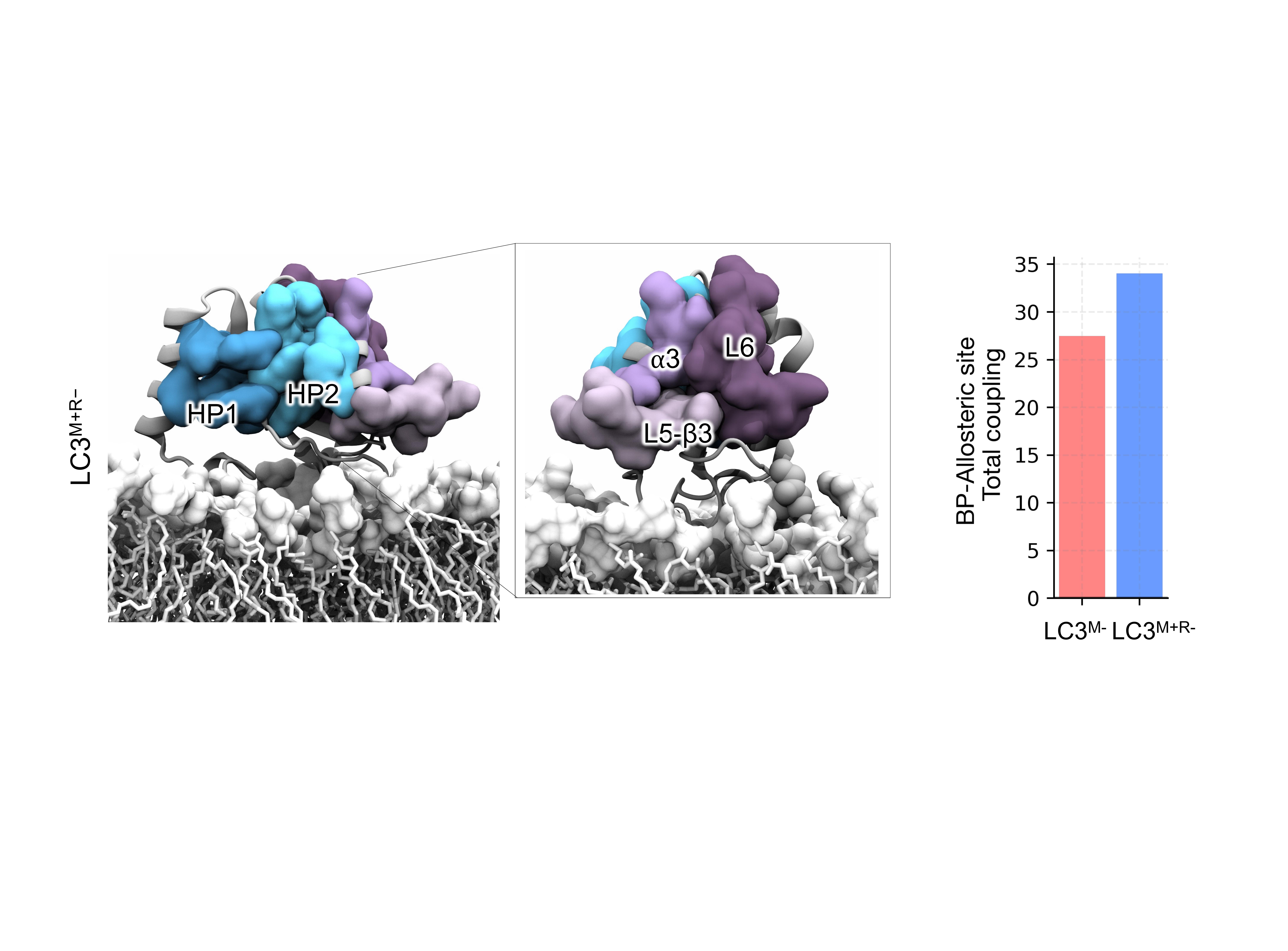

**Supplementary Figure 13. α3-L5-β3-L6 region is positioned next to the HP2 pocket.** (Left) The representative structure snapshot of LC3^M+R-^ is shown. Both front and side regions are visualized. The hydrophobic pockets (HP1, HP2) are visualized in surface representation and colored in shades of blue. α3-L5-β3-L6 region is also visualized as a surface and colored in shades of violet. (Right) The total coupling between allosteric site and binding pocket is plotted for both LC3^M-^ and LC3^M+R-^ states.

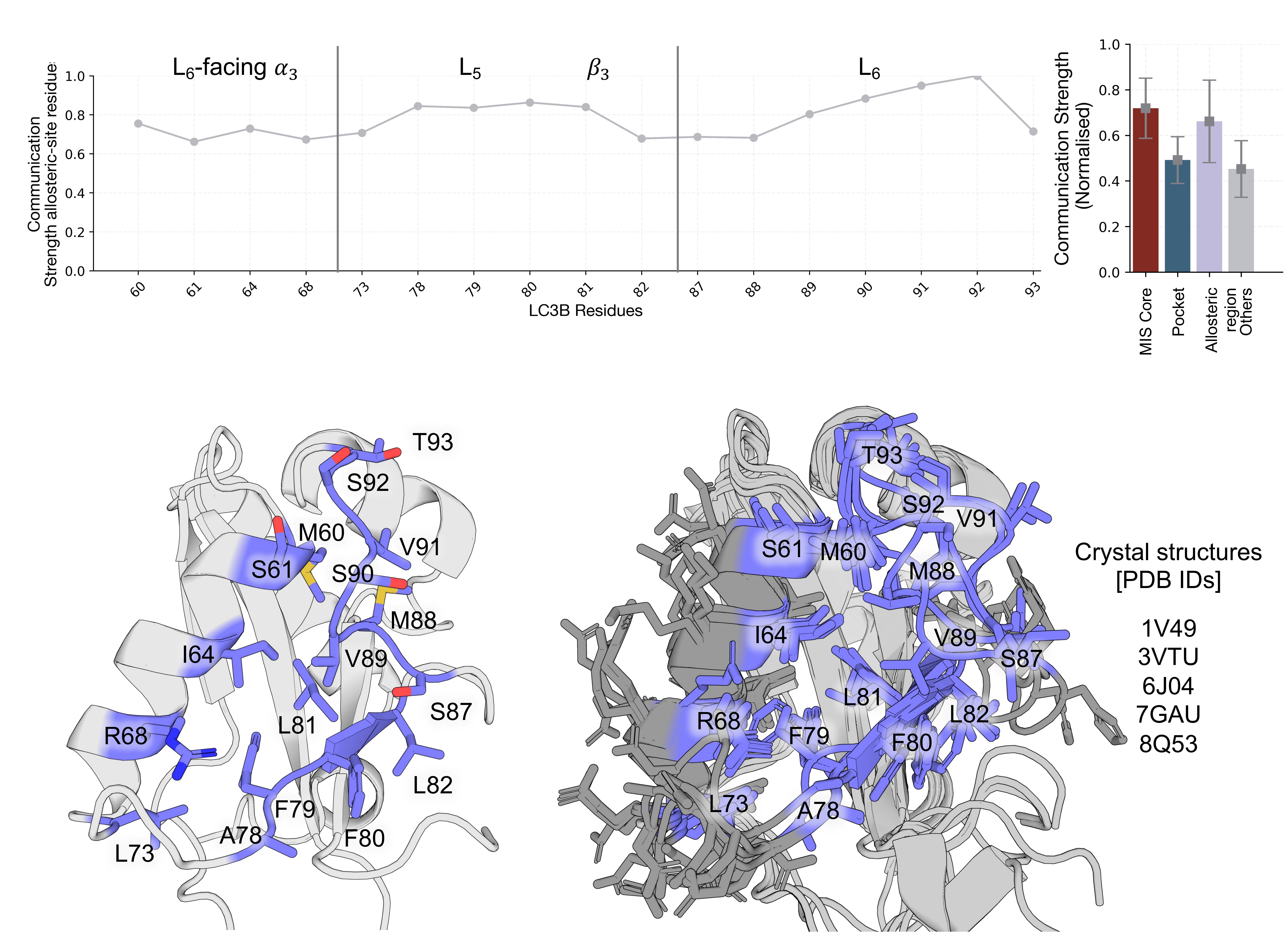

**Supplementary Figure 14. Normalized communication strength in the allosteric site.** Communication strength values are plotted for the residues that constitute the allosteric site of LC3^M+R-^ (greater than the communication strength threshold of 0.65). The corresponding secondary structure elements are highlighted on the plot. (Left) The mean communication strength (observed in LC3^M+R-^) of different regions of LC3 is plotted as a bar graph. The error bar represents the standard deviation. (Bottom left) The residues showing higher strength (>0.65) are mapped onto the representative structure of the LC3^M+R-^ state. (Bottom right) The Human apo crystal structures are superimposed, and α3-L5-β3-L6 region is shown as sticks. The allosteric site residues are colored in slate.

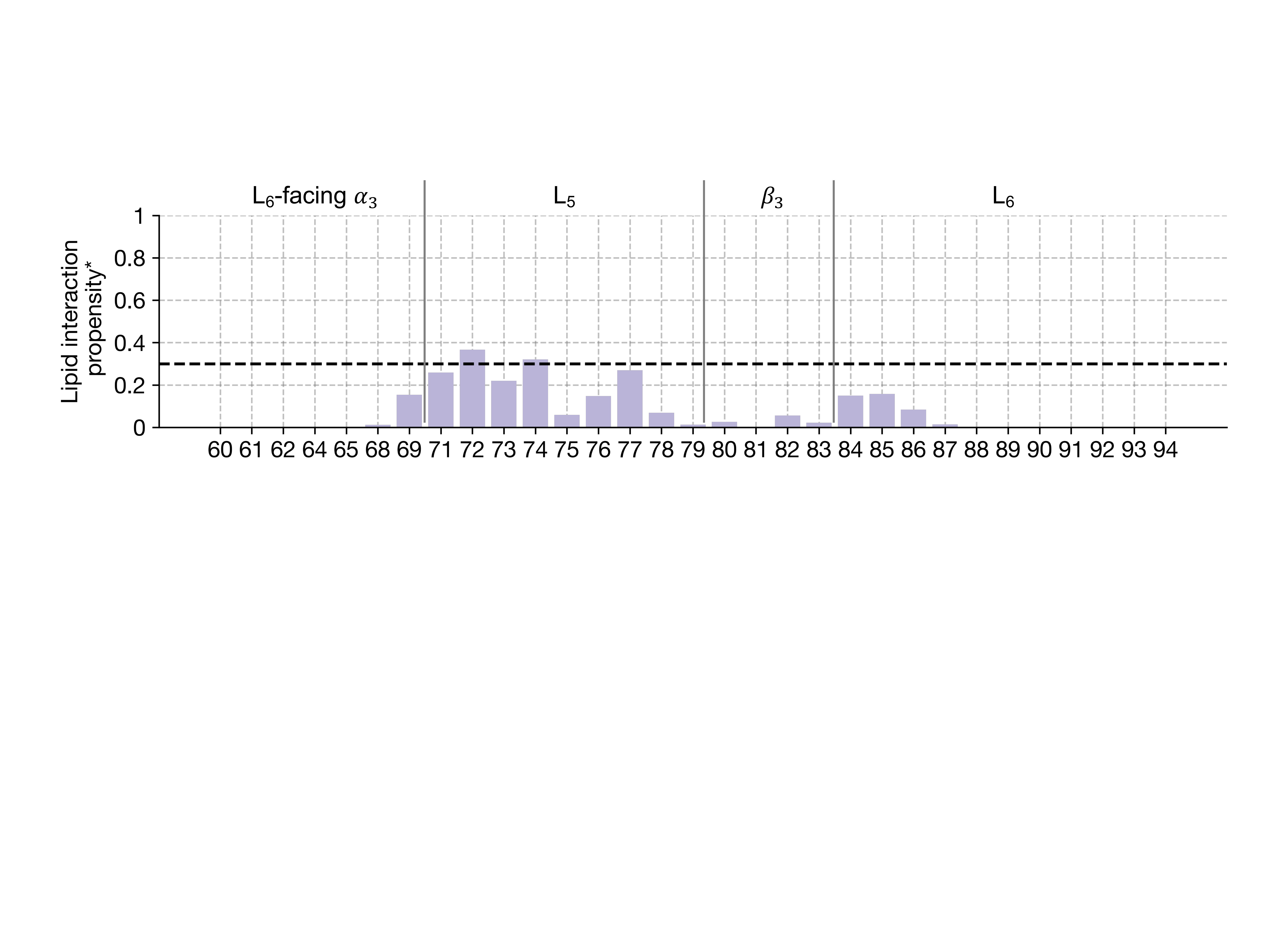

**Supplementary Figure 15. L5 residues of the α3-L5-β3-L6 region** **participate in transient lipid interactions.** The bar plot quantifies the lipid interaction propensity of the α3-L5-β3-L6 region residues (a residue is considered to be in the vicinity of the lipid at a time frame if the minimum distance between a residue and the lipids is less than 5.5 Å). Residues with a propensity greater than 0.3 are considered to face towards the lipids. The propensity is calculated for LC3^M+R-^ state.

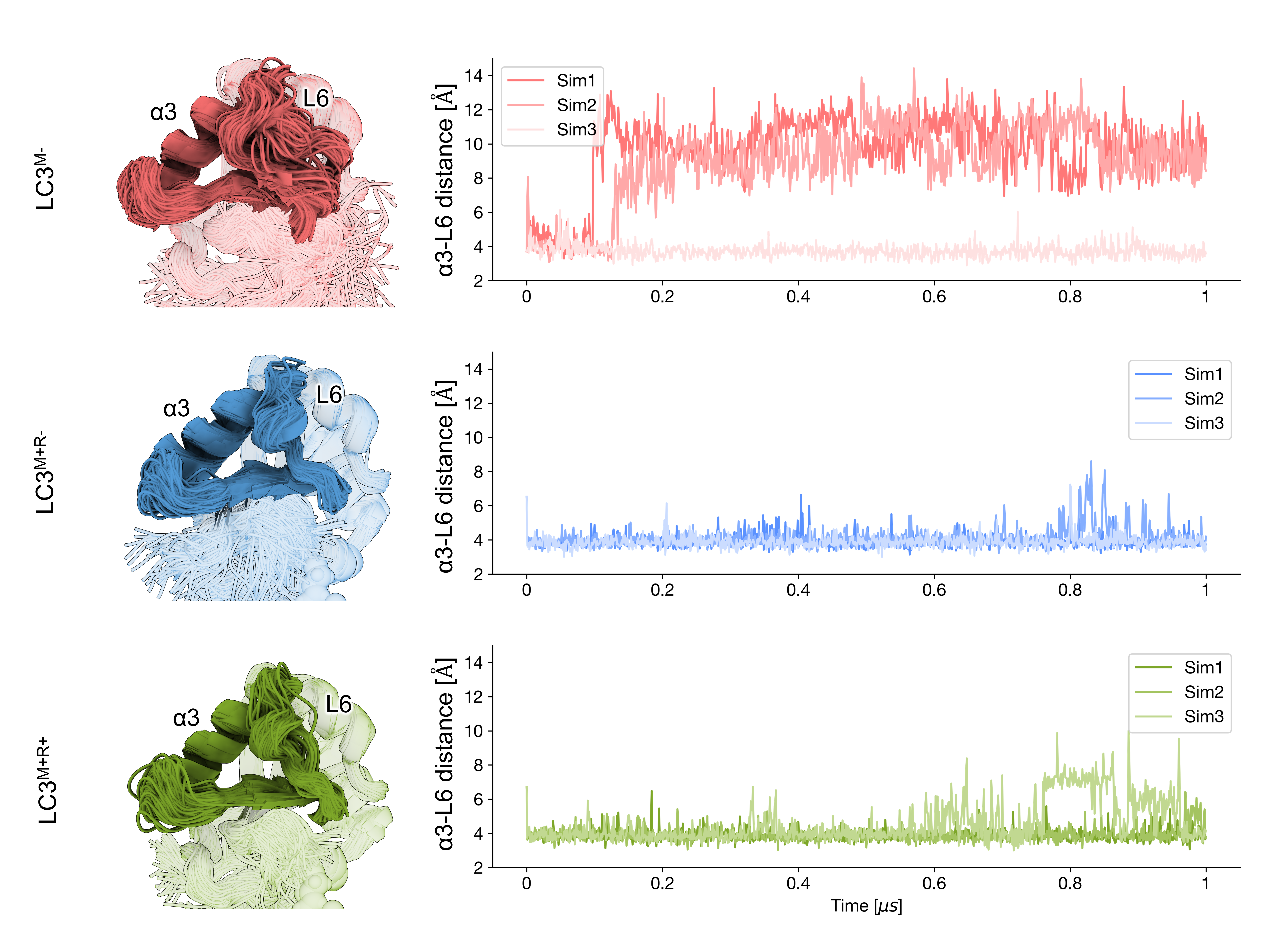

**Supplementary Figure 16: Allosteric site dynamics.** The ensemble of allosteric site conformations is shown for all three states of LC3. The corresponding α3-L6 distances are plotted for all independent simulations of different LC3 states.

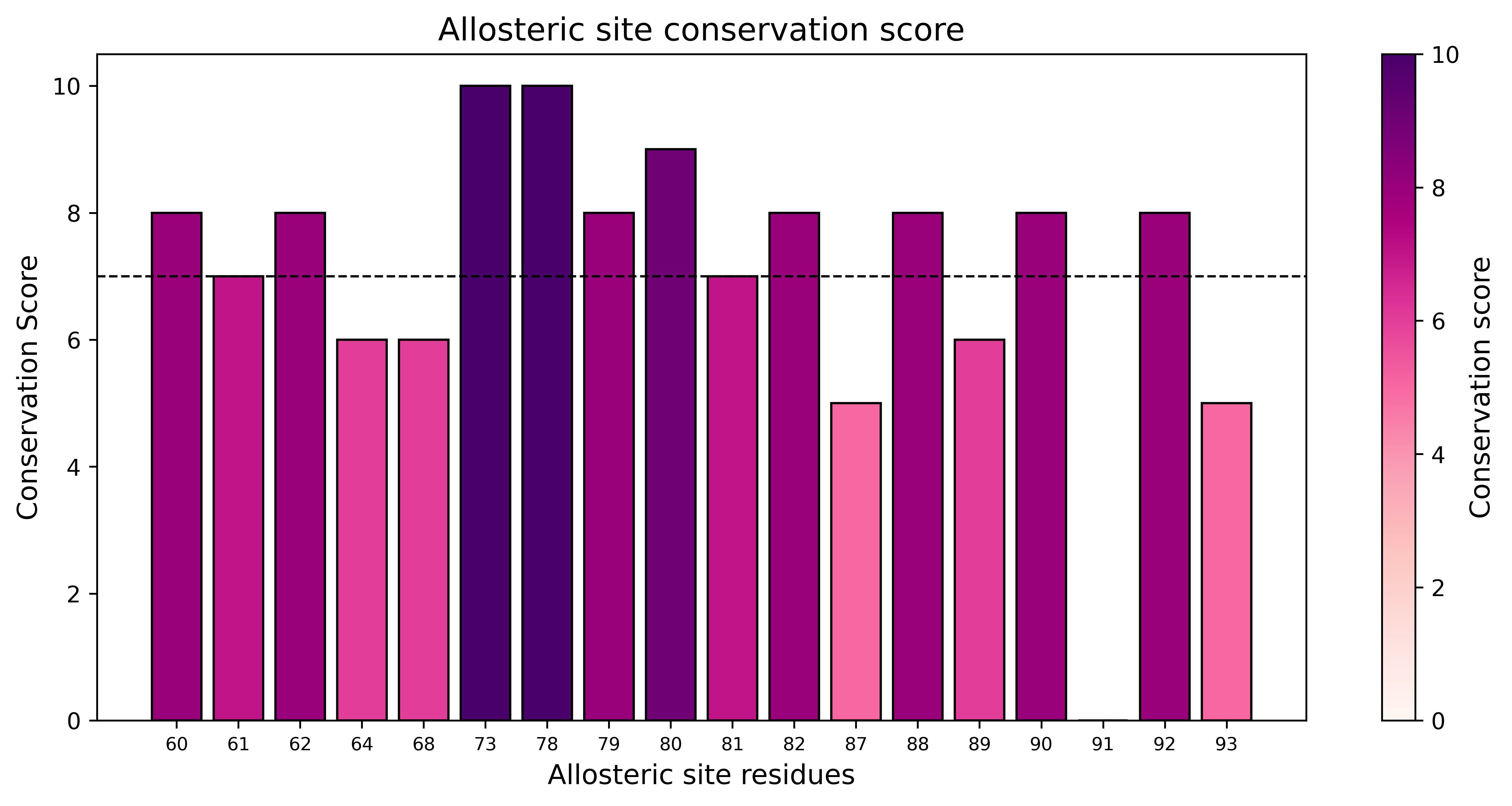

**Supplementary Figure 17: Conservation score of residues in allosteric site.** Conservation score of allosteric site residues encompassing the L6-facing α3 (M60, S61, I64, R68), L5 (L73, A78, F79), β3 (F80, L81, L82) and L6 (S87 to T93). The scores range from 0, least conserved to 10, most conserved.

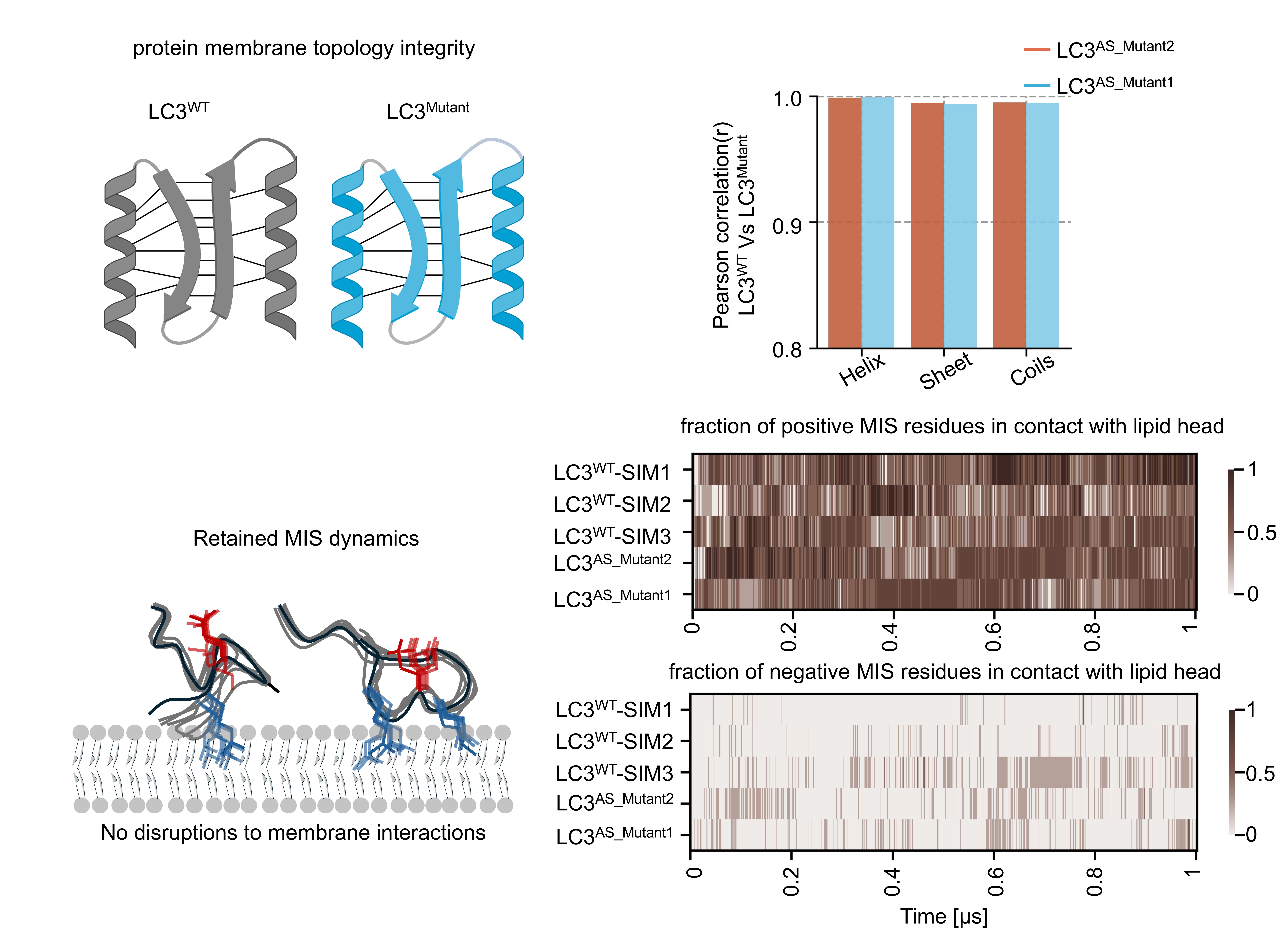

**Supplementary Figure 18: Inactive LC3^AS_Mutant2^ and active LC3^AS_Mutant1^ do not affect the overall structural integrity and retain membrane interaction.** The schematic diagram (top left) highlights the conserved ubiquitin-like fold in LC3^WT^ and mutants in silico. The bar plot (top right) quantifies the Pearson correlation coefficient of secondary structure fractions between mutants (Inactive LC3^AS_Mutant2^ and active LC3^AS_Mutant1^) and LC3^WT^. (bottom left) The diagram illustrates the nature of the LC3-membrane interaction mediated by charged residues. (bottom right) The fraction of positive MIS residues in contact with the lipid head is computed as a function of time and plotted as a heatmap. The contact is defined when the distance between the charged group of the residue and lipid head is less than 5Å. All three replicates of LC3^WT^ are shown along with two LC3 mutants. A similar heatmap is plotted to quantify the fraction of negative MIS residues in contact with the lipid head. Both WT and mutant systems are in the LC3^M+R-^ state**.**

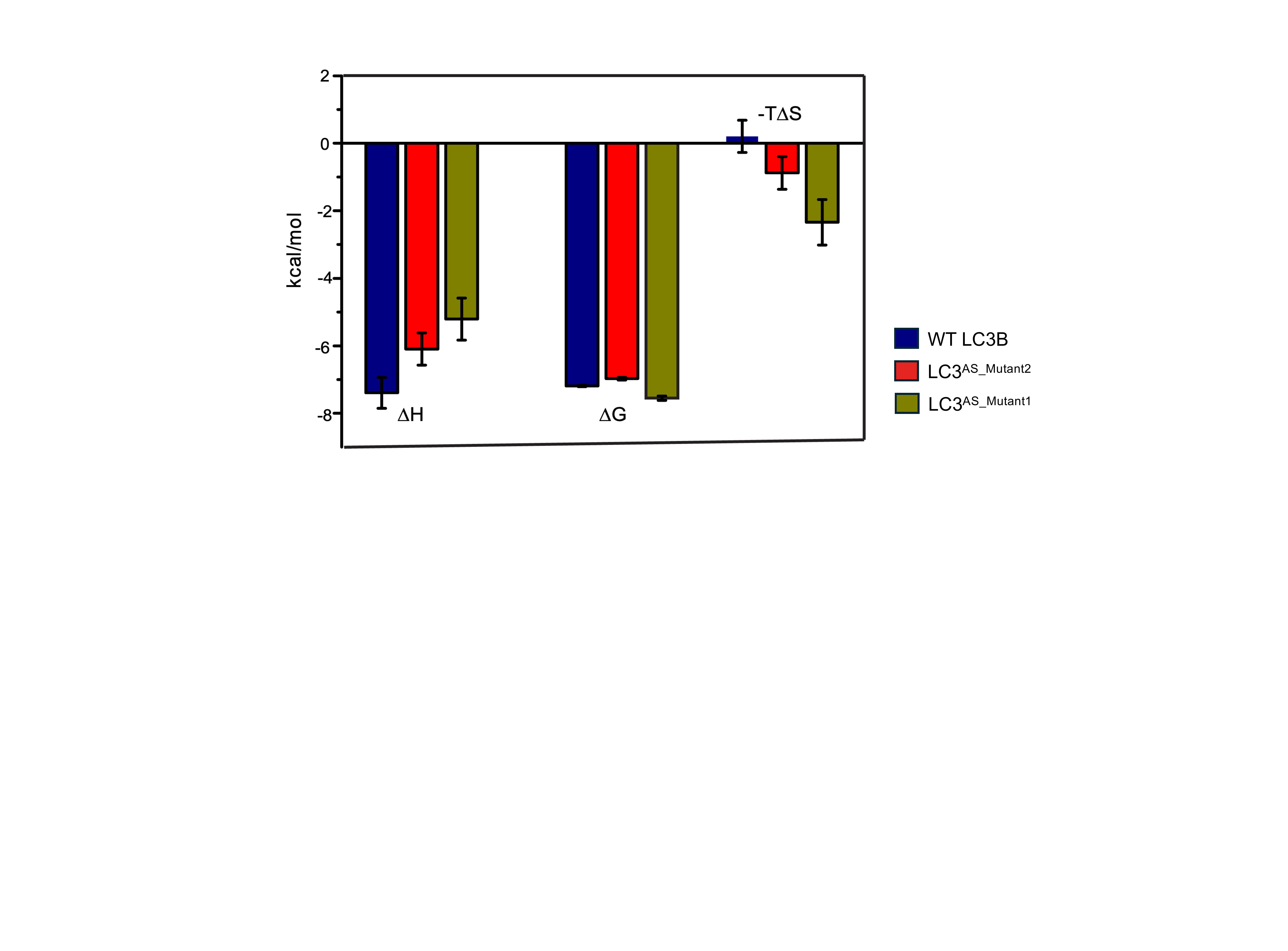

**Supplementary Figure 19. ITC experiments showed active LC3^AS_Mutant1^ enhanced p62 binding.** The bar plot below shows the integrated heat values for LC3^WT^ and LC3^Mutant^ proteins.

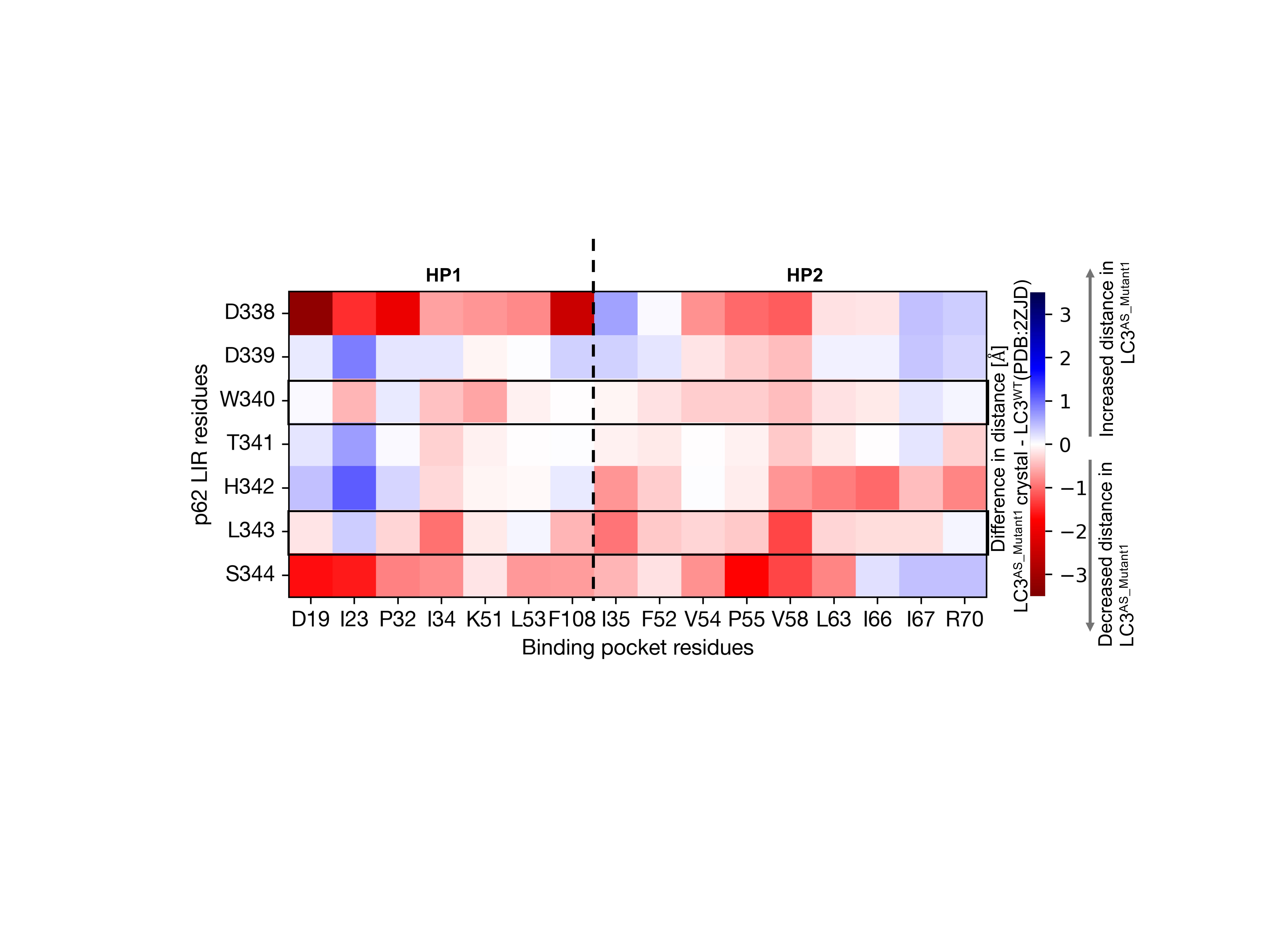

**Supplementary Figure 20. The relative orientation of p62-LIR residues has changed in the active LC3^AS_Mutant1^ crystal structure.** The heatmap quantifies the difference in minimum distance measured between p62-LIR residues and hydrophobic pockets. The negative distances indicate a decrease in residue-residue distance, and the positive values indicate increased residue-residue distances compared to the WT crystal structure (PDB ID: 2ZJD). The rows corresponding to critical LIR residues (W340 and L343) are highlighted.

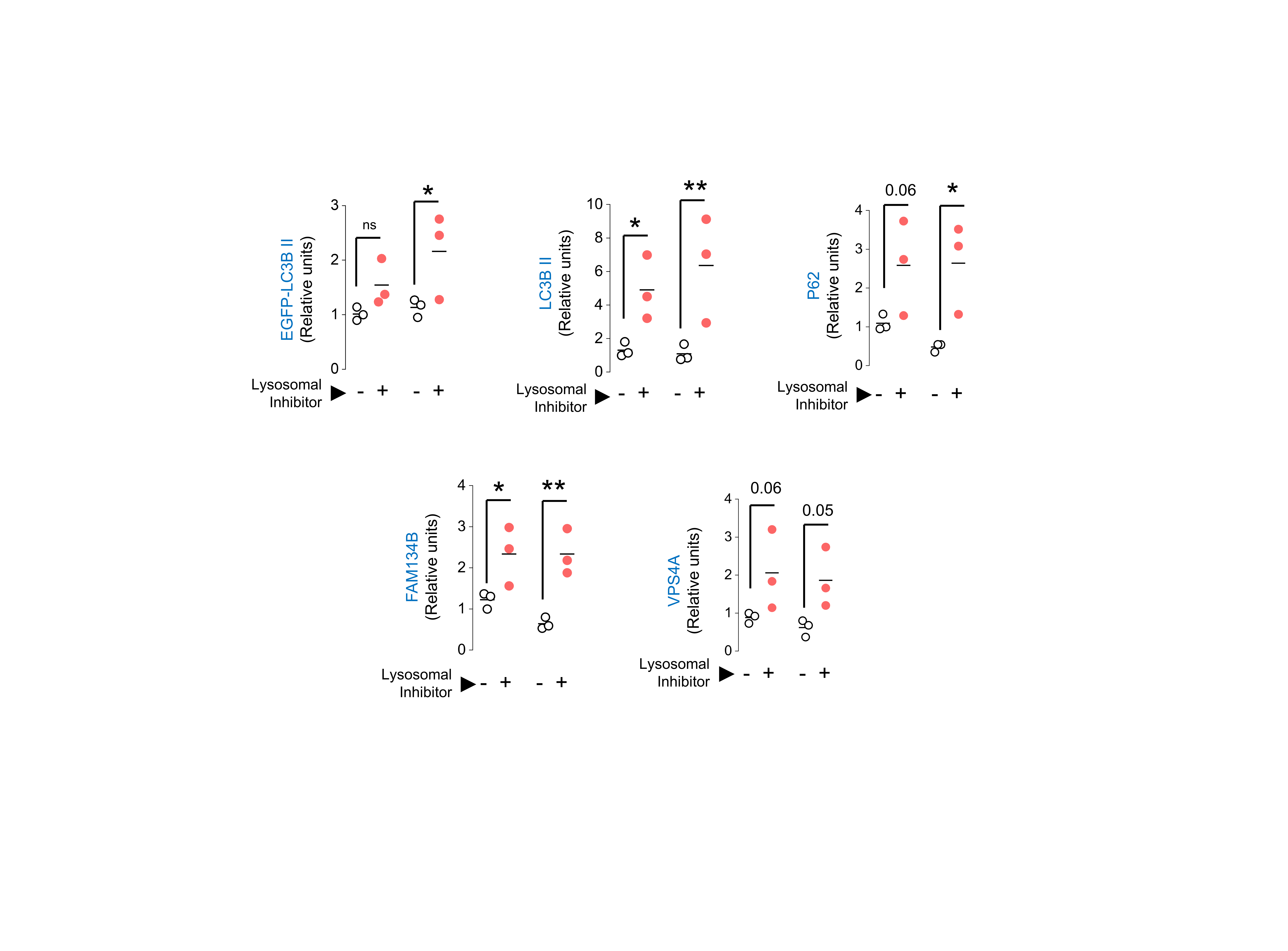

**Supplementary Figure 21. Autophagy flux assays revealed higher cargo turnover in lysosome for active LC3^AS_Mutant1^.** Scatter plots show quantification for the % of lysosomal flux in LC3B^WT^ and active LC3^AS_Mutant1^ for p62, FAM134B, and VPS4A receptors.

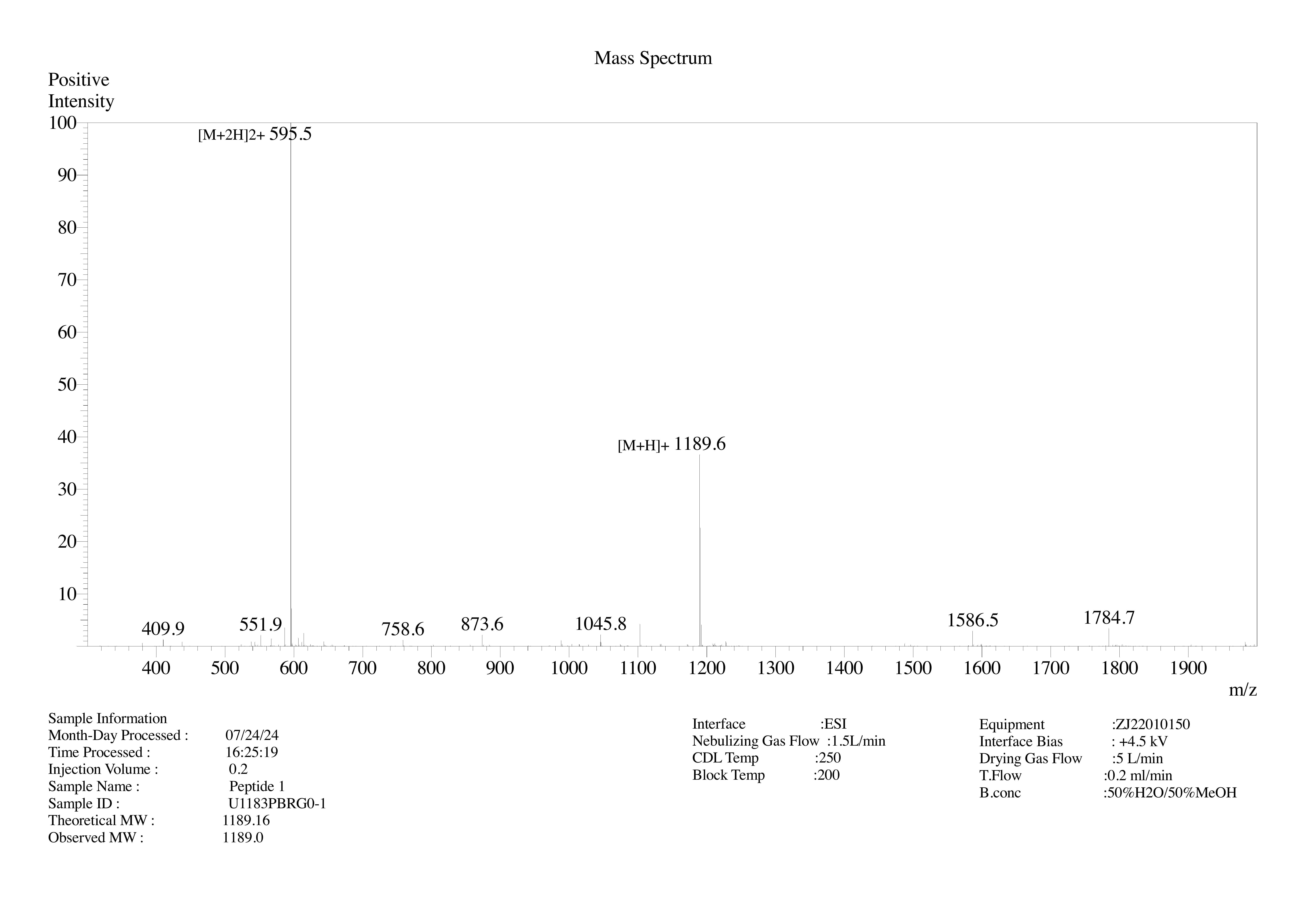

**Supplementary Figure 22. Mass spectrometry analysis of p62-LIR peptide synthesized by Genscript used for ITC and crystallization experiments.**

**Supplementary Videos**

The supplementary videos can be accessed from the figshare link <https://figshare.com/s/f009879f892b56df8e0f>.

**Supplementary Video 1.** The dynamics of HP1, HP2 volumes are shown for LC3^M-^, LC3^M+R-^ and LC3^M+R+^ states. The pocket volume is visualized as *QuickSurf,* and the residues of the HP1 and HP2 are shown as sticks. The protein backbone is shown in cartoon representation. The movie is recorded with the smoothening over the running window of three consecutive frames at 30 frames per second. Each frame in the movie corresponds to 1ns in the MD simulation trajectory.

**Supplementary Video 2.** The orientation of the allosteric site elements (at the side area of LC3) is shown as observed in MD simulation trajectories for the three LC3 states. The allosteric site elements (α3, L5, β3, and L6) are colored with different shades of violet and are visualized as *QuickSurf*. The protein backbone is shown in cartoon representation. Here, each frame represents 1ns. The movie is rendered at 30 frames per second and is smoothened over the running window of three consecutive frames.

**Supplementary Video 3.** The dynamics of the allosteric site are shown for inactive LC3^AS_Mutant2^ and active LC3^AS_Mutant1^ *in-silico* mutants, respectively. The elements of the allosteric site are visualized as *QuickSurf.* The protein backbone is shown in cartoon representation. The mutated positions at the allosteric site are colored red. The movie is smoothened over the running window of three frames and saved at 30 frames per second.

**Supplementary Table 1: Unique contacts observed in cytosolic LC3B (LC3^M-^), and Membrane-bound LC3 apo (LC3^M+^) states.**

| **LC3B Regions** | **Cytosolic LC3B (LC3^M-^)** | **Membrane-bound LC3 apo (LC3^M+^)** |
| --- | --- | --- |
| MIS-Rest | 6 | 2 |
| Pocket-Rest | 4 | 3 |
| Intra MIS | 12 | 3 |
| Intra Pocket | 2 | 2 |
| Intra Rest | 7 | 15 |

**Supplementary Table 2: Thermodynamic parameters for LC3^WT^, and mutants binding to p62 peptide.**

|  | **n** | **K_d_ (µM)** | **∆H (kcal/mol)** | **∆G (kcal/mol)** | **-T∆S (kcal/mol)** |
| --- | --- | --- | --- | --- | --- |
| LC3^WT^ | 1.65 ±0.08 | 5.42 ± 0.19 | -7.39 ± 0.45 | -7.19 ± 0.02 | 0.20 ± 0.48 |
| LC3^AS_Mutant1^ | 1.21 ±0.05 | 2.92 ± 0.3 | -5.21 ± 0.62 | -7.55 ± 0.06 | -2.34 ± 0.67 |
| LC3^AS_Mutant2^ | 0.98 ±0.04 | 7.72 ± 0.5 | -6.09 ± 0.47 | -6.98 ± 0.04 | -0.88 ± 0.48 |

**Supplementary Table 3. MD simulation system information for three LC3 states**

| **System** | **System type** | **Box dimension** | **No of replica** | **Total Simulation time per replica [in µs]** | **Cumulative time**  **[in µs]** |
| --- | --- | --- | --- | --- | --- |
| Cytosolic LC3 | LC3^M-^ | 115.3 Å x 115.3 Å x 140.0 Å | 3 | 1 | 3 |
| Membrane-bound LC3 (ER-like membrane) | LC3^M+R-^ | 115.3 Å x 115.3 Å x 140.0 Å | 3 | 1 | 3 |
| Membrane-bound LC3 with p62-LIR | LC3^M+R+^ | 115.3 Å x 115.3 Å x 140.0 Å | 3 | 1 | 3 |
| **Cumulative simulation time** | | | | | **9 µs** |

**Supplementary Table 4. MD simulation system for LC3 mutant candidates**

| **Mutant candidates** | **Potentially**  **active/inactive** | **System type** | **Box dimension** | **Total Simulation time [in µs]** |
| --- | --- | --- | --- | --- |
| A75K | Active | LC3^M+R-^ | 115.3 Å x 115.3 Å x 140.0 Å | 1 |
| I64D_V89P_V91D | Active | LC3^M+R-^ | 115.3 Å x 115.3 Å x 140.0 Å | 1 |
| S90A | Active | LC3^M+R-^ | 115.3 Å x 115.3 Å x 140.0 Å | 1 |
| N76A_F79A | Active | LC3^M+R-^ | 115.3 Å x 115.3 Å x 140.0 Å | 1 |
| M60A | Active | LC3^M+R-^ | 115.3 Å x 115.3 Å x 140.0 Å | 1 |
| L81M | Active | LC3^M+R-^ | 115.3 Å x 115.3 Å x 140.0 Å | 1 |
| A75D | Active | LC3^M+R-^ | 115.3 Å x 115.3 Å x 140.0 Å | 1 |
| I64K_V89D_V91F | Inactive | LC3^M+R-^ | 115.3 Å x 115.3 Å x 140.0 Å | 1 |
| S61K_S90D | Inactive | LC3^M+R-^ | 115.3 Å x 115.3 Å x 140.0 Å | 1 |
| A75D_N76E_F79D | Inactive | LC3^M+R-^ | 115.3 Å x 115.3 Å x 140.0 Å | 1 |
| I64F | Inactive | LC3^M+R-^ | 115.3 Å x 115.3 Å x 140.0 Å | 1 |
| M60K_M88D | Inactive | LC3^M+R-^ | 115.3 Å x 115.3 Å x 140.0 Å | 1 |
| I64T_L81M | Inactive | LC3^M+R-^ | 115.3 Å x 115.3 Å x 140.0 Å | 1 |
| **Cumulative simulation time** | | | | **13 µs** |
